# Intracellular Trafficking SNARE Protein, Syntaxin-6, is a Modifier of Prion and Tau Pathogenesis *in vivo* and in Cellular Models

**DOI:** 10.1101/2025.02.19.635172

**Authors:** Elizabeth Hill, Mitali Patel, Juan M Ribes, Jacqueline Linehan, Fuquan Zhang, Michael Farmer, Tatiana Jakubcova, Shyma Hamdan, Andrew Tomlinson, Tiziana Ercolani, Christian Schmidt, Parvin Ahmed, George Thirlway, Silvia Purro, Fabio Argentina, Aline T Marinho, Emma Jones, Nicholas Kaye, Craig Fitzhugh, Rohan de Silva, Graham S Jackson, Sebastian Brandner, Peter Kloehn, John Collinge, Thomas J Cunningham, Simon Mead

## Abstract

Syntaxin-6, a SNARE protein involved in intracellular protein trafficking, is a proposed risk factor for sporadic prion disease, progressive supranuclear palsy and Alzheimer’s disease. However, no study has validated its functional role in these diseases, explored the disease stage at which it is acting nor its mechanism of action. Here, we show that syntaxin-6 acts at early stages of prion disease in experimental mice by increasing disease transmission risk following inoculation with low prion doses. Conversely, syntaxin-6 does not affect prion propagation kinetics or toxicity during established disease. Syntaxin-6 manipulation in cellular models profoundly alters the subcellular distribution and morphologies of disease-related PrP and modifies prion export. Furthermore, syntaxin-6 knockout in a transgenic tauopathy mouse model exerts protective effects on numerous physiological, behavioural and neuropathological outcome measures. Therefore, our studies firmly establish syntaxin-6 as a modifier of prion and tau pathogenesis, providing key insights into a fundamental mechanism of neurodegeneration.

## Introduction

Prion diseases are fatal, infectious neurodegenerative diseases, characterised by recruitment of the cellular prion protein (PrP^C^) into fibrillar, multimeric amyloid assemblies^1^. Sporadic Creutzfeldt-Jakob disease (sCJD), the most common human prion disease, is thought to occur when seeds of misfolded PrP form spontaneously and then propagate by fibril growth and fission. Collective evidence over decades provides compelling evidence that other neurodegenerative disorders incorporate so-called “prion-like” mechanisms^1,2^ including evidence for human-human transmission of Aβ pathology driving cerebral amyloid angiopathy and Alzheimer’s disease (AD)^3,4^. Furthermore, in the tauopathies, tau protein misfolds into distinct pathological assemblies, which seed further tau misfolding and spread intercellularly, while faithfully maintaining phenotypic properties, reminiscent of prion strains^5–9^. As there are fundamental similarities between prion disease and tauopathy pathogenesis, it is reasonable to hypothesise that there may be shared molecular modifiers of these processes.

Human genetics studies offer a valuable approach to identify molecular players implicitly causal in disease pathogenesis. Variants at the syntaxin-6 (*STX6)* locus have been identified by genome-wide association studies (GWAS) as shared genetic risk factors for sCJD^10^ and the most common primary tauopathy, progressive supranuclear palsy (PSP)^11–15^. Furthermore, a recent proteome-wide association study (PWAS) found a causal association between STX6 protein levels and AD^16^. These studies propose syntaxin-6 as a candidate prion/prion-like modifier, exerting pleiotropic risk effects across multiple neurodegenerative diseases. However, genetic data only provide suggestive evidence for the most likely gene driving disease risk at a locus. Although increased *STX6* expression is linked to disease risk by transcriptomic and proteomic analyses^15–18^, providing a causal genetic mechanism, functional studies are necessary to validate this mechanism and translate these genetic discoveries into more precise pathogenic disease mechanisms.

Syntaxin-6 encodes an intracellular trafficking protein^19,20^, which functions in a highly cell-type specific manner^21^. Recent work suggests that *STX6* risk variant driven expression changes are strongest in oligodendrocytes^15,17,18,22^, supporting its study in a multicellular system to capture non-cell autonomous effects. Facilitating this, we generated *Stx6^-/-^* mice in collaboration with MRC Harwell, which are viable with no gross neurological, physiological or behavioural phenotypes^23^. However, prion infection of *Stx6^-/-^*mice only had very modest effects on the endpoint clinical outcome measures providing no clear evidence of a modifying role^23^.

Prion disease proceeds in two distinct mechanistic phases in prion-inoculated mice^24,25^. The clinically and neuropathologically silent first phase is characterised by an exponential increase in prion titre. This is followed by a second phase, after prion titres have plateaued, where neurotoxicity and neuropathological features become established, culminating in the onset of clinical disease. It is possible that there is a similar progression of events in sCJD, although here disease is initiated by spontaneous prion formation as a rare stochastic event and subsequent spread throughout the central nervous system. We hypothesised that syntaxin-6 confers risk of prion disease by modification of one of these key three stages: the establishment of disease, prion propagation or prion-induced toxicity. This hypothesis may be extended to tau pathogenesis given that tau seeding kinetics have similar features to the two-phase kinetics model in a tauopathy mouse model, with histopathological markers emerging later^26^, along with evidence for a dissociation between tau propagation and tau-induced toxicity^27–33^.

The main aims of this work were to functionally validate a role for syntaxin-6 in prion and tau pathogenesis *in vivo,* explore the disease stage at which syntaxin-6 is acting and inform on causal pathogenic disease mechanisms. To achieve this, we conducted prion transmission studies in *Stx6^+/+^* and *Stx6^-/-^* mice to evaluate differences in the establishment of prion infection, prion propagation and neurotoxicity. To extend this to tau pathogenesis, we crossed *Stx6^-/-^* mice with a widely used P301S transgenic tauopathy mouse model^34^ to assess alterations in disease progression. These studies propose syntaxin-6 as a modifier of prion and tau pathogenesis *in vivo,* modulating early stages of the disease. Furthermore, our results support altered trafficking and export of prions by syntaxin-6 as the plausible molecular susceptibility mechanism. Thus, this work informs on a fundamental mechanism of neurodegeneration, providing important, novel insights into the role of a pleiotropic prion/prion-like modifier, grounded in human genetics evidence.

## Results

### Syntaxin-6 modifies prion pathogenesis *in vivo* by modulating the risk of disease development

The “gold standard” paradigm for studying prion disease pathogenesis is prion transmission in mice. Mice are naturally susceptible to prion infection, developing *bona fide* disease with faithful recapitulation of the clinical and neuropathological hallmarks of human disease when experimentally inoculated with prions such as the mouse-adapted scrapie prion strain, RML^35^. To determine whether *Stx6* knockout modified the risk of mice developing prion disease, we intracerebrally infected *Stx6^+/+^* and *Stx6^-/-^* mice (n=90/genotype) with a 10-fold serial dilution series of 10% (w/v) RML prion-infected brain homogenate. The focus of this study was to examine prion doses with a partial attack rate (defined as <90% across both arms of the study: 10^-5^, 10^-6^, 10^-7^, 10^-8^), where the likelihood of disease development was uncertain, providing a paradigm to assess whether syntaxin-6 modulated that risk.

For each of the 10^-5^, 10^-6^, 10^-7^ and 10^-8^ concentrations, the proportion of prion disease cases relative to the animals surviving to the end of the study (attack rate, see methods) was consistently lower in *Stx6^-^*^/-^ mice relative to *Stx6^+/+^* mice, suggesting they were more resistant to disease development (**Table 1**; **Fig. 1**). Logistic regression analysis demonstrated a significant effect of dose (P < 0.0001) and genotype (P = 0.05) on prion disease diagnosis, with infected *Stx6^+/+^*mice having 2.19 [95% CI: 1.01-4.56] times higher odds of developing prion disease compared to *Stx6^-/-^* animals at concentrations 10^-5^ and lower. This suggests syntaxin-6 knockout reduces susceptibility to prion infection, which is further supported by calculations estimating the “effective” dose administered to *Stx6^+/+^* and *Stx6^-/-^* mice, which was reduced in *Stx6^-/-^* mice by a log order of magnitude relative to *Stx6^+/+^* mice (10^5.82^ LD_50_/mL and 10^6.61^ LD_50_/mL respectively).

**Figure 1.**
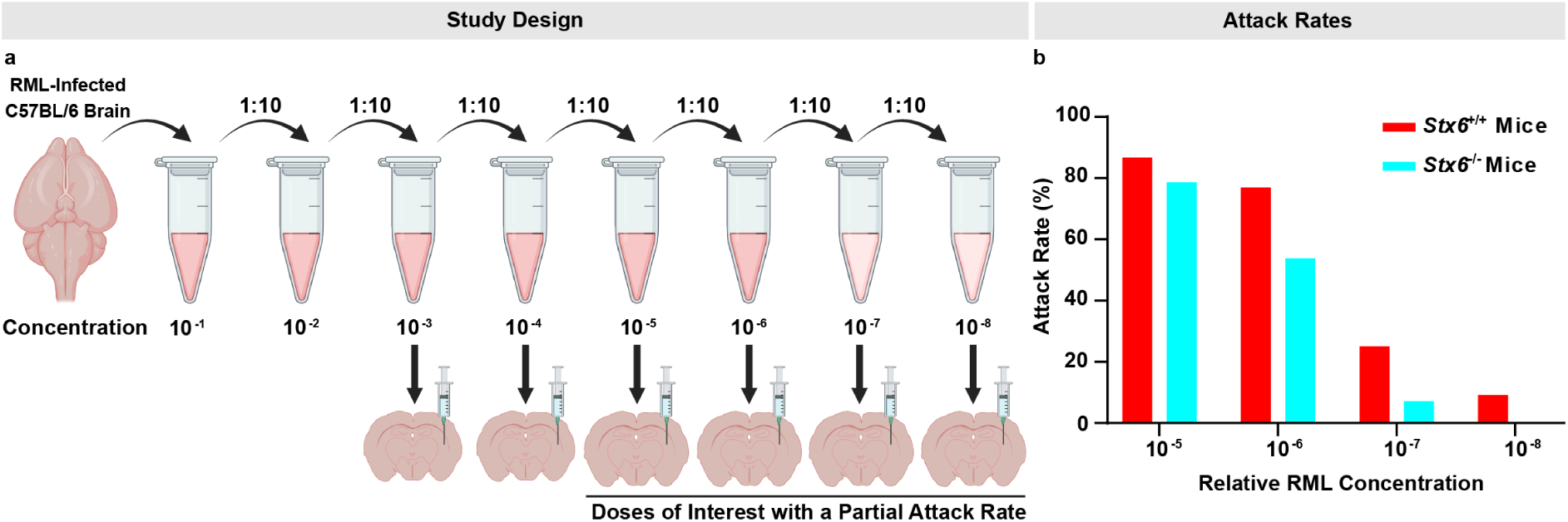
Syntaxin-6 Knockout Renders Mice Less Susceptible to Prion Disease When Administered Low Doses of Prions. **(a)** Experimental design of intracerebral inoculation of *Stx6*^+/+^ and *Stx6*^-/-^ mice with a 10-fold dilution series of 10% (w/v) RML prion-infected brain homogenate (n=15/genotype/dose) with prion doses with a partial attack rate (<90%) being of interest. Created in BioRender. One, S. (2025). https://BioRender.com/t36g169. **(b)** Bar chart showing the attack rates of prion disease in mice administered 10^-5^, 10^-6^, 10^-7^ or 10^-8^ concentrations of RML prions.

**Table 1.** Assessment of Susceptibility Differences in *Stx6^+/+^* and *Stx6^-/-^*Mice Inoculated with RML Prion Dilutions with a Partial Attack Rate. Attack rates of *Stx6^+/+^* and *Stx6^-/-^* mice infected with dilutions of 10% (w/v) RML prion-infected brain homogenate with a partial attack rate (defined as <90% across both arms of the study). Attack rate was defined as the total of affected mice as a proportion of the number of inoculated mice, excluding mice which died due to unrelated health concerns or animals where there was a discrepancy between the clinical and pathological observations. Calculated odds ratios are shown with 95% confidence intervals (CI) determined by the Baptista-Pike method^82^. The p-value refers to the results of logistic regression analysis with prion dose and genotype as factors.

| Concentration | *Stx6*<sup>+/+</sup> Mice |  | *Stx6*<sup>-/-</sup> Mice |  | Odds Ratio (95% CI) | P-Value (Genotype) |
| --- | --- | --- | --- | --- | --- | --- |
|  | Attack Rate | Attack Rate (%) | Attack Rate | Attack Rate (%) |  |  |
| 10<sup>-5</sup> | 13/15 | 86.7 | 11/14 | 78.6 | 1.77 (0.309-11.2) | **0.0500** |
| 10<sup>-6</sup> | 10/13 | 76.9 | 7/13 | 53.8 | 2.86 (0.509-12.7) |  |
| 10<sup>-7</sup> | 3/12 | 25.0 | 1/14 | 7.14 | 4.33 (0.537-59.9) |  |
| 10<sup>-8</sup> | 1/11 | 9.09 | 0/15 | 0.00 | Infinity (0.152-infinity) |  |
| Combined | 27/51 | 52.9 | 19/56 | 33.9 | **2.19 (1.01-4.56)** |  |

### Syntaxin-6 does not alter prion propagation or prion-induced neurotoxicity *in vivo* during established disease

This modifying effect of syntaxin-6 on disease susceptibility *in vivo* could either be acting through directly modulating the establishment of prion infection, or alternatively, by altering subsequent prion propagation or prion-induced neurotoxicity. To systematically interrogate a role for syntaxin-6 in prion replication and neurotoxicity, we infected *Stx6^+/+^* and *Stx6^-/-^* mice (n=110/genotype) with 1% (w/v) RML prion-infected brain homogenate, with animals subsequently being culled at multiple predefined time points or at the onset of clinical disease (**Fig. 2a**). This allowed age-matched, cross-sectional analyses of prion-related phenotypes in the evolving stages of disease, with timed culls of PBS-inoculated mice providing the negative control (**Fig. 2b**).

**Figure 2.**
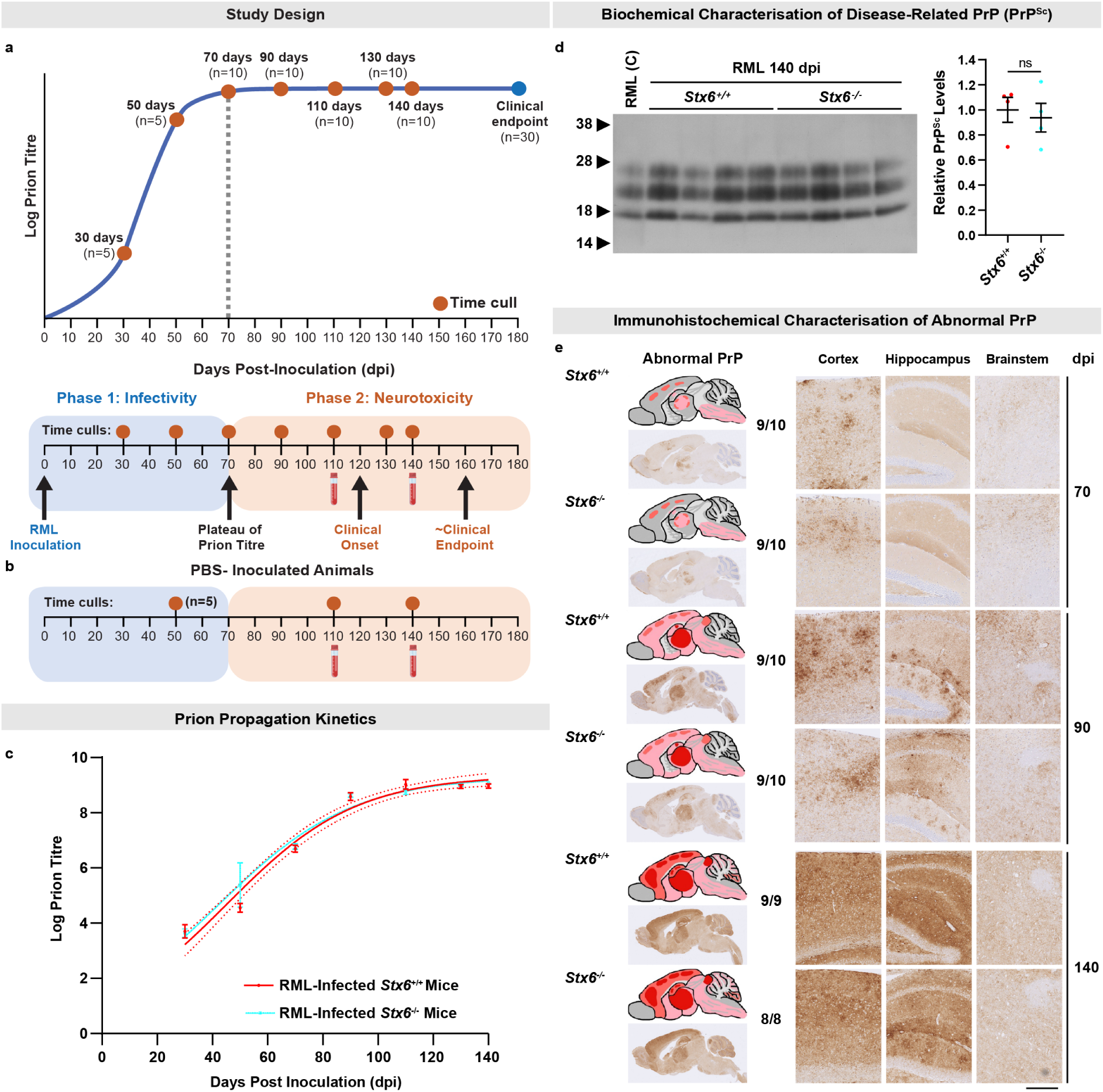
Syntaxin-6 Knockout has no Effect on Prion Propagation Kinetics or Levels of Disease-Related PrP in Mice Infected with RML Prions. **(a)** Experimental design mapped onto the two-phase kinetics model whereby RML-infected *Stx6^+/+^* and *Stx6^-/-^* mice were culled at predefined time points to assess for differences in prion propagation and neurotoxicity-related outcome measures. **(b)** Timed culls of PBS-inoculated controls. **(c)** Prion titres (log tissue-culture infectious units (TCIU) per gram brain) of RML-infected *Stx6*^+/+^ and *Stx6*^-/-^ mice (n=5-10/genotype/time point), across the incubation period (dpi, days post inoculation). All PBS-inoculated brain homogenates were negative for infectivity (not shown). Curves were fitted using the logistic growth model (goodness of fit, r^2^: 0.928 (*Stx6^+/+^*), 0.926 (*Stx6^-/-^)).* Bars indicate mean ± SEM with dotted lines representing 95% confidence intervals. **(d)** 10% (w/v) brain homogenates from RML-infected *Stx6^+/+^* and *Stx6^-/-^* mice at 140 dpi were analysed by immunoblotting with the anti-PrP antibody, ICSM35, after digestion with proteinase K (50 µg/mL, 37°C, 1 h). Semi-quantification of the PrP^Sc^ signal was performed using densitometry with each sample being normalised to the average of RML-infected *Stx6^+/+^* mice. Bar graphs represent mean ± SEM of 4 biological replicates/genotype. C, positive control RML sample. Statistical differences were tested with a Student’s t-test (P = 0.693). **(e)** Spatiotemporal differences of PrP deposition were assessed by immunohistochemistry using the anti-PrP antibody, ICSM35, at 70-, 90- and 140-days post inoculation (dpi) (n=9-10/genotype). A schematic is shown to represent the overall staining pattern in the respective groups (pale pink: mild PrP deposition; pink shading: moderate PrP deposition; red shading: intense PrP deposition). Representative images of the whole brain section as well as magnified images from the cortex, hippocampus and brainstem are shown. Scale bar, 2.5mm (overview), 0.5mm (zoom). Numbers shown next to the schematics report the number of animals in the group positive for the pathology shown as a fraction of the total number of animals in each group.

The automated scrapie cell assay (ASCA)^36,37^ was used to provide measurement of prion titres throughout the disease course (**Fig. 2c**). Prion titres in both RML-infected *Stx6^+/+^* and *Stx6^-/-^*mice increased rapidly at comparable rates before reaching a similar maximal prion titre of ∼10^8.5^ infectious units/g at ∼90 days post inoculation (dpi), in line with the two-phase kinetics model^24,25^. This suggests syntaxin-6 does not alter prion propagation kinetics in established disease. This was further supported by biochemical assessment of disease-related proteinase K (PK)-resistant PrP (PrP^Sc^) at 140 dpi, which was comparable in infected *Stx6^+/+^* and *Stx6^-/-^*mice, with indistinguishable electrophoretic mobility and glycosylation patterns (**Fig. 2d**). This was also corroborated by immunohistochemical detection of disease-related PrP, where we found no differences in the appearance, extent and distribution of PrP deposits with both the onset and evolution of deposition being broadly comparable in RML-infected *Stx6^+/+^* and *Stx6^-/-^* mice (**Fig. 2e**). Taken together, these results suggest that syntaxin-6 is not involved in prion propagation nor in modulating the levels of disease-related PrP during established disease.

Subsequently, we explored whether there were any differences in the onset and/or progression of markers of neurotoxicity or neurodegeneration in RML-infected *Stx6^+/+^* and *Stx6^-/-^* mice. There were no differences in the extent or distribution of intraneuronal vacuoles (“spongiosis”) (**Extended Fig. 1a-b**), synaptic integrity (**Extended Fig. 1c**) or the spatiotemporal evolution of astrogliosis and microgliosis (**Extended Fig. 1d-e**) across the disease course. Furthermore, we observed comparable levels of disease-associated PK-sensitive PrP species (**Extended Fig. 2a-b**) and toxicity levels in prion-infected brain homogenates using a validated neurotoxicity assay^38^ (**Extended Fig. 2c-d**). There were also no consistent differences in serum neurofilament light chain (NfL) levels (**Extended Fig. 2e**), which is a sentinel neurodegeneration biomarker in prion disease^39^. Finally, RML-infected *Stx6^+/+^* and *Stx6^-/-^* mice exhibited comparable neurological phenotypes, time to first symptom (*Stx6^-/-^* median [95% confidence interval] = 129 days [126-129] vs. *Stx6^+/+^* = 129 days [126-131]), incubation times (*Stx6^-/-^* median [95% confidence interval] = 139 days [134-141] vs. *Stx6^+/+^* = 140.5 days [139-144]) and clinical progression (**Extended Fig. 2f-h**). These results suggest syntaxin-6 is not involved in prion-induced neurotoxicity.

Brain transcriptomic analyses of *Stx6^+/+^* and *Stx6^-/-^*mice suggested compensatory mechanisms were not at play at the RNA level with there being no significant upregulation of genes encoding other syntaxins/trafficking proteins (**Supplementary Table 1**). Therefore, taken together with the positive results of the titration study, these findings suggest syntaxin-6 modifies the establishment of disease, with no discernible effect on prion propagation nor prion-induced toxicity in established disease.

### Syntaxin-6 modifies prion-related phenotypes in cellular models with a role in prion trafficking and export

As we had established a role for syntaxin-6 in modulating prion pathogenesis *in vivo*, we proceeded to explore cellular mechanisms which may underlie the observed *in vivo* effect. We stably knocked down *Stx6* in the prion-susceptible PK1 neuroblastoma cell line^36^ by ∼85-90% in 3 independent cell lines relative to cell lines expressing a non-silencing control (NSC) scrambled shRNA (**Extended Fig. 3a-b**). We additionally generated two independent cell lines with ∼9-10-fold overexpression of syntaxin-6 (**Extended Fig. 3d-e**). The SCA was subsequently employed to explore whether syntaxin-6 plays a role in susceptibility to prion infection. The output of this assay is “spot count” with the detection of foci of aggregated PrP (spots detected via the ELISpot assay) differentiating infected and non-infected cells, which allows prion titres to be calculated^36^.

Infection of PK1 *Stx6* knockdown cell lines with RML prions resulted in a statistically robust increase in the spot count, which was broadly consistent across split numbers and prion dilutions (**Fig. 3a-b**). In contrast, the spot count was abolished in cell lines with diminished PrP^C^ levels (PK1 *Prnp* KD1 and KD2), the substrate for conversion, confirming the expected performance of the assay (**Fig. 3b**). Furthermore, when we challenged PK1 *Stx6* knockdown cell lines with a limiting dose of prions, cells became susceptible to prions, in contrast to controls which were resistant to infection (**Extended Fig. 4a-b**). Strengthening these findings, infection of PK1 *Stx6* overexpression cell lines resulted in a robust reduction in the spot count (**Fig. 3a, c**), with there being a strong negative gene-dosage association between *Stx6* levels and susceptibility to prion infection across all of the aforementioned cell lines (Spearman r = 0.85, P = 0.0062) (**Fig. 3d**).

**Figure 3.**
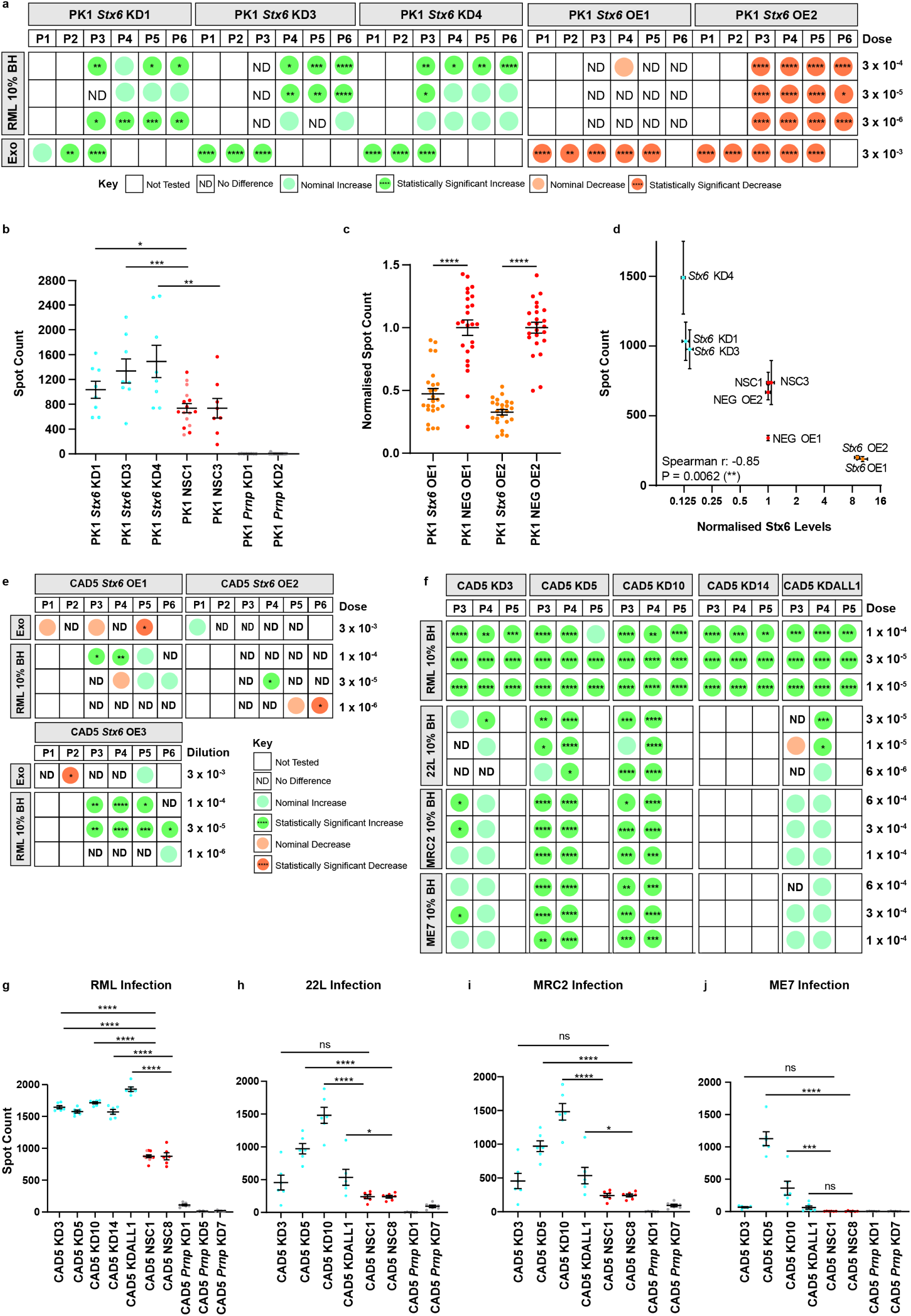
Syntaxin-6 Manipulation in Prion Susceptible Cells Alters Cell-Associated Infectivity Following Prion Infection. **(a)** Matrix summarising the effects of syntaxin-6 manipulation on the spot count of infected cell number in the scrapie cell assay (SCA) in PK1 neuroblastoma cells. The results of three independent PK1 *Stx6* knockdown (KD) cell lines and two PK1 independent *Stx6* overexpression (OE) cell lines are shown with each column representing a different passage number (P1-6). Each row details the infection paradigm, including different dilutions of 10% (w/v) RML-infected brain homogenate (BH) or a crude infected exosome preparation (exo). Statistics are based on one-way ANOVA followed by Fisher’s LSD test of the pre-planned comparison of the relevant control (NSC or NEG OE) to each cell line with *Stx6* manipulation on a plate-by-plate basis. All statistics are based on the raw spot count except for PK1 *Stx6* overexpression cell lines, where the spot count was normalised to the haemotoxylin total cell count. Nominal differences were defined as means which surpassed the threshold of the mean spot count of the negative control ± 0.5 standard deviations. **(b)** Representative example of the spot count of infected cell number at the 5th split in the SCA following infection with 3 x 10^-4^ RML prions (8 technical replicates/cell line). As the assay was conducted across multiple plates, spot counts of subsequent plates were normalised to the mean spot count of the non-silencing control (NSC1) on plate 1. NSC1 was technically replicated across two plates as indicated in separate colours. **(c)** Representative example of the spot count of infected cell number at the 3rd split in the SCA following infection with an infected exosome fraction (3 x 10^-3^; 24 technical replicates/cell line). The spot count was normalised to haemotoxylin total cell count to determine the proportion of cells infected before being normalised to the mean of the relevant NEG OE cell line. **(d)** Graph illustrating the relationship between Stx6 protein level (normalised to the relevant negative control) and the spot count in the SCA shown in the representative examples in (b) and (c) in PK1 cells. The strength of correlation was assessed by the Spearman’s rank correlation coefficient (Spearman r = -0.85, P = 0.0062). **(e, f)** Matrix summarising the effects of syntaxin-6 manipulation on the spot count in the SCA in CAD5 catecholaminergic cells as in (a) with the additional interrogation of infection with other mouse-adapted prion strains including 22L, MRC2 and ME7. **(g-j)** Representative examples of the spot count of infected cell number at the 4th split in the SCA following infection with 1 x 10^-5^ RML prions, 1 x 10^-5^ 22L, 6 x 10^-4^ MRC2 and 6 x 10^-4^ ME7 (6 technical replicates/cell line/strain). As the assay was conducted across multiple plates, spot counts of subsequent plates were normalised to the mean spot count of non-silencing control (NSC1) on plate 1. NSC1 was technically replicated across two plates as indicated in separate colours. Statistical differences were assessed by one-way ANOVA followed by Fisher’s LSD test on planned comparisons. *P < 0.05, **P < 0.01, ***P < 0.001, ****P < 0.0001.

**Figure 4.**
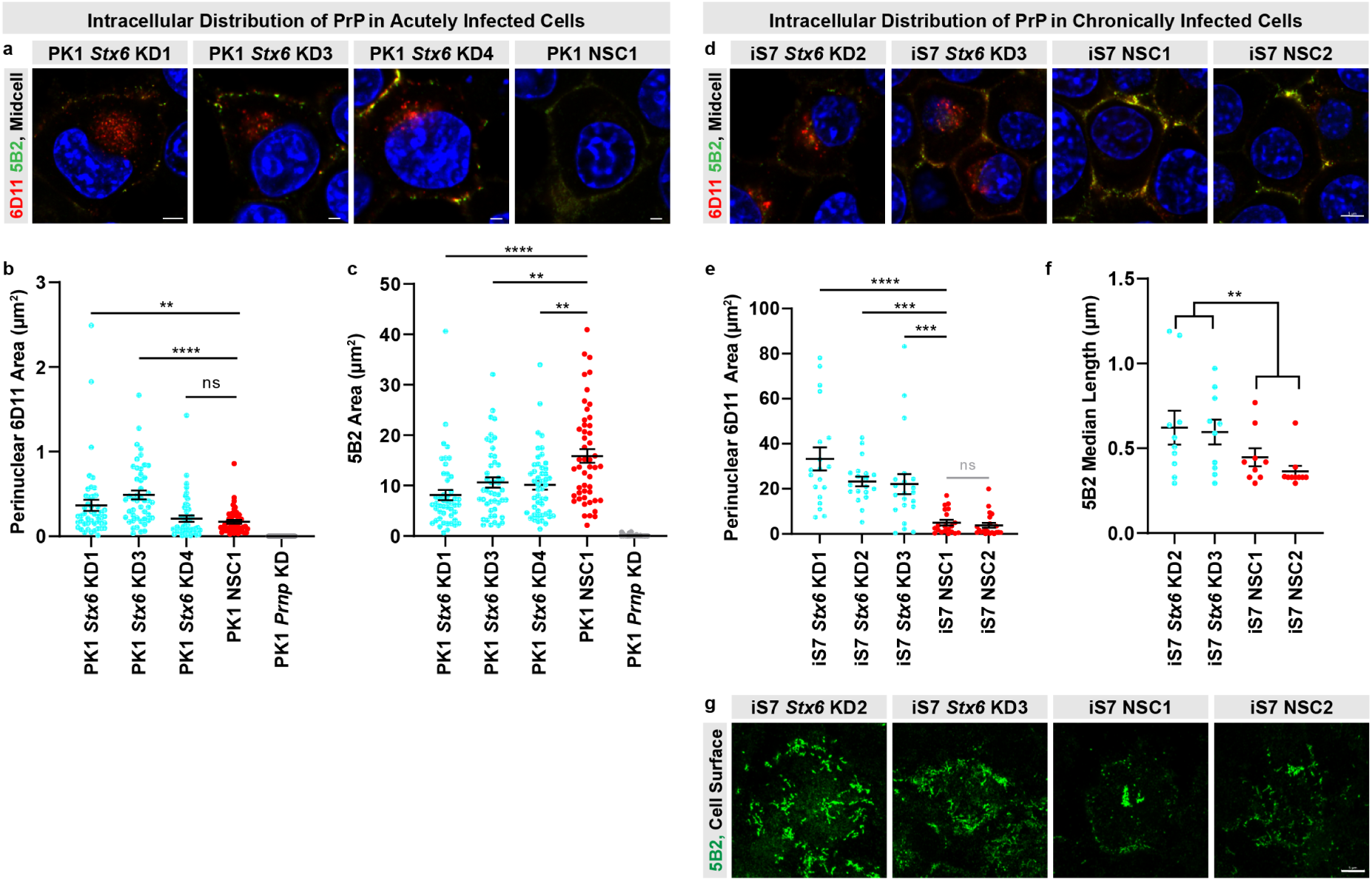
Syntaxin-6 Knockdown Alters the Intracellular Distribution and Aggregate Morphology of Disease-Related PrP in Prion-Infected PK1 Cells. **(a)** Representative images of three independent *Stx6* knockdown cell lines and one non-silencing control (NSC1) stained with the discriminatory anti-PrP antibody pair, 5B2 and 6D11, which preferentially immunolabel disease-related PrP assessed by confocal laser-scanning microscopy. DAPI, nuclear stain, blue; 6D11, red; 5B2, green. Scale bar, 5 μm (*Stx6* KD1) or 2 μm (*Stx6* KD3-4, NSC1). **(b)** Quantification of the 6D11 signal area in the perinuclear space normalised by total cell count as indicated by DAPI. Statistical differences were assessed by one-way ANOVA on log transformed data followed by Fisher’s LSD test on planned comparisons. Each dot represents an individual image (n=47-49/cell line). One outlier (8.28) was excluded in the graph from the *Stx6* KD1 group for visual clarity but was included in the statistical analysis of log-normal data. **(c)** Quantification of the total 5B2 signal area coverage at mid-cell level normalised by total cell count as indicated by DAPI. Each dot represents an individual image (n=47-49/cell line). Statistical differences were assessed by one-way ANOVA on log transformed data followed by Fisher’s LSD test on planned comparisons. **(d)** Representative images of iS7 cells with stable *Stx6* knockdown, or control iS7 cells expressing a NSC shRNA construct, co-stained with two anti-PrP antibodies (6D11, red and 5B2, green) as well as DAPI nuclear stain (blue) at mid cell level. Scale bar, 5 µm. **(e)** Quantification of 6D11-positive area of disease-related PrP in the perinuclear space with the line representing mean ± SEM and individual dots representing an individual cell (19-21 cells/cell line). Statistical differences were assessed by one-way ANOVA followed by Fisher’s LSD test on planned comparisons. **(f)** Quantification of the median length of elongated disease-related PrP aggregates derived from maximum intensity projections at the plasma membrane and extracellular matrix level (9-10 images/cell line). Statistical differences were assessed on log-normal data using one-way ANOVA to test for the effect of *Stx6* expression level on length. **(g)** High magnification images of elongated disease-related PrP aggregates at plasma membrane and extracellular matrix level displayed as maximal intensity projections of z-stack images to capture the length across all planes of the aggregate. Scale bar, 5 µm. *P < 0.05, **P < 0.01, ***P < 0.001, ****P < 0. 0001.

As we had found profoundly altered prion-related phenotypes in PK1 cells with syntaxin-6 manipulation, we additionally manipulated the expression of syntaxin-6 in CAD5 catecholaminergic cells^40^ to increase the generalisability of our findings. Therefore, 5 independent CAD5 cell lines with ∼59%-80% knockdown of syntaxin-6 were generated (**Extended Fig. 3g-i**), as well as 3 independent cell lines with ∼8-14-fold overexpression of syntaxin-6 (**Extended Fig. 3k-l**). Although we found no consistent effect in *Stx6* overexpressing CAD5 cells (**Fig. 3e**), *Stx6* knockdown in CAD5 cells unequivocally exhibited an increased spot count in response to RML infection, corroborating the PK1 data (**Fig. 3f-g**). Strengthening these findings, this extended to other mouse-adapted prion strains including 22L (**Fig. 3h**), MRC2 (**Fig. 3i**) and ME7 (**Fig. 3j**) prions. Collectively, these results suggest syntaxin-6 modifies susceptibility to prion infection in cellular models, with confidence stemming from the largely congruent effects observed across multiple different paradigms, prion strains and cell types.

To discount confounding factors underlying these results, we confirmed broadly comparable growth rates of the cell lines (**Extended Fig. 3c, f, j, m**) and determined that the differing spot counts were not reflective of differences in cell viability (**Extended Fig. 4g-i**). There were no consistent differences in total PrP levels (**Extended Fig. 4j, l, m**) with the exception of a subtle reduction in PK1 cells with *Stx6* overexpression (**Extended Fig. 4k**). Although this is unlikely to be a driver of the altered spot count given the modesty of the difference and the evidence that the rate of prion propagation is not a function of PrP expression in cells^41–44^, we explored the underlying driver of this potential epiphenomenon. There were no differences in *Prnp* mRNA levels (**Extended Fig. 4n**) or PrP^C^ degradation kinetics (**Extended Fig. 4o**) arguing against a role for syntaxin-6 in PrP^C^ synthesis or its intracellular degradation in cellular models respectively. Therefore, we found no evidence for confounders underlying the SCA results.

Due to the prominent effects on the spot count, we performed complementary phenotypic characterisation of the infected *Stx6* knockdown cell lines by confocal microscopy using two antibodies, 5B2 and 6D11, which each detect a distinct disease-associated PrP staining profile in this cellular system^45^. 5B2 immunostaining recognises elongated disease-related PrP aggregates at the plasma membrane and extracellular matrix in infected cells, whereas 6D11 immunostaining is seen as punctate, perinuclear staining in addition to some plasma membrane and extracellular matrix staining. There was a statistically significant increase in the accumulation of perinuclear 6D11-positive disease-related PrP in PK1 *Stx6* KD1 and KD3 (**Fig. 4a-b**) as well as a statistically robust reduction in 5B2-positive plasma membrane staining in all *Stx6* knockdown cell lines (**Fig. 4c**). This demonstrates a redistribution of disease-related PrP with syntaxin-6 knockdown, in keeping with a trafficking mechanism of action. Confocal imaging of infected *Stx6* overexpression cells revealed a universal reduction in 6D11 and 5B2 staining (**Extended Fig. 4c-f**) suggesting that syntaxin-6 overexpression was sufficient to almost clear the infection, corroborating the SCA data.

To more thoroughly explore phenotypic differences in the distribution and aggregate morphology of disease-related PrP, we additionally stably knocked down syntaxin-6 in chronically infected PK1 cells (a subclone called iS7)^45^ allowing us to explore a role for syntaxin-6 in prion accumulation isolated from infection (**Extended Fig. 5a-b**). Recapitulating our observations with freshly infected cells (**Fig. 4a-b**), we found a prominent accumulation of perinuclear 6D11-positive disease-related PrP with *Stx6* knockdown in chronically infected cells (**Fig. 4d-e**). We also observed longer 5B2-positive elongated disease-related PrP aggregates at the plasma membrane and extracellular matrix with *Stx6* knockdown (P = 0.0062) (**Fig. 4f-g**) but not total load (**Extended Fig. 5i**). However, secreted infectivity (**Extended Fig. 5g**) and prion steady-state levels (**Extended Fig. 5h**) were comparable with syntaxin-6 manipulation with there being no consistent differences in total PrP, PrP^Sc^ or disease-related PrP (**Extended Fig. 5c-f, j-k**). Taken together, these results demonstrate a cellular redistribution and structural reorganisation of disease-related PrP in chronically infected cells with syntaxin-6 knockdown lending further support for syntaxin-6 mediated trafficking of disease-related PrP.

To test whether a role for syntaxin-6 in prion export could explain the altered spot counts, we transiently knocked down syntaxin-6 in chronically infected iS7 PK1 cells followed by collecting conditioned media to infect PK1 reporter cells (**Fig. 5a**). Here, we saw a robust reduction in secreted infectivity with syntaxin-6 knockdown measured by the SCA (18.2% ± 3.7% (mean ± SEM) reduction) (**Fig. 5b**). Corroborating this, when we harvested media from the infected stable *Stx6* knockdown and overexpression PK1 cell lines described previously, syntaxin-6 knockdown resulted in reduced relative secreted infectivity titres, with the converse being observed with syntaxin-6 overexpression, following correction for baseline differences in cell-associated infectivity (**Extended Fig. 4p, q**). In further support for a role of syntaxin-6 in prion export, as opposed to involvement in an intracellular trafficking step, we observed a generalised increase in the degree of colocalisation of 6D11-positive disease-related PrP with markers of intracellular organelles (**Fig. 5c-f**). Syntaxin-6 knockdown resulted in increased colocalisation of 6D11 with the early endosome marker, EEA1, (**Fig. 5d**) and the lysosome marker, LAMP1, (**Fig. 5e**) with some additional evidence for the TGN marker, TGN46 (**Fig. 5f**). Collectively, this provides strong evidence for prion export being the syntaxin-6-driven molecular susceptibility mechanism. This mechanism can consolidate the observations made in cells (with export facilitating clearance, enabled by regular cell media replenishment) with the risk effect observed *in vivo* (with export facilitating spread and the establishment of infection).

**Figure 5.**
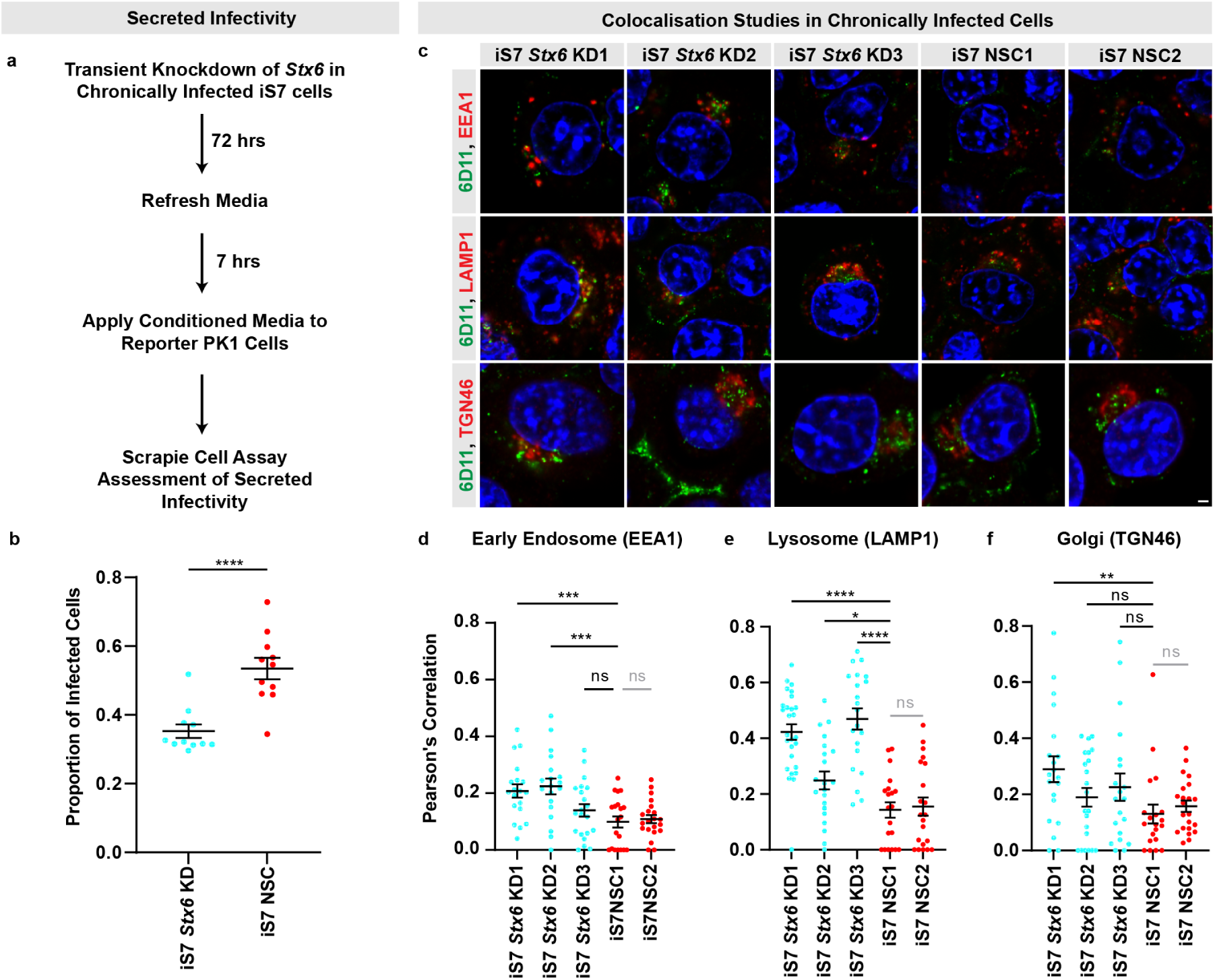
Syntaxin-6 Modifies Prion Export in Chronically Infected PK1 Cells. **(a)** Experimental design for assessing a modifying effect of syntaxin-6 knockdown on secreted infectivity from chronically infected cells (iS7 subclone). **(b)** The proportion of infected reporter PK1 cells (spot count normalised to haemotoxylin total cell count) following the application of conditioned media harvested from iS7 cells in which *Stx6* has been transiently knocked down or a non-silencing control (NSC) shRNA had been employed. **(c)** Representative images of iS7 cells co-labelled with 6D11 and markers of intracellular compartments to assess for altered distribution of disease-related PrP with syntaxin-6 knockdown in the early endosome (EEA1), lysosomes (LAMP1) and in the *trans*-Golgi (TGN46). Scale bar, 2 µm. **(g-i)** Assessment of levels of colocalisation between 6D11-positive PrP and organelle markers, expressed as Pearson’s correlation coefficient for EEA1 (n=19-21 cells/cell line), LAMP1 (n=20-27 cells/cell line) and TGN46 (n=20-22 cells/cell line). Line represents mean ± SEM with one-way ANOVA followed by Fisher’s LSD test being used to test statistical differences. *P < 0.05, **P < 0.01, ***P < 0.001, ****P < 0. 0001.

Therefore, in addition to modulating early stages of prion pathogenesis *in vivo,* this establishes a role for syntaxin-6 in altering prion-related phenotypes in cellular models with syntaxin-6-mediated prion export being the likely molecular susceptibility mechanism.

### Protective effect of syntaxin-6 knockout on numerous physiological, behavioural, neuropathological and clinical outcome measures in a transgenic P301S tauopathy model

Given that *STX6* is a shared genetic risk factor, we hypothesised that syntaxin-6 may also be a modifier of tau pathogenesis *in vivo* with a common susceptibility mechanism. Therefore, to functionally validate and study a role for syntaxin-6 in tau pathogenesis, we crossed *Stx6^-/-^* mice with a transgenic P301S mutant tauopathy mouse model (h*Tau*^P301S/P301S^ mice)^34^, which recapitulates many of the molecular and cellular features of human tauopathies. We subsequently conducted a longitudinal time course investigation of *Stx6^+/+^;*h*Tau*^P301S/P301S^ and *Stx6^-/-^;*h*Tau*^P301S/P301S^ mice, assessing numerous behavioural, clinical, neuropathological, biochemical and molecular phenotypes (**Fig. 6a**).

**Figure 6.**
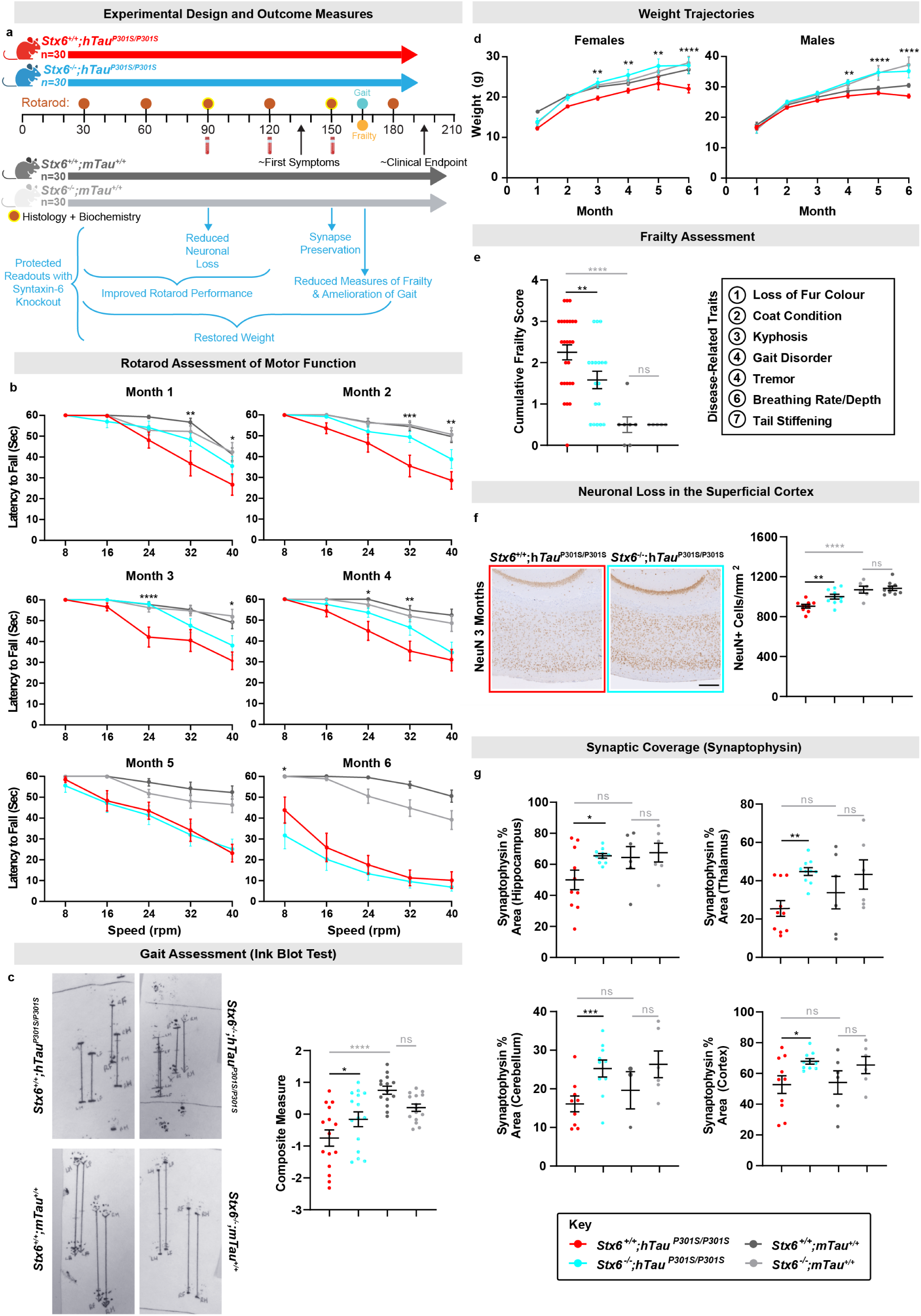
Protective Effect of Syntaxin-6 Knockout on Numerous Behavioural, Physiological and Neuropathological Outcome Measures in a Transgenic Tauopathy Mouse Model. **(a)** Overview of the experimental design summarising the four arms of the study and timeline of the different outcome measures assessed with a summary of the protective readouts with syntaxin-6 knockout in tauopathy mice below. (b) *Stx6*^+/+^;h*Tau*^P301S/P301S^, *Stx6*^-/-^;h*Tau*^P301S/P301S^, *Stx6*^+/+^;m*Tau*^+/+^ and *Stx6*^-/-^;m*Tau*^+/+^ mice (n = 15/genotype, mixed sex) were subjected to 5 different speeds on a rotarod (8–40 rpm), receiving two trials/speed. The latency to fall (maximum trial length = 60 s) was recorded at each speed in the two trials. Data represent mean latency ± SEM. A 3-way repeated measures mixed-effect model was fitted to the data at each month with genotype and sex as factors and speed as a repeated factor. Stars relate to the results of the Fisher’s LSD post-hoc test of the pre-planned comparison between *Stx6*^+/+^;h*Tau*^P301S/P301S^ and *Stx6*^-/-^;h*Tau*^P301S/P301S^ mice. **(c)** Gait assessment in 5.5-month-old mice (n = 15/genotype, mixed sex). Left: representative images of ink blots. Right: graph showing the mean composite measure ± SEM of disease-relevant gait parameters. A Z-score was calculated for each prioritised disease-relevant gait parameter per animal. Each dot represents the average Z score/mouse, which were analysed using one-way ANOVA with genotype as the factor of interest and sex as a blocking factor followed by Fisher’s LSD test of preplanned comparisons. **(d)** Body weight of female (left) and male (right) mice at 1–6 months (n=5-10/sex/genotype). Data are expressed as mean body weight (g) ± SEM. Statistical differences were determined using a 2-way repeated measures mixed model approach with genotype as the treatment factor and month as the repeated factor. As there was a significant interaction between genotype and month, each month was assessed separately using Fisher’s LSD test on the preplanned comparison between *Stx6*^+/+^;h*Tau*^P301S/P301S^ mice and *Stx6*^-/-^;h*Tau*^P301S/P301S^ mice. **(e)** Frailty assessment in *Stx6*^+/+^;h*Tau*^P301S/P301S^ (n=26), *Stx6^-^*^/-^;h*Tau*^P301S/P301S^ (n=18), *Stx6*^+/+^;m*Tau*^+/+^ (n=7) and *Stx6*^-/-^;m*Tau*^+/+^ (n=5) mice. Graph shows cumulative frailty score (18 parameters each assessed using a 3-point score: normal (0), moderate (0.5) or severe (1)). Each dot represents the cumulative frailty score for an individual mouse. Following rank transformation, the data were analysed using one-way ANOVA, with genotype as the treatment factor, with sex and batch as blocking factors, followed by Fisher’s post-hoc LSD test of preplanned comparisons. **(f)** Representative images of NeuN staining in the superficial cortex (500 μm deep from the cortical surface) in *Stx6*^+/+^;h*Tau*^P301S/P301S^ and *Stx6^-^*^/-^;h*Tau*^P301S/P301S^ mice at 3 months (left). Scale bar represents 250 μm. Quantification of image-based threshold analysis of NeuN staining in superficial cortex (right) (n=6-10/genotype). One-way ANOVA was performed with genotype as the treatment factor and sex as blocking factor, followed by Fisher’s LSD post-hoc test of pre-planned comparisons. **(g)** Quantification of synaptophysin staining in different brain regions in *Stx6*^+/+^;h*Tau*^P301S/P301S^ and *Stx6^-^*^/-^;h*Tau*^P301S/P301S^ mice (n=5-10/genotype, mixed sex) at 5 months. A 2-way repeated measures mixed model approach was used for statistical analysis using the unstructured covariance structure to model the within-subject correlations, with genotype as the treatment factor, brain region as the repeated factor and sex as a blocking factor. This was followed by planned comparisons on the predicted means to compare the effect of genotype on staining in the different brain regions. The brain stem was also assessed with there being no difference between *Stx6*^+/+^;h*Tau*^P301S/P301S^ and *Stx6^-^*^/-^;h*Tau*^P301S/P301S^ mice. *P < 0.05, **P < 0.01, ***P < 0.001, ****P < 0.0001.

As h*Tau*^P301S/P301S^ mice exhibit a prominent, progressive motor impairment^46^, we assessed whether syntaxin-6 knockout modifies rotarod performance (**Fig. 6b**). From month 1, *Stx6*^+/+^;h*Tau*^P301S/P301S^ mice displayed a motor defect compared to non-transgenic control *Stx6*^+/+^;m*Tau*^+/+^ mice, which persisted across the 6 months examined. In contrast, there was evidence for a partial rescue of the motor impairment in *Stx6^-/-^;*h*Tau*^P301S/P301S^ mice from months 1-4. Although rotarod performance converged in months 5 and 6, when mice were presenting with overt and debilitating clinical signs, syntaxin-6 knockout still exerted a protective effect in 5.5-month-old mice when we assessed gait as a less challenging assessment of motor dysfunction. Here, prioritised, disease-relevant gait parameters were harmonised into a composite score for each animal, validating a robust deficit in gait in *Stx6*^+/+^;h*Tau*^P301S/P301S^ mice relative to control *Stx6*^+/+^;m*Tau*^+/+^ mice (P < 0.0001) (**Fig. 6c**). In contrast, there was partial rescue of this impairment in *Stx6^-/-^;*h*Tau*^P301S/P301S^ mice (P = 0.033), providing an independent measure of functional amelioration with syntaxin-6 knockout.

As we had found that syntaxin-6 reduction mitigated some of the detrimental effects of tauopathy at the functional level, we also assessed physiological outcome measures. Weight trajectories provide an objective readout of general health and wellbeing, which were compromised in *Stx6*^+/+^;h*Tau*^P301S/P301S^ mice (**Fig. 6d**). Conversely, *Stx6^-^*^/-^;h*Tau*^P301S/P301S^ mice showed evidence of weight normalisation, which serves as a positive correlate of disease amelioration. We also conducted an 18-point clinical exam in 5.5-month-old animals (derived from a validated frailty assessment^47^) (**Fig. 6e**). *Stx6*^+/+^;h*Tau*^P301S/P301S^ mice exhibited greater measures of frailty than the control *Stx6*^+/+^;m*Tau*^+/+^ mice (P < 0.0001), which was partially rescued in *Stx6*^-/-^;h*Tau*^P301S/P301S^ mice (P = 0.0052). In 6/7 disease-relevant frailty parameters, *Stx6*^+/+^;h*Tau*^P301S/P301S^ mice showed a higher proportion of animals with impairments (**Supplementary Table 2**). Although no differences were observed in the more subjective endpoint clinical outcome measures (**Supplementary Table 3**), the prominent functional rescue observed in these more sensitive functional tests demonstrates that syntaxin-6 knockout favourably modifies the disease course in this transgenic tauopathy model.

To explore whether differences in pathological tau species were associated with the prominent functional rescue with syntaxin-6 knockout, we assessed established pathological tau epitopes by either immunohistochemistry or biochemistry. Surprisingly, immunohistochemical characterisation of AT8-positive phospho-tau in the spinal cord was robustly increased in *Stx6^-^*^/-^;h*Tau*^P301S/P301S^ mice relative to *Stx6*^+/+^;h*Tau*^P301S/P301S^ mice at 3 months but not at 5 months (**Extended Fig. 6**). There was also a subset of brain regions with elevated AT8-positive phospho-tau or MC1-positive misfolded tau in *Stx6*^-/-^;h*Tau*^P301S/P301S^ mice (**Extended Fig. 7**). However, we did not observe any differences in AT8-positive or PHF1-positive phospho-tau species in total brain homogenates by western blot at either 3 months or 5 months (**Extended Fig. 8a-d**) and total tau levels were comparable (**Extended Fig. 8e**). There were also no differences in seeded tau aggregation when we applied brain homogenates prepared from 2- month-old mice to HEK tau biosensor cells^26^ (**Extended Fig. 8f-h**). Given that we observed localised increases of tau pathology in *Stx6*^-/-^;h*Tau*^P301S/P301S^ mice by immunohistochemistry despite the total biochemical load of phospho-tau species and seeding-competent species remaining constant, these results are in keeping with altered trafficking of pathological tau species with syntaxin-6 knockout, with a higher proportion of tau species being in a highly aggregated state. Furthermore, these results support an uncoupling of the functional outcome measures with the levels of pathological tau, which has been observed previously (see discussion).

Despite the persistence of tau pathology, we hypothesised that we may observe a protective effect of syntaxin-6 knockout on neuropathological hallmarks related to function given the evidence of rescue at the behavioural level. Therefore, we quantified NeuN-positive cell density in the superficial cortex, which exhibits progressive neuronal loss in this tauopathy model^48^. We recapitulated this finding with 3-month-old *Stx6^+/+^*;*hTau*^P301S/P301S^ mice having a significant reduction in the number of NeuN-positive cells/mm^2^ relative to control *Stx6^+/+^;*m*Tau^+/+^* mice (P < 0.0001), which was partially rescued with syntaxin-6 knockout (P = 0.0055) (**Fig. 6f**). Although neuronal loss was comparable between *Stx6*^+/+^;h*Tau*^P301S/P301S^ and *Stx6^-^*^/-^;h*Tau*^P301S/P301S^ mice by 5 months (**Extended Fig. 9a**), we observed evidence of increased synaptic integrity with syntaxin-6 knockout at this time point (**Fig. 6g**). Indeed, synaptic coverage had been reported in the literature to be reduced in the hippocampus, cortex, brain stem and cerebellum in this tauopathy mouse model^49^. We found higher synaptophysin staining coverage with syntaxin-6 knockout in the hippocampus (P = 0.0151), thalamus (P = 0.0023), cerebellum (P = 0.0007) and the cortex (P = 0.0127) at 5 months. However, neither neuroinflammatory phenotypes nor serum NfL levels correlated with the protective effect of syntaxin-6 knockout (**Extended Fig. 9b-d**). Therefore, these results suggest a partial rescue of neurodegeneration-related neuropathological hallmarks despite the persistence of pathological tau species and neuroinflammation.

Collectively, our results have demonstrated evidence for a protective effect of syntaxin-6 knockout on numerous behavioural, physiological and neuropathological outcome measures, as well as changes in the distribution/aggregation state of pathological tau species in keeping with an intracellular trafficking mechanism of action.

## Discussion

We provide evidence that syntaxin-6 may be a common modifier of prion (or prion-like) disease in cell and mouse models with distinct pathologies. This provides functional support for previous GWAS findings and mechanistic insight into the risk imparted by syntaxin-6 variants in sCJD and PSP. We show that syntaxin-6 knockout *in vivo* reduces susceptibility to prion infection, suggesting a role for syntaxin-6 in disease establishment. We provide evidence for altered intracellular trafficking and export of prions as a cellular susceptibility mechanism which may underlie this observation. We further show that syntaxin-6 knockout in a transgenic tauopathy mouse model resulted in a functional amelioration of numerous physiological, behavioural and neuropathological outcome measures consistent with relevance of syntaxin-6 to multiple neurodegenerative diseases which share prion-like features.

The influence of syntaxin-6 on prion susceptibility in prion-infected mice aligns with the human genetic data linking increased expression of syntaxin-6 to higher disease risk^10,18^. However, knockout did not affect prion propagation kinetics or neurotoxicity in established disease, mirroring the human findings that *STX6* risk variants do not impact rates of sCJD progression^50^. This aligns with other work suggesting a distinction between genetic susceptibility factors and drivers of disease progression^51–53^.

In cellular models, syntaxin-6 levels inversely correlated with prion infection levels measured by the SCA. This was recapitulated with *Stx6* manipulation in several cell lines in a strain-independent manner, demonstrating the generalisability of these findings. Underlying this, syntaxin-6 knockdown diminished prion infectivity in conditioned media, consistent with a proposed function of syntaxin-6 in the export of proteins from cells^55,56^, a key mechanism of prion spread^57,58^. This suggests a novel prion cellular susceptibility mechanism linked to prion export, which enhances prion clearance from infected cells.

Syntaxin-6 knockdown in cells additionally resulted in a conspicuous redistribution of disease-associated PrP to a perinuclear compartment, in keeping with syntaxin-6 being involved in trafficking of disease-associated PrP. Indeed, previous work showing disease-associated PrP colocalises with syntaxin-6 provides a biologically plausible basis for this^59^. However, we did not find evidence of a block at any specific intracellular trafficking step; instead, we observed evidence of increased disease-associated PrP colocalisation with markers of every intracellular compartment examined. This generalised increase in intracellular load of disease-associated PrP provides a further line of evidence for syntaxin-6 mediating prion export.

We additionally observed longer disease-associated PrP aggregates at the plasma membrane in chronically infected cells with syntaxin-6 knockdown, suggesting enhanced fibrillisation kinetics. This aligns with previous *in vitro* work suggesting that syntaxin-6 inhibits the ordered formation of PrP fibrils through direct binding^59^. Despite observing profound phenotypic differences in chronically infected cells with syntaxin-6 knockdown, there were no changes in total prion load during established infection, in line with the *in vivo* results. Collectively, the cell-based work supports a role for syntaxin-6 in disease-associated PrP trafficking and export, further supporting the contention that vesicle trafficking is a key mechanism contributing to prion pathogenesis^60–64^.

Given the genetic link of *STX6* to tauopathies, we extended our analysis to a tauopathy mouse model^34^, demonstrating that syntaxin-6 knockout mitigated motor deficits, frailty measures, and neurodegeneration-related pathology. Interestingly, despite increased localised AT8- and MC1-positive tau pathology, total phospho-tau and seeding-competent tau levels remained unchanged, suggesting an altered pathological tau distribution akin to the prion-related findings. This can be consolidated with reduced tau export with syntaxin-6 knockout, aligning with prior reported cellular evidence^65^. Furthermore, the observed dissociation between tau burden and toxicity aligns with previous studies showing functional preservation despite persistent pathology^27–33^. These findings emphasise intracellular trafficking as a key pathway in neurodegeneration, consistent with other GWAS-identified susceptibility factors such as *SORL1*, *BIN1*, and *PICALM*^66,67^.

Targets underpinned by human genetics evidence increase the likelihood of therapeutic translation^68–71^. Although PrP and tau lowering offers a genetically validated therapeutic approach in prion disease and tauopathies respectively, with preclinical efficacy^39,72,73^ and target engagement/safety demonstrated in humans^74–76^, extending our therapeutic repertoire would be valuable. Our results suggest that the modifying effects of syntaxin-6 do not lend themselves well to drug development in symptomatic patients. Indeed, we show that syntaxin-6 reduction had no effect in established mouse prion disease and the effects occurred primarily in the initial stages of the disease in a tauopathy mouse model. Given its early modifying effect, *STX6*-lowering could potentially be useful in prevention in the at-risk patient population, such as genetic mutation carriers. This resonates with recent work suggesting that targeting GWAS-identified susceptibility factors is more likely to yield efficacy in pre-symptomatic populations^51^.

Limitations of this work include the use of acquired models of prion infection, resulting in a disconnect with the risk-conferring effect identified in sporadic human aetiology^10^. Although we employed a tauopathy mouse model which develops disease without requiring inoculation with tau fibrils, this was driven by the overexpression of human tau with a rare FTD-associated mutation^77–80^. Of note, this model generates tau filaments which are structurally distinct to known wild type tau filaments in human tauopathy patients (although tau filament structures in FTD patients harbouring the P301S mutation are not yet known)^81^. Nevertheless, due to the paucity of “better” models, this work provides an important proof-of-concept to inform future studies.

In conclusion, we propose syntaxin-6 as a modifier of both prion and tau early pathogenesis *in vivo* and provide evidence for syntaxin-6 altering the trafficking and export of prions. Therefore, this work has advanced our overall understanding of the relevance of syntaxin-6 to neurodegenerative diseases as a pleiotropic risk factor, informing on the disease stage it is acting, as well as proposing a fundamental susceptibility mechanism of neurodegeneration.

## Funding

The work was funded by the Medical Research Council (UK). SM and JC are National Institute for Health Research (NIHR) Senior Investigators.

## Conflicts of Interest

J.C. is the director of D-Gen, Ltd., an academic spin-out company working in the field of prion disease diagnosis, decontamination and therapeutics. J.C. and G.S.J. are shareholders of D-Gen. D-Gen supplied the ICSM35 antibody used for PrP immunohistochemistry and ICSM18 antibody used for cell experiments. The other authors declare no competing interests.

## Acknowledgements

We would like to thank Jonathan Wadsworth, Selina Wray, Henrik Zetterberg, Iryna Benilova, Emmanuel Risse, Nour Majbour, Jan Bieschke, Malin K Sandberg, Adam Wenborn, Marc Diamond and Michel Goedert for providing advice and/or resource sharing. We would like to acknowledge Stephanie Canning and Azy Khalili for assistance with flow cytometry. We would also like to thank the Biomarkers Lab at UCL, UCL genomics, all staff at our biological services facility and our histology team including Connor Preston, Tamsin Nazari, Jordin Orellana and Yau Mun Lim, as well as Richard Newton for help with graphical design.

## Extended Figures

**Extended Figure 1.**
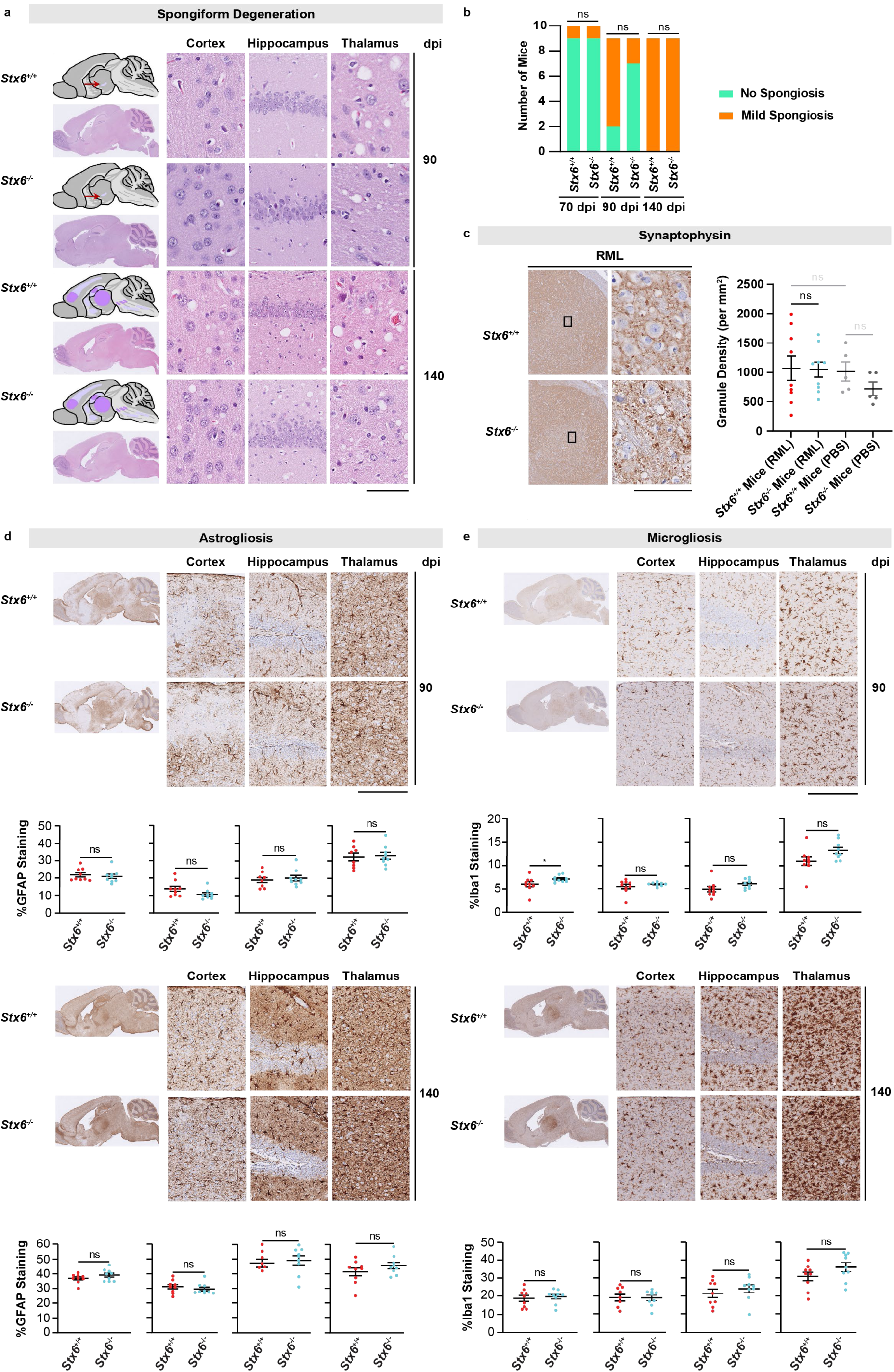
Spatiotemporal Development and Load of the Neuropathological Hallmarks of Prion Disease are Comparable in RML-Infected *Stx6^+/+^* and *Stx6^-/-^* Mice. **(a)** Spongiform vacuolation was assessed by haematoxylin and eosin (H&E) staining with schematics (left) showing the regional distribution of spongiosis (light lilac, mild spongiosis; medium lilac, moderate spongiosis; dark purple, severe spongiosis) with the red arrow indicating the mild spongiosis seen in the inferior thalamus at 90 days post inoculation (dpi). Representative images of whole brain sections are shown below the schematics as well as the cortex, hippocampus and thalamus at 90 and 140 dpi (right). Scale bar, 60 μm (cortex and thalamus), 120 μm (hippocampus), 1.5 mm (whole section). **(b)** Scoring of spongiosis with statistical differences at each time point assessed using Fisher’s exact test. **(c)** Overview of synaptophysin staining in the thalamus in RML-inoculated *Stx6^+/+^* and *Stx6^-/-^* mice. The square indicates the approximate area corresponding to the high magnification image. Scale bar, 1 mm (left), 50 μm (right). Quantification of synaptic granule density in the thalamus at 140 dpi is shown on the right. Statistical differences were assessed using one-way ANOVA with the planned comparison between RML-infected *Stx6^+/+^* and *Stx6^-/-^* mice. **(d)** Astrogliosis was assessed by quantification of GFAP staining at 90 dpi and 140 dpi in neuropathologically validated infected animals. Representative images are shown with the quantification indicated below. Post-rank transformation, a 2-way repeated measures mixed model approach was used for statistical analysis using the unstructured covariance structure to model the within-subject correlations, with genotype as the treatment factor and brain region as the repeated factor. This was followed by planned comparisons on the predicted means to compare the effect genotype on staining in the different brain regions. Scale bar, 250 μm (cortex, hippocampus, thalamus), 2.5 mm (whole section). **(e)** Microgliosis was assessed by quantification of Iba1 staining at 90 dpi and 140 dpi in neuropathologically validated infected animals. Representative images are shown with the quantification indicated below. Post-rank transformation, a 2-way repeated measures mixed model approach was used for statistical analysis using the unstructured covariance structure to model the within-subject correlations, with genotype as the treatment factor and brain region as the repeated factor. This was followed by planned comparisons on the predicted means to compare the effect of genotype on staining in the different brain regions. Scale bar, 250 μm (cortex, hippocampus, thalamus), 2.5 mm (whole section).

**Extended Figure 2.**
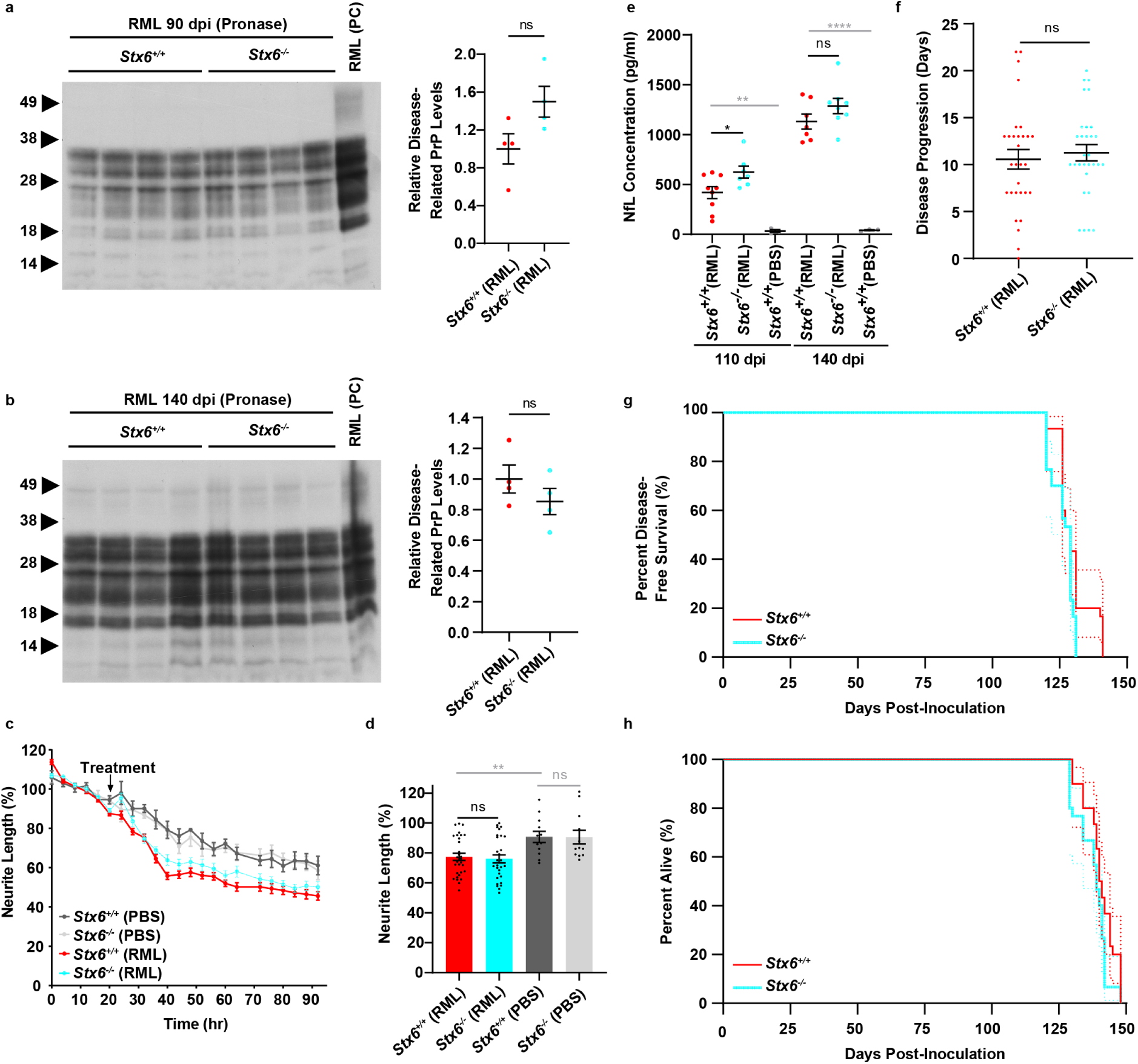
No Differences in Neurotoxicity-Related Outcome Measures in RML-Infected *Stx6^+/+^* and *Stx6^-/-^* Mice. **(a)** Immunoblotting with the anti-PrP antibody, ICSM35, of brain homogenates from mice 90 days post inoculation (dpi), following pronase E digestion (100 µg/ml, 37°C, 30 min). The corresponding semi-quantification of the disease-related PrP signal is shown on the right. Each sample was normalised to the average of RML-infected *Stx6^+/+^* mice with statistical differences assessed using the Student’s t-test. Bar graphs represent mean ± SEM of 4 biological replicates/genotype. PC, positive control. **(b)** Pronase digested brain homogenates at 140 dpi. **(c)** Representative neurotoxicity assay showing normalised neurite retraction over time in primary neurons in response to treatment with a 2.5 x 10^-4^ concentration of RML-infected *Stx6^+/+^* and *Stx6^-/-^* brain homogenates from mice culled at 140 dpi (n=10 technical replicates). PBS-inoculated *Stx6^+/+^* and *Stx6^-/-^* controls culled at 140 dpi were also analysed in quadruplicate. Graph shows mean ± SEM. **(d)** Neurite retraction 12 hrs post-treatment in 3 independent cell cultures with statistical differences assessed using one-way ANOVA followed by Fisher’s LSD post-hoc test. **(e)** Serum NfL levels in RML-infected *Stx6^+/+^* and *Stx6^-/-^* mice at 110 dpi (8-9/genotype) and 140 dpi (7-9/genotype) were measured. PBS-inoculated *Stx6^+/+^* mice (n=5/time point) were also included as controls. Graphs show mean ± SEM, with symbols representing individual animals. Data were analysed using one-way ANOVA followed by Fisher’s LSD post-hoc test. **(f)** Graph showing disease progression, defined as the difference between time to first symptom to prion disease diagnosis (days). P-value derived from an unpaired t-test. **(g)** Kaplan Meier plot showing disease-free survival probability until animals developed first symptoms of prion disease. Dashed lines represent 95% confidence intervals. **(h)** Kaplan Meier plot showing incubation periods (time from inoculation to definite prion disease diagnosis). *P < 0.05, **P < 0.01, ****P < 0.0001.

**Extended Figure 3.**
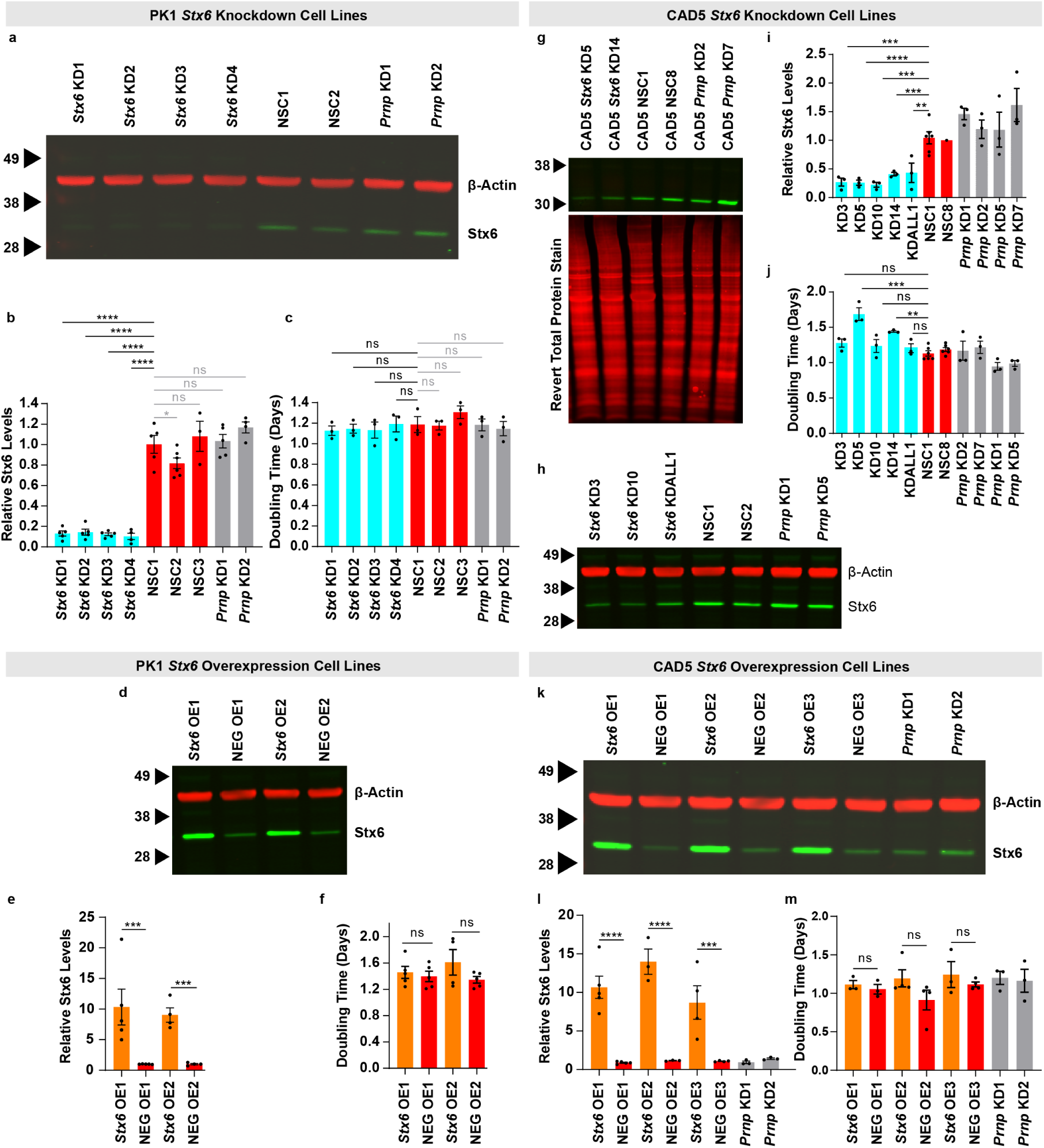
Characterisation of Prion Susceptible Cell lines with Syntaxin-6 Manipulation. **(a)** Immunoblot probing for syntaxin-6 levels (bottom, green) and β-Actin (top, red) in PK1 cells stably expressing shRNAs targeting *Stx6* or *Prnp* or shRNAs encoding a non-silencing control scrambled sequence (NSC1-2). **(b)** Quantification of syntaxin-6 band intensity relative to the β-Actin loading control followed by normalisation to the average normalised signal of the NSC and *Prnp* knockdown cell lines. Bars show mean ± SEM. Each dot represents a different biological replicate with cells being harvested at different passage numbers to confirm stable knockdown. Statistical differences were assessed using one-way ANOVA followed by Fisher’s LSD test assessing differences of all cell lines to NSC1. **(c)** Growth rate measured as the doubling time in days (n=3 biological replicas). Bars and error bars: mean ± SEM. Significance levels based on one-way ANOVA with Fisher’s LSD post-hoc test comparing to NSC1. **(d)** Immunoblot probing for syntaxin-6 levels and β-Actin in PK1 cells which have either been stably transfected with a vector encoding the *Stx6* open reading frame (*Stx6* OE) or a vector encoding an unrelated sequence (NEG OE). **(e)** Quantification of syntaxin-6 band intensity normalised to the β-Actin loading control followed by normalisation to the corresponding negative control line. Bars show mean ± SEM. Following rank transformation, statistical differences were assessed using one-way ANOVA followed by the pre-planned comparisons of matched cell lines. **(f)** Growth rate measured as the doubling time in days (n=3-4 biological replicas). Bars and error bars: mean ± SEM. Significance levels based on one-way ANOVA with Fisher’s LSD post-hoc test of pre-planned comparisons of matched cell lines. **(g,h)** Representative immunoblot for syntaxin-6 levels as well as either total protein stain (g) or β-Actin (h) in CAD5 cells stably expressing shRNAs targeting *Stx6* or *Prnp* or shRNAs encoding a NSC sequence. **(i)** Quantification of syntaxin-6 band intensity normalised to either total protein levels or β-Actin, relative to NSC8. Statistical differences were assessed using one-way ANOVA followed by Fisher’s LSD test assessing differences of all cell lines to NSC1. **j)** Growth rate measured as the doubling time in days (n=3 biological replicas). Bars and error bars: mean ± SEM. As the assessment was conducted across three batches, batch 2 and 3 values were normalised to NSC1 in batch 1 for visual comparability. Significance levels based on one-way ANOVA with Fisher’s LSD post-hoc test of pre-planned comparisons with each batch being assessed separately. **(k)** Immunoblot for syntaxin-6 levels and β-Actin in CAD5 cells which had either been stably transfected with a vector encoding the *Stx6* open reading frame (*Stx6* OE) or a vector encoding an unrelated sequence (NEG OE). **(l)** Quantification of syntaxin-6 band intensity normalised to the β-Actin loading control followed by normalisation to the corresponding negative control line. Bars show mean ± SEM. Statistical differences were assessed using one-way ANOVA followed by pre-planned comparisons of matched cell lines. **(m)** Growth rate measured as doubling time in days (n=3-4 biological replicas). Bars and error bars: mean ± SEM. Significance levels based on one-way ANOVA with Fisher’s LSD post-hoc test of pre-planned comparisons of matched cell lines. *P < 0.05, **P < 0.01, ***P < 0.001, ****P < 0.0001.

**Extended Figure 4.**
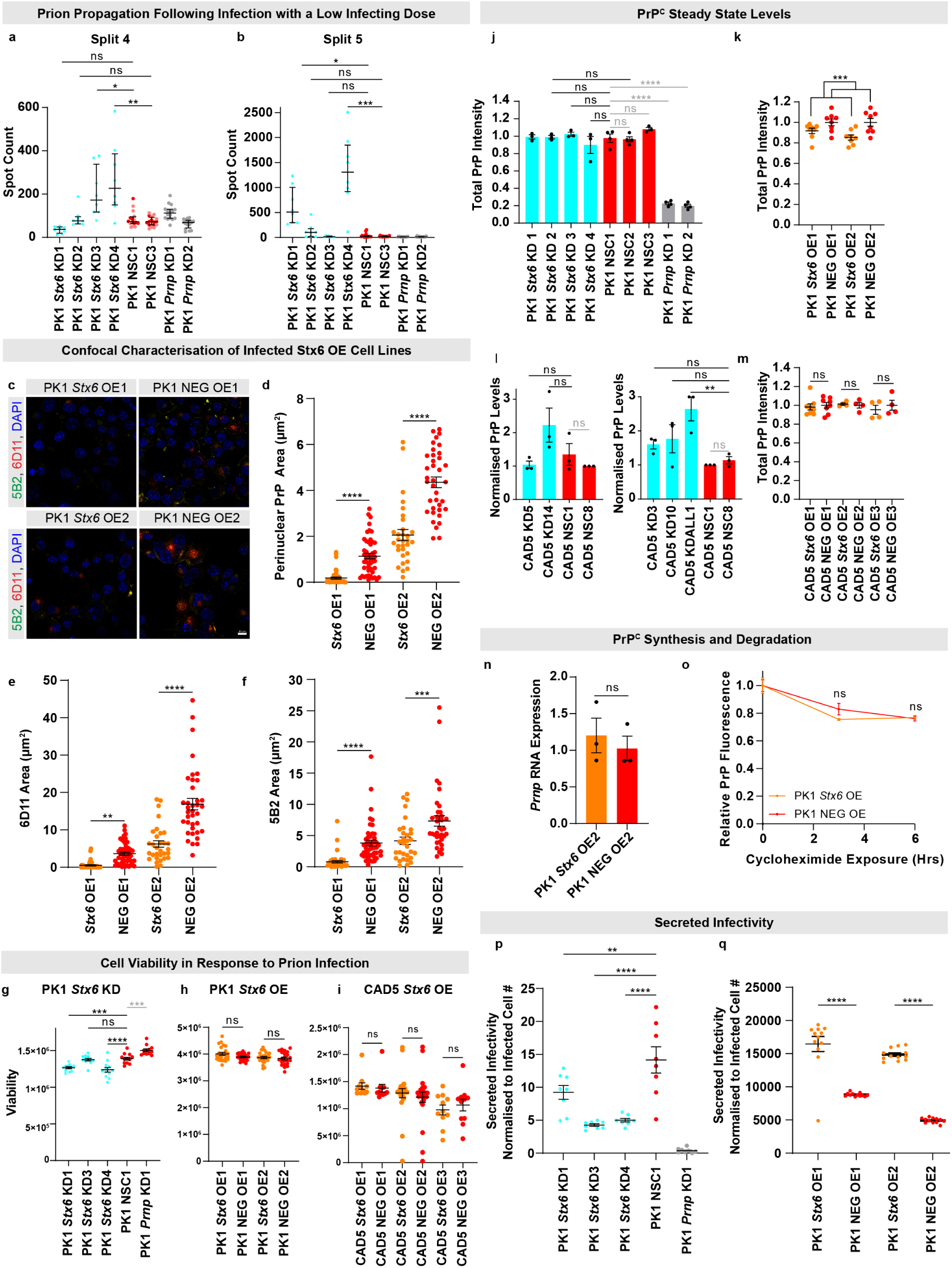
Further Characterisation of PrP-Related Phenotypes and Exploration of Explanators of the Altered Spot Count in Cell Lines with Syntaxin-6 Manipulation. **(a,b)** Representative example of the spot count of infected cell number at the 4^th^ (a) and 5th (b) split in the SCA following infection of PK1 knockdown cells with a limiting dose of RML prions, 3 x 10^-6^ (8 technical replicates/cell line). As the assay was conducted across multiple plates, spot counts of subsequent plates were normalised to the median spot count of non-silencing control (NSC1) on plate 1. NSC1 was technically replicated across two plates as indicated in separate colours. Significance levels based on Fisher’s exact test of proportions of wells with a spot count greater than an empirically determined threshold based on the background spot count with uninfected cells. **(c)** Representative images of two independent *Stx6* overexpression PK1 cell lines and corresponding negative control PK1 cell lines (NEG) stained with the discriminatory anti-PrP antibody pair, 5B2 and 6D11, assessed by laser-scanning microscopy. DAPI, nuclear stain, blue; 6D11, red; 5B2, green. Scale bar, 10 μm. **(d-f)** Quantification of the 6D11 signal area in the perinuclear region (d), total 6D11 staining area (e) and total 5B2 staining area (f) normalised by total cell count as indicated by DAPI. Statistical differences were assessed by one-way ANOVA followed by Fisher’s LSD test on planned comparisons. Each dot represents an individual image (n=31-48/cell line). **(g-i)** Assessment of cell viability at the 3^rd^ split after prion infection with infected exosomes (n=12 technical replicates/cell line) in PK1 *Stx6* knockdown (g), PK1 *Stx6* overexpression (h), and CAD5 *Stx6* overexpression cell lines (i). Significance levels based on one-way ANOVA with Fisher’s LSD post-hoc test of pre-planned comparisons. **(j-m)** Assessment of total PrP levels in the different cell lines either by flow cytometry (n=3-7/cell line) (j, k, m) or quantitative western blotting (n=3/cell line) (l). For statistical analysis, an average of the negative control lines was calculated, which was subsequently used to normalize the other values. For PK1 *Stx6* overexpression cell lines, the data was then analysed using a 2-way ANOVA approach, with syntaxin-6 expression level and cell line as the treatment factors. This was followed by pre-planned comparisons of the paired cell lines. For the other cell lines, one-way ANOVA was conducted followed by Fisher’s LSD test. **(n)** *Prnp* RNA levels as determined by RT-qPCR in one representative paired cell line. Statistical differences were assessed by an unpaired t-test. **(o)** Cells were treated with 100 µg/mL cycloheximide for the indicated time points (3 replicas/cell line/time point) with total PrP levels subsequently being determined by fluorescence intensity measured by flow cytometry. The initial fluorescence of each cell line was normalised to 1 to monitor the relative protein decay over time. Data represent the mean ± SEM. Statistical differences were assessed by two-way ANOVA followed by Fisher’s LSD test with time and *Stx6* expression as factors, which revealed a significant effect of time (P<0.0001) but not *Stx6* expression. **(p,q)** Conditioned media was collected from infected PK1 stable *Stx6* knockdown (p) and stable *Stx6* overexpression (q) cell lines and subsequently used to infect PK1 reporter cells followed by SCA analysis. Graphs show the spot count of reporter cells at the 4^th^ split normalised to the spot count of the cells the conditioned media was harvested from to correct for differences in baseline cell-associated infectivity. Statistical differences were assessed by one-way ANOVA followed by Fisher’s LSD test. *P < 0.05, **P < 0.01, ***P < 0.001, ****P< 0.0001.

**Extended Figure 5.**
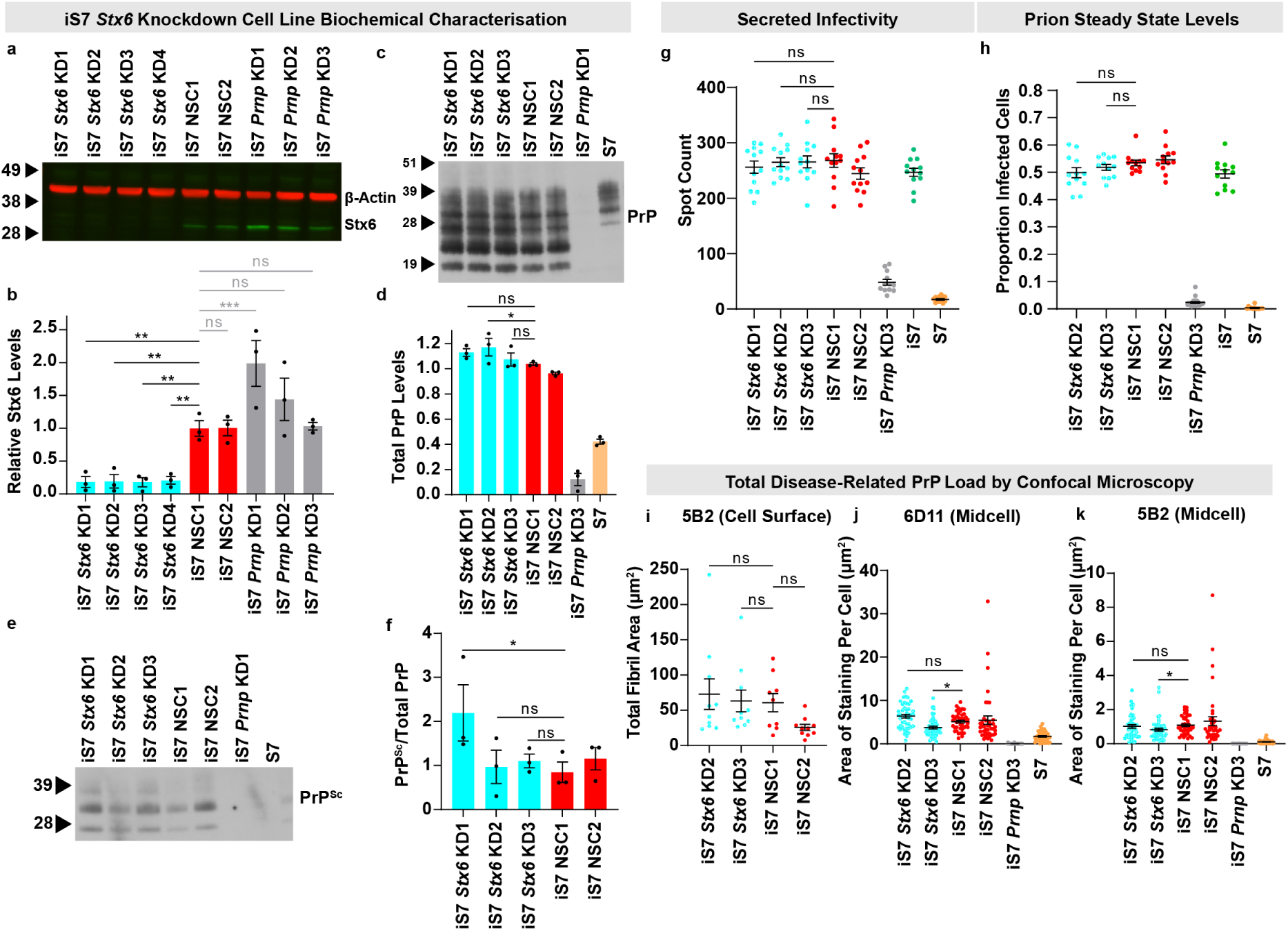
Further Characterisation of Chronically Infected PK1 Cells with Syntaxin-6 Manipulation. **(a)** Immunoblot probing for syntaxin-6 levels (bottom, green) and β-Actin (top, red) in chronically infected cells (iS7), which are either stably expressing shRNAs targeting *Stx6* or *Prnp* or shRNAs containing a non-silencing control scrambled sequence (NSC1-2). **(b)** Quantification of the syntaxin-6 band intensity relative to the β-Actin loading control followed by normalisation to the average normalised signal of the NSC cell lines. Bars show mean ± SEM. Each dot represents a different biological replicate with cells being harvested at three different passage numbers to confirm stable knockdown. Statistical differences were assessed using one-way ANOVA followed by Fisher’s LSD test assessing differences of all cell lines to NSC1. **(c)** Total PrP load in iS7 cells with *Stx6* manipulation as assessed with immunoblotting with the anti-PrP antibody, 6D11. S7 cells are uninfected cells used as a control. **(d)** Corresponding quantification with total PrP levels being normalised to an average of the NSC controls with statistics performed as described above (n=3/cell line). **(e)** Representative immunoblot showing PrP^Sc^ levels following digestion with proteinase K (PK). **(f)** Quantification of PK-resistant PrP^Sc^ levels, which were first normalised to total PrP levels before normalisation to an average of the NSC controls on each gel with statistics performed as described above. **(g)** Secreted infectivity was assessed by applying conditioned media from iS7 cell lines with stable *Stx6* manipulation to reporter PK1 cells that were subsequently assessed in the SCA. Graph shows the spot count at split 3 with the line representing the mean ± SEM (individual dots represent 12 technical replicates). Statistical differences were assessed by one-way ANOVA followed by Fisher’s LSD test on planned comparisons. **(h)** Prion steady state levels in iS7 cell lines were assessed in the SCA. Graph shows the spot count normalised to the hematoxylin total cell count at split 3 to provide a proportion of infected cells. Line represents mean ± SEM (individual dots represent the 12 technical replicates). Statistical differences were assessed by one-way ANOVA followed by Fisher’s LSD test on planned comparisons. **(i)** Quantification of the total fluorescence area of the elongated disease-related PrP aggregates from the maximum intensity projections at plasma membrane level. Statistical differences were assessed by one-way ANOVA followed by Fisher’s LSD test on planned comparisons (9-10 images/cell line). **(j, k)** Quantification of total fluorescence area of 6D11 and 5B2 staining normalised to cell number/image by confocal microscopy. Line represents mean ± SEM and individual dots indicating results from a single image (41-49 images/cell line with an average of 43 cells/image). Statistical differences were assessed by one-way ANOVA followed by Fisher’s LSD test on planned comparisons post log transformation. *p < 0.05, **p < 0.01, ***p<0.001, ****p<0.0001.

**Extended Figure 6.**
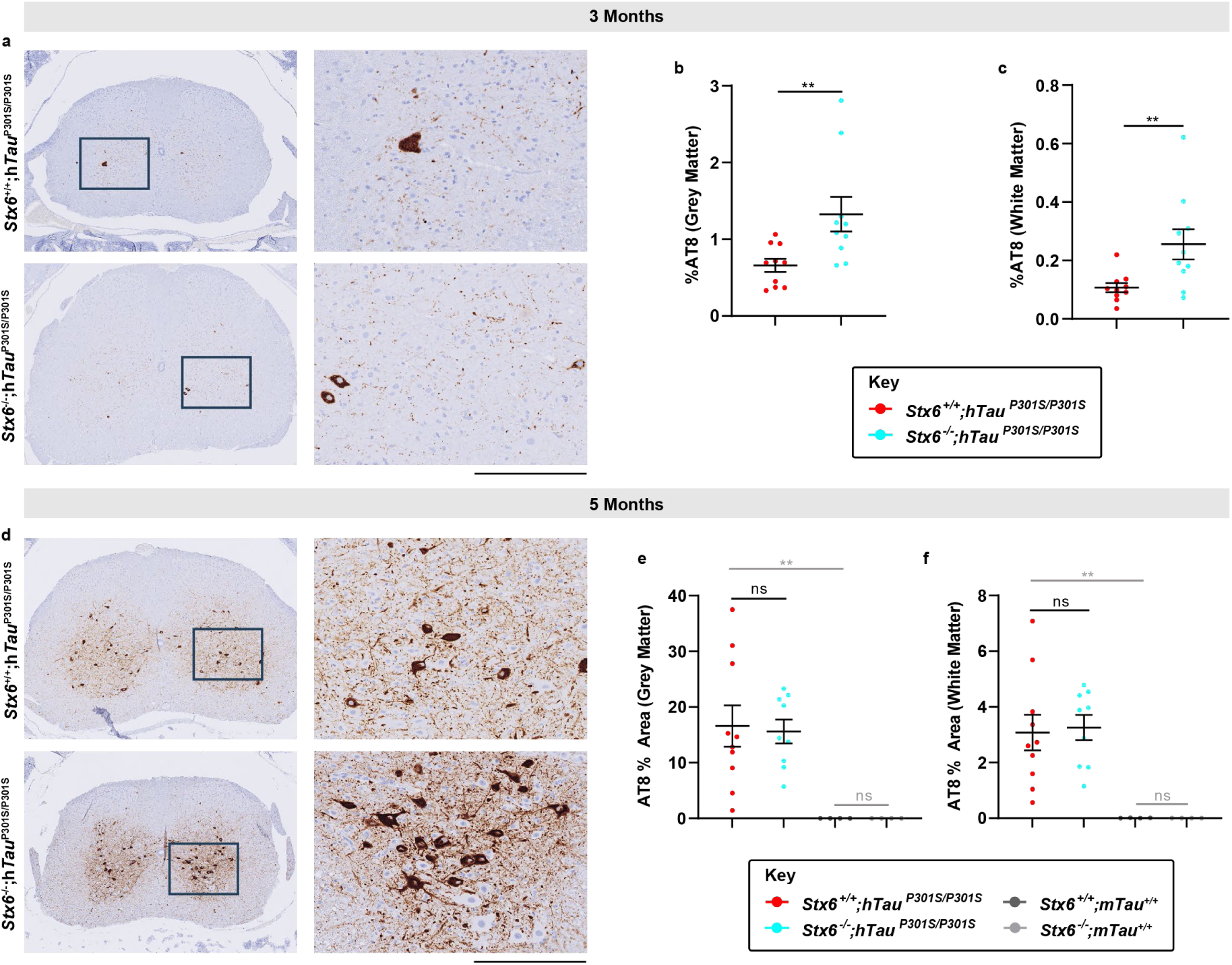
Increased AT8-Positive Tau in the Spinal Cord of *Stx6*^-/-^;h*Tau*^P301S/P301S^ Mice Relative *Stx6*^+/+^;h*Tau*^P301S/P301S^ Mice at Early Stages of the Disease. Spinal cords at the thoracic level of tauopathy mice were stained with an antibody detecting the pSer202/pThr205 phospho-epitope of tau (AT8) at 3 and 5 months of age. **(a, d)** Representative images of staining in *Stx6*^+/+^;h*Tau*^P301S/P301S^ and *Stx6*^-/-^;h*Tau*^P301S/P301S^ mice at 3 months (a) and 5 months (d). Scale bar, 800 µm (overview), 200 µm (zoom). **(b,c,e,f)** Corresponding quantification of AT8 staining density in the grey matter (b,e, H-shape or butterfly pattern) and the white matter (c,f, outer ring) (n=10/ main experimental arm). A 2-way repeated measures mixed model approach was used for statistical analysis using the unstructured covariance structure to model the within-subject correlations, with genotype as the treatment factor, brain region as the repeated factor and sex as a blocking factor. This was followed by planned comparisons on the predicted means to compare the effect of genotype on staining in the different regions. This analysis was performed post-rank transformation for the 3-month data. Two animals were excluded due to anatomical deviations. Data represent means ± SEM. **P < 0.01.

**Extended Figure 7.**
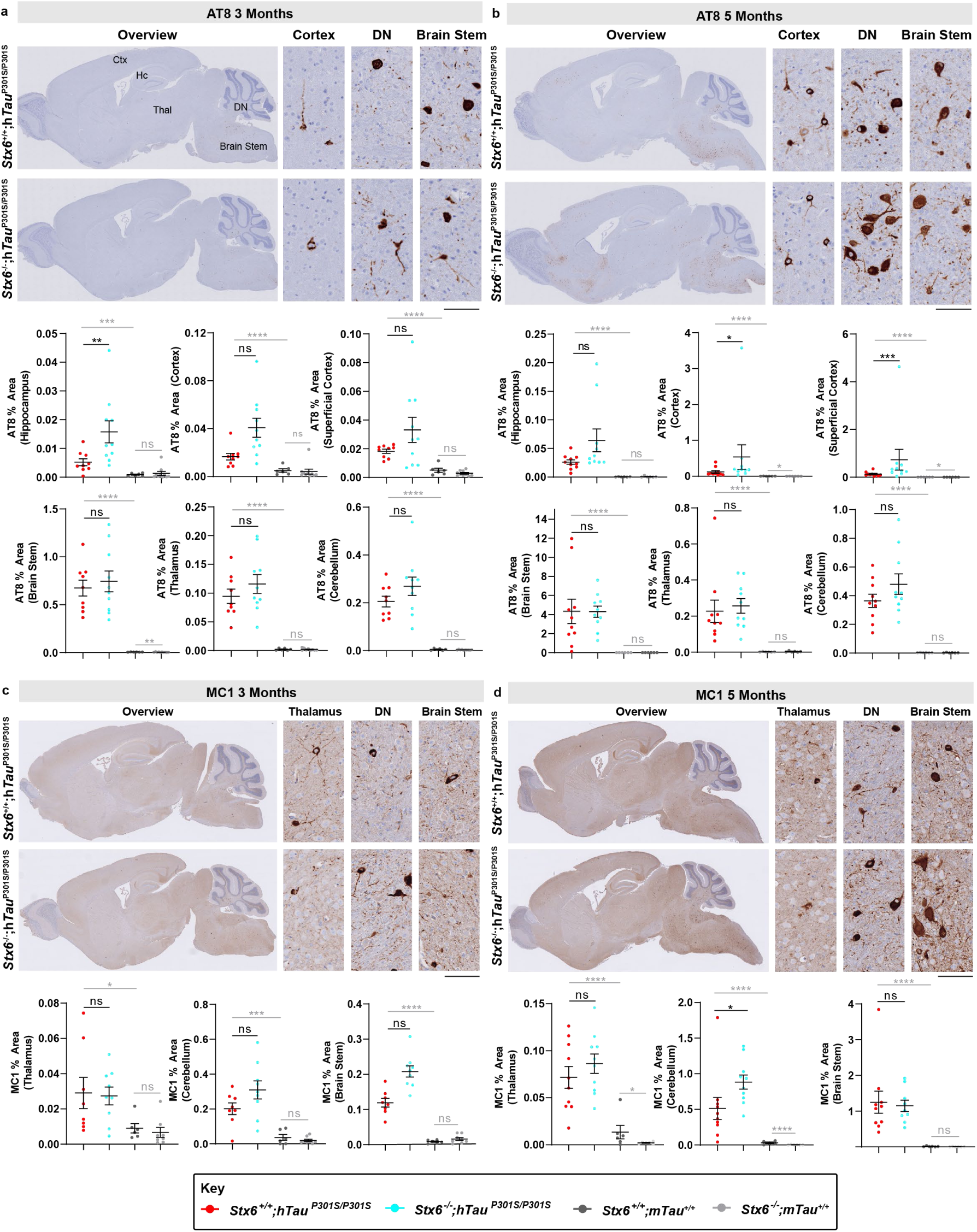
Immunohistochemical Assessment of AT8-Positive and MC1-Positive Tau Reveals Distribution Differences in *Stx6*^-/-^;h*Tau*^P301S/P301S^ Mice Relative to *Stx6*^+/+^;h*Tau*^P301S/P301S^ Mice. **(a, b)** Brain sections of tauopathy mice were stained with an antibody detecting the pSer202/pThr205 phospho-epitope of tau (AT8) at 3 months of age (a) or 5 months of age (b). Whole brain sections (left) as well as representative images from the cortex, the dentate nucleus (DN) of the cerebellum and brain stem are shown. Quantification of AT8 staining density is indicated below (n=10/main experimental arm). Following rank transformation, a 2-way repeated measures mixed model approach was used for statistical analysis using the unstructured covariance structure to model the within-subject correlations, with genotype as the treatment factor, brain region as the repeated factor and sex as a blocking factor. This was followed by planned comparisons on the predicted means to compare the effect of genotype on staining in the different brain regions. All brain regions assessed are shown as all had a disease-associated increase in staining. **(c, d)** Brain sections of tauopathy mice were stained with a tau conformational antibody (MC1) at 3 months of age (c) or 5 months of age (d). Whole brain sections (left) as well as representative images from the thalamus, the DN and the brain stem are shown. Quantification of MC1 staining density is indicated below (n=10/main experimental arm). Following rank transformation, a 2-way repeated measures mixed model approach was used for statistical analysis using the unstructured covariance structure to model the within-subject correlations, with genotype as the treatment factor, brain region as the repeated factor and sex as a blocking factor. This was followed by planned comparisons on the predicted means to compare the effect of genotype on staining in the different brain regions. The three brain regions with a disease-associated increase in staining at 3 months are shown. At 5 months, there was also a disease-associated increase in staining in the cortex and hippocampus but there were no differences between *Stx6*^+/+^;h*Tau*^P301S/P301S^ and *Stx6*^-/-^;h*Tau*^P301S/P301S^ mice. Data represent means ± SEM. Scale bar, 2mm (overview), 100 µm (high magnification). *P < 0.05, **P < 0.01, ***P < 0.001, ****P < 0.0001.

**Extended Figure 8.**
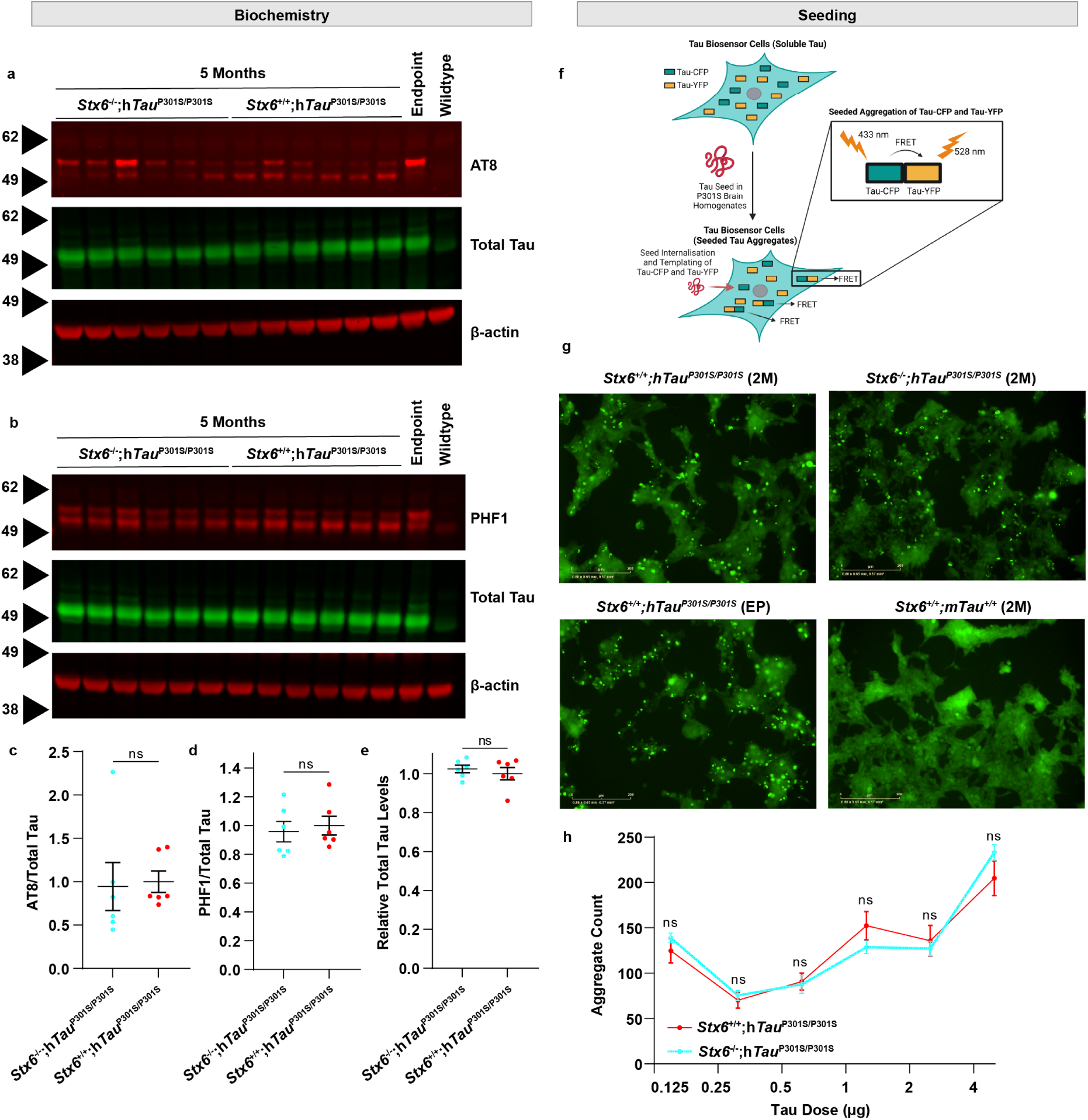
No Differences in the Biochemical and Cell-Based Assessment of Total Pathological Tau Species in *Stx6*^+/+^;h*Tau*^P301S/P301S^ and *Stx6*^-/-^;h*Tau*^P301S/P301S^ Mice. **(a)** Immunoblot of total brain homogenates of 5-month-old animals using an antibody detecting the pSer202/pThr205 phospho-epitope of tau (AT8, top), an antibody against total tau (K9JA, middle) and β-actin as a loading control (bottom). **(b)** Immunoblot of brain homogenates of 5-month-old animals using an antibody detecting Ser396/404 (PHF1), an antibody against total tau (K9JA, middle) and β-actin loading control (bottom). **(c)** Quantification of the AT8 signal intensity normalised to the β-Actin loading control corrected total tau band intensity. This was finally normalised to the average intensity values of the *Stx6^+/+^;*h*Tau*^P301S/P301S^ mice. Line and error bars: mean ± SEM with each dot representing an individual mouse. Significance levels based on results of an unpaired t-test (n=6/genotype). **(d)** Quantification of PHF1 intensity with normalisation and statistics as described above (n=6/genotype). **(e)** Quantification of total tau (n=6 animals/genotype, with each dot representing an average of 3 technical replicates across gels. No differences were also found at 3 months or endpoint (data not shown). **(f)** Schematic of tau biosensor cells employed in this study to assess for differences in seeded tau aggregation following the treatment of brain homogenates prepared from tauopathy mice with or without syntaxin-6. Created in BioRender. One, S. (2025). https://BioRender.com/i89e413. **(g)** Example images acquired in the 488 channel on the IncuCyte live cell imaging platform 72 hours after tau biosensor cells were treated with 10% (w/v) brain homogenates (5 µg total protein/well) prepared from *Stx6^+/+^;*h*Tau^P301S/P310S^ and Stx6^-/-^;*h*Tau^P301S/P301S^* mice at 2 months (2M) of age. As a positive control, cells were also treated with *Stx6*^+/+^;h*Tau*^P301S/P301S^ brain homogenate prepared from a mouse at clinical endpoint (EP). As a negative control, cells were treated with wildtype mouse brain homogenate, which was used to set up a sensitive aggregate mask detecting the green, punctate tau aggregates but not the diffuse background staining. Scale bar, 200 µm. **(h)** A titration series of brain homogenates prepared from 2-month-old *Stx6*^+/+^;h*Tau*^P301S/P301S^ and *Stx6*^-/-^;h*Tau*^P301S/P301S^ mice was applied to tau biosensor cells. Graph shows aggregate count 48 hrs post-treatment using the Incucyte live cell imager. Statistics refer to the results of two-way ANOVA with tau dose and genotype as factors, followed by Fisher’s LSD test. There was a significant effect of tau dose (P < 0.0001) but no effect of genotype.

**Extended Figure 9.**
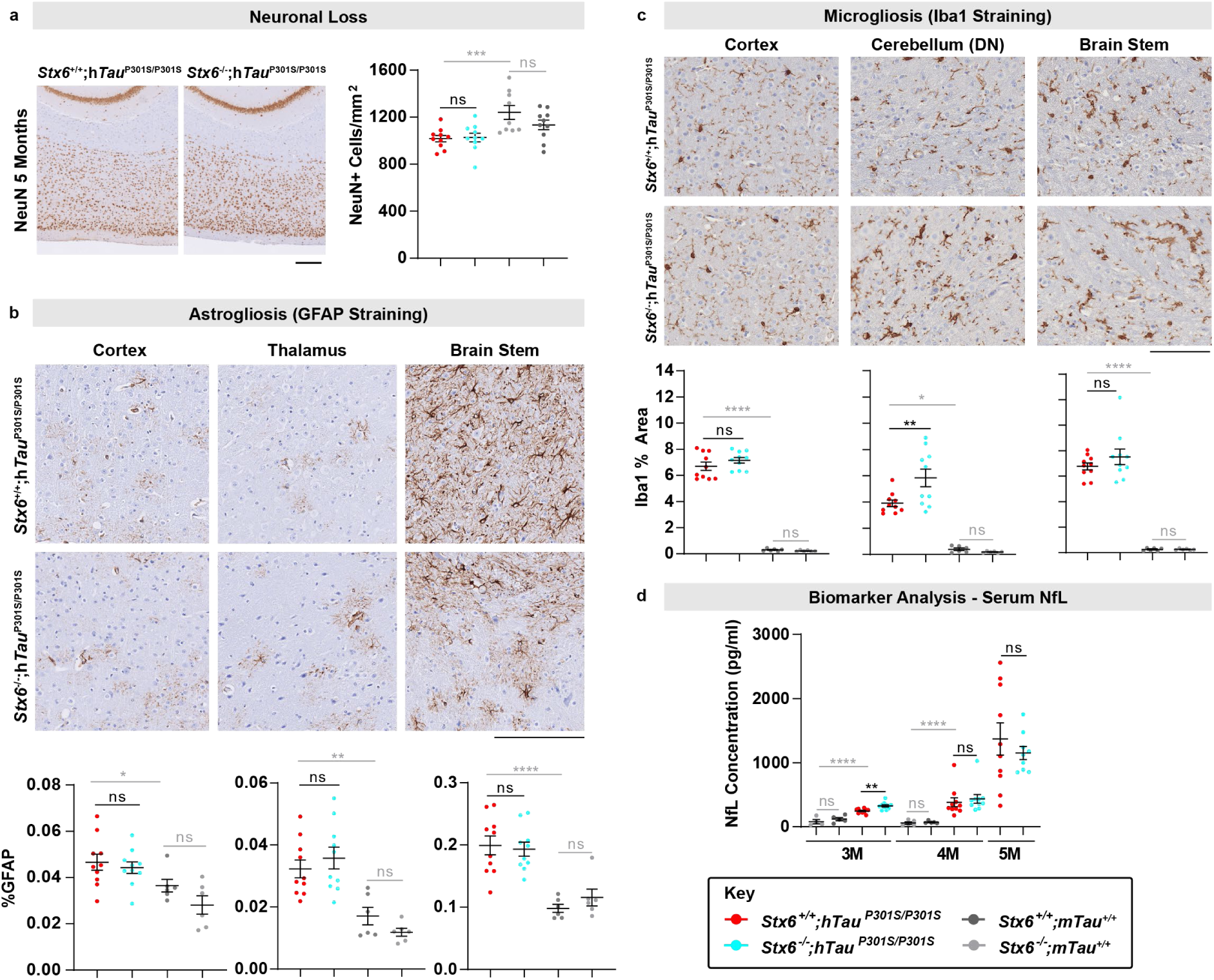
Assessment of Other Potential Correlative Readouts Underlying the Functional Rescue in *Stx6*^-/-^;h*Tau*^P301S/P301S^ Mice. **(a)** Representative images of NeuN staining in the superficial cortex (500 μm deep from the cortical surface) in *Stx6*^+/+^;h*Tau*^P301S/P301S^ and *Stx6^-^*^/-^;h*Tau*^P301S/P301S^ mice (n=10/genotype for main experimental comparison, mixed sex) at 5 months (left). Scale bar represents 250 μm. Quantification of image-based threshold analysis of NeuN staining in the superficial cortex (right). One-way ANOVA was performed with genotype as the treatment factor and sex as a blocking factor, followed by Fisher’s LSD post-hoc test of pre-planned comparisons. **(b)** Astrogliosis was assessed by staining brain sections from 5-month-old mice with an anti-GFAP antibody. Representative images from the cortex, thalamus and brain stem (regions with a disease-associated increase in staining) are shown with quantification indicated below (n=10/genotype). Following rank transformation, a 2-way repeated measures mixed model approach was used for statistical analysis using the unstructured covariance structure to model the within-subject correlations, with genotype as the treatment factor, brain region as the repeated factor and sex as a blocking factor. This was followed by planned comparisons on the predicted means to compare the effect of genotype on staining in the different brain regions. Brain regions with a disease-associated increase in GFAP staining are shown. Scale bar, 500 µm. **(c)** Microgliosis was assessed by staining brain sections from 5-month-old mice with an anti-Iba1 antibody. Representative images from the cortex, the dentate nucleus (DN) region of the cerebellum and brain stem are shown with quantification indicated below (n=10/genotype). Following rank transformation, a 2-way repeated measures mixed model approach was used for statistical analysis using the unstructured covariance structure to model the within-subject correlations, with genotype as the treatment factor, brain region as the repeated factor and sex as a blocking factor. This was followed by planned comparisons on the predicted means to compare the effect of genotype on staining in the different brain regions. Scale bar, 100 µm. Data in graphs represent means ± SEM. The hippocampus and thalamus also showed a disease-associated increase in staining in tauopathy mice, with there being no effect of syntaxin-6 knockout (data not shown). **(d)** Serum neurofilament light chain (NfL) levels were measured at 3, 4 and 5 months (n=5-10/genotype). The data were analysed within each month using a one-way ANOVA approach, with genotype as the treatment factor and sex as a blocking factor followed by Fisher’s LSD post-hoc test of pre-planned comparisons. Statistical analysis was performed on rank transformed data at 4 months to meet the ANOVA assumptions. *P < 0.05, **P < 0.01, ***P < 0.001, ****P < 0.0001.

## Supplementary Tables

**Supplementary Table 1. Transcriptomic Analysis of Uninfected *Stx6*^+/+^ and *Stx6*^-/-^ Mouse Brain to Explore Compensatory Mechanisms.** Table showing the top 50 STRING interactions for syntaxin-6 and the average normalised read counts for *Stx6*^+/+^ (WT, n=4) and *Stx6*^-/-^ (KO, n=5) mice, as well as the associated fold changes and p-values.

Excel file uploaded separately.

**Supplementary Table 2.**
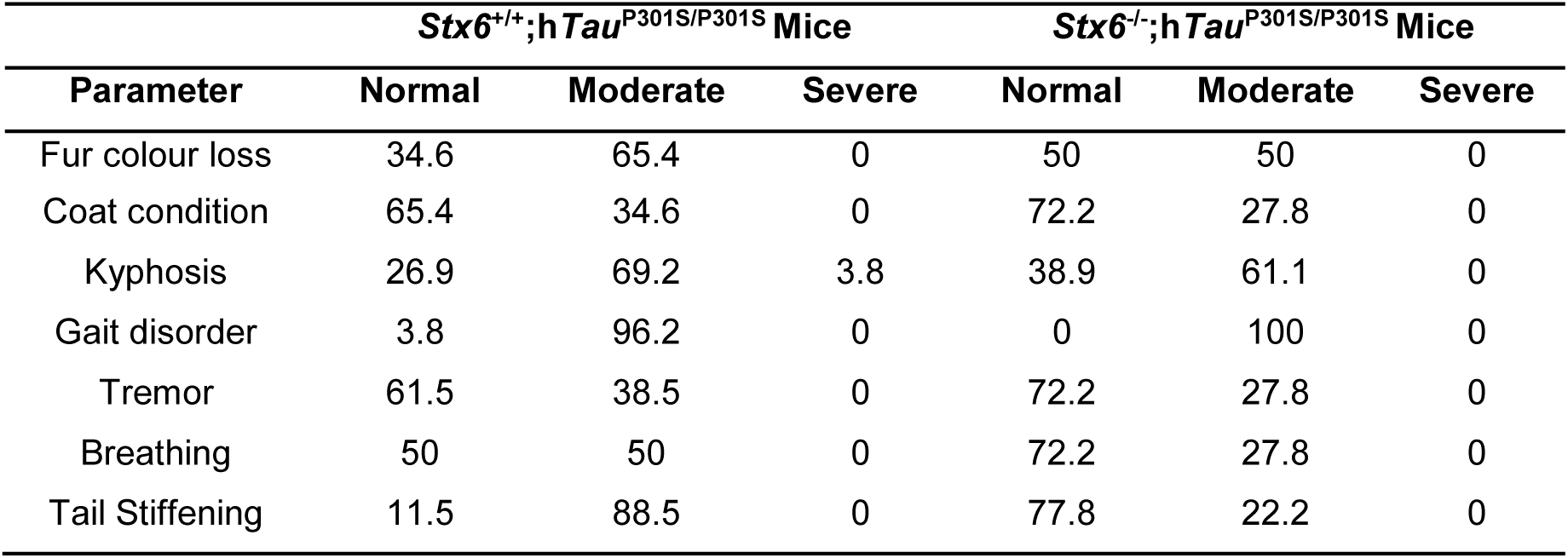
Percentage Distribution of Frailty Scores in *Stx6*^+/+^;h*Tau*^P301S/P301S^ and *Stx6*^-/-^;h*Tau*^P301S/P301S^ Mice. Table showing disease-related frailty parameters and the percentage of animals of each genotype that were recorded to have no deficit (score = 0), a mild deficit (score = 0.5) or a severe deficit (score = 1).

**Supplementary Table 3.** Time to First Symptom and Time to Culling in *Stx6*^+/+^;h*Tau*^P301S/P301S^ and *Stx6*^-/-^;h*Tau*^P301S/P301S^ Mice. Results of the Kaplan-Meier survival analysis and log rank test assessing differences in time to humane culling or time to first symptom based on *Stx6* expression level. Median time (days) is detailed with corresponding 95% confidence intervals (CI).

|  | Genotype | N start | N events | Median (days) | Lower 95% CI | Upper 95% CI | Difference (days) | P-Value Log Rank Test |
| --- | --- | --- | --- | --- | --- | --- | --- | --- |
| Time to Cull | *Stx6*<sup>+/+</sup>;hTau<sup>P301S/P301S</sup> | 20 | 20 | 211.5 | 208 | 225 | -3 | 0.258 |
|  | *Stx6*<sup>-/-</sup>;hTau<sup>P301S/P301S</sup> | 20 | 19 | 203 | 198 | 222 |  |  |
| First Symptom | *Stx6*<sup>+/+</sup>;hTau<sup>P301S/P301S</sup> | 20 | 20 | 172 | 172 | 175 | +3 | 0.154 |
|  | *Stx6*<sup>-/-</sup>;hTau<sup>P301S/P301S</sup> | 20 | 20 | 177 | 173 | 178 |  |  |

## References

1 Collinge, J. Mammalian prions and their wider relevance in neurodegenerative diseases. Nature 539, 217–226, doi:10.1038/nature20415 (2016).

2 Jucker, M. & Walker, L. C. Propagation and spread of pathogenic protein assemblies in neurodegenerative diseases. Nat Neurosci 21, 1341–1349, doi:10.1038/s41593-018-0238-6 (2018).

3 Purro, S. A. et al. Transmission of amyloid-β protein pathology from cadaveric pituitary growth hormone. Nature 564, 415–419, doi:10.1038/s41586-018-0790-y (2018).

4 Banerjee, G. et al. Iatrogenic Alzheimer’s disease in recipients of cadaveric pituitary-derived growth hormone. Nature Medicine 30, 394–402, doi:10.1038/s41591-023-02729-2 (2024).

5 Frost, B., Jacks, R. L. & Diamond, M. I. Propagation of tau misfolding from the outside to the inside of a cell. The Journal of biological chemistry 284, 12845–12852, doi:10.1074/jbc.M808759200 (2009).

6 Mirbaha, H. et al. Seed-competent tau monomer initiates pathology in a tauopathy mouse model. Journal of Biological Chemistry 298, 102163, 10.1016/j.jbc.2022.102163 (2022).

7 Guo, J. L. & Lee, V. M. Seeding of normal Tau by pathological Tau conformers drives pathogenesis of Alzheimer-like tangles. The Journal of biological chemistry 286, 15317–15331, doi:10.1074/jbc.M110.209296 (2011).

8 Clavaguera, F. et al. Transmission and spreading of tauopathy in transgenic mouse brain. Nature cell biology 11, 909–913, doi:10.1038/ncb1901 (2009).

9 Sanders, D. W. et al. Distinct tau prion strains propagate in cells and mice and define different tauopathies. Neuron 82, 1271–1288, doi:10.1016/j.neuron.2014.04.047 (2014).

10 Jones, E. et al. Identification of novel risk loci and causal insights for sporadic Creutzfeldt-Jakob disease: a genome-wide association study. The Lancet. Neurology 19, 840–848, doi:10.1016/s1474-4422(20)30273-8 (2020).

11 Höglinger, G. U. et al. Identification of common variants influencing risk of the tauopathy progressive supranuclear palsy. Nature Genetics 43, 699–705, doi:10.1038/ng.859 (2011).

12 Ferrari, R. et al. Assessment of common variability and expression quantitative trait loci for genome-wide associations for progressive supranuclear palsy. Neurobiology of Aging 35, 1514.e1511–1514.e1511, doi:10.1016/j.neurobiolaging.2014.01.010 (2014).

13 Chen, J. A. et al. Joint genome-wide association study of progressive supranuclear palsy identifies novel susceptibility loci and genetic correlation to neurodegenerative diseases. Molecular Neurodegeneration 13, doi:10.1186/s13024-018-0270-8 (2018).

14 Chen, Z. et al. Genome-wide survey of copy number variants finds MAPT duplications in progressive supranuclear palsy. Movement disorders : official journal of the Movement Disorder Society 34, 1049–1059, doi:10.1002/mds.27702 (2019).

15 Farrell, K. et al. Genetic, transcriptomic, histological, and biochemical analysis of progressive supranuclear palsy implicates glial activation and novel risk genes. Nature Communications 15, 7880, doi:10.1038/s41467-024-52025-x (2024).

16 Wingo, A. P. et al. Integrating human brain proteomes with genome-wide association data implicates new proteins in Alzheimer’s disease pathogenesis. Nature Genetics 53, 143–146, doi:10.1038/s41588-020-00773-z (2021).

17 Jiang, D., Nan, H., Chen, Z., Zou, W.-Q. & Wu, L. Genetic insights into drug targets for sporadic Creutzfeldt-Jakob disease: Integrative multi-omics analysis. Neurobiology of disease, 106599, 10.1016/j.nbd.2024.106599 (2024).

18 Küçükali, F. et al. Multiomic analyses direct hypotheses for Creutzfeldt-Jakob disease risk genes. Brain, awaf032, doi:10.1093/brain/awaf032 (2025).

19 Bock, J. B., Klumperman, J., Davanger, S. & Scheller, R. H. Syntaxin 6 functions in trans-Golgi network vesicle trafficking. Molecular biology of the cell 8, 1261–1271, doi:10.1091/mbc.8.7.1261 (1997).

20 Bock, J. B., Lin, R. C. & Scheller, R. H. A new syntaxin family member implicated in targeting of intracellular transport vesicles. The Journal of biological chemistry 271, 17961–17965, doi:10.1074/jbc.271.30.17961 (1996).

21 Wendler, F. & Tooze, S. Syntaxin 6: the promiscuous behaviour of a SNARE protein. *Traffic (Copenhagen*, Denmark) 2, 606–611, doi:10.1034/j.1600-0854.2001.20903.x (2001).

22 Bryois, J. et al. Cell-type-specific cis-eQTLs in eight human brain cell types identify novel risk genes for psychiatric and neurological disorders. Nature Neuroscience 25, 1104–1112, doi:10.1038/s41593-022-01128-z (2022).

23 Jones, E. et al. Characterisation and prion transmission study in mice with genetic reduction of sporadic Creutzfeldt-Jakob disease risk gene Stx6. Neurobiology of disease 190, 106363, 10.1016/j.nbd.2023.106363 (2024).

24 Sandberg, M. K., Al-Doujaily, H., Sharps, B., Clarke, A. R. & Collinge, J. Prion propagation and toxicity in vivo occur in two distinct mechanistic phases. Nature 470, 540–542, doi:10.1038/nature09768 (2011).

25 Sandberg, M. K. et al. Prion neuropathology follows the accumulation of alternate prion protein isoforms after infective titre has peaked. Nature Communications 5, 4347, doi:10.1038/ncomms5347 (2014).

26 Holmes, B. B. et al. Proteopathic tau seeding predicts tauopathy in vivo. Proceedings of the National Academy of Sciences 111, E4376, doi:10.1073/pnas.1411649111 (2014).

27 Santacruz, K. et al. Tau suppression in a neurodegenerative mouse model improves memory function. *Science (New York*, N.Y.) 309, 476–481, doi:10.1126/science.1113694 (2005).

28 Spires, T. L. et al. Region-specific dissociation of neuronal loss and neurofibrillary pathology in a mouse model of tauopathy. Am J Pathol 168, 1598–1607, doi:10.2353/ajpath.2006.050840 (2006).

29 Le Corre, S. et al. An inhibitor of tau hyperphosphorylation prevents severe motor impairments in tau transgenic mice. Proceedings of the National Academy of Sciences of the United States of America 103, 9673–9678, doi:10.1073/pnas.0602913103 (2006).

30 Andorfer, C. et al. Cell-cycle reentry and cell death in transgenic mice expressing nonmutant human tau isoforms. The Journal of neuroscience : the official journal of the Society for Neuroscience 25, 5446–5454, doi:10.1523/jneurosci.4637-04.2005 (2005).

31 Udeochu, J. C. et al. Tau activation of microglial cGAS-IFN reduces MEF2C-mediated cognitive resilience. Nat Neurosci 26, 737–750, doi:10.1038/s41593-023-01315-6 (2023).

32 Apicco, D. J. et al. Reducing the RNA binding protein TIA1 protects against tau-mediated neurodegeneration in vivo. Nat Neurosci 21, 72–80, doi:10.1038/s41593-017-0022-z (2018).

33 Wegmann, S. et al. Removing endogenous tau does not prevent tau propagation yet reduces its neurotoxicity. The EMBO journal 34, 3028–3041, 10.15252/embj.201592748 (2015).

34 Allen, B. et al. Abundant tau filaments and nonapoptotic neurodegeneration in transgenic mice expressing human P301S tau protein. The Journal of neuroscience : the official journal of the Society for Neuroscience 22, 9340–9351, doi:10.1523/JNEUROSCI.22-21-09340.2002 (2002).

35 Chandler, R. L. Encephalopathy in mice produced by inoculation with scrapie brain material. Lancet 1, 1378–1379, doi:10.1016/s0140-6736(61)92008-6 (1961).

36 Klöhn, P. C., Stoltze, L., Flechsig, E., Enari, M. & Weissmann, C. A quantitative, highly sensitive cell-based infectivity assay for mouse scrapie prions. Proceedings of the National Academy of Sciences of the United States of America 100, 11666–11671, doi:10.1073/pnas.1834432100 (2003).

37 Schmidt, C. et al. A systematic investigation of production of synthetic prions from recombinant prion protein. Open Biology 5 (2015).

38 Benilova, I. et al. Highly infectious prions are not directly neurotoxic. Proceedings of the National Academy of Sciences 117, 23815, doi:10.1073/pnas.2007406117 (2020).

39 Minikel, E. V. et al. Prion protein lowering is a disease-modifying therapy across prion disease stages, strains and endpoints. Nucleic Acids Research 48, 10615–10631, doi:10.1093/nar/gkaa616 (2020).

40 Qi, Y., Wang, J. K., McMillian, M. & Chikaraishi, D. M. Characterization of a CNS cell line, CAD, in which morphological differentiation is initiated by serum deprivation. The Journal of neuroscience : the official journal of the Society for Neuroscience 17, 1217–1225, doi:10.1523/jneurosci.17-04-01217.1997 (1997).

41 Bhamra, S. et al. Prion propagation is dependent on key amino acids in Charge cluster 2 within the prion protein. bioRxiv, 2022.2008.2008.503133, doi:10.1101/2022.08.08.503133 (2022).

42 Enari, M., Flechsig, E. & Weissmann, C. Scrapie prion protein accumulation by scrapie-infected neuroblastoma cells abrogated by exposure to a prion protein antibody. Proceedings of the National Academy of Sciences 98, 9295–9299, doi:10.1073/pnas.151242598 (2001).

43 Vorberg, I., Raines, A., Story, B. & Priola, S. A. Susceptibility of common fibroblast cell lines to transmissible spongiform encephalopathy agents. J Infect Dis 189, 431–439, doi:10.1086/381166 (2004).

44 Marbiah, M. M. et al. Identification of a gene regulatory network associated with prion replication. The EMBO journal 33, 1527–1547, doi:10.15252/embj.201387150 (2014).

45 Ribes, J. M. et al. Prion protein conversion at two distinct cellular sites precedes fibrillisation. Nature Communications 14, 8354, doi:10.1038/s41467-023-43961-1 (2023).

46 Scattoni, M. L. et al. Early behavioural markers of disease in P301S tau transgenic mice. Behav Brain Res 208, 250–257, doi:10.1016/j.bbr.2009.12.002 (2010).

47 Sukoff Rizzo, S. J., et al. Assessing Healthspan and Lifespan Measures in Aging Mice: Optimization of Testing Protocols, Replicability, and Rater Reliability. Current Protocols in Mouse Biology 8, e45, 10.1002/cpmo.45 (2018).

48 Hampton, D. W. et al. Cell-mediated neuroprotection in a mouse model of human tauopathy. The Journal of neuroscience : the official journal of the Society for Neuroscience 30, 9973–9983, doi:10.1523/jneurosci.0834-10.2010 (2010).

49 Vogler, L. et al. Assessment of synaptic loss in mouse models of β-amyloid and tau pathology using [(18)F]UCB-H PET imaging. Neuroimage Clin 39, 103484, doi:10.1016/j.nicl.2023.103484 (2023).

50 Hummerich, H. et al. Genome wide association study of clinical duration and age at onset of sporadic CJD. PloS one 19, e0304528, doi:10.1371/journal.pone.0304528 (2024).

51 Mortberg, M. A., Vallabh, S. M. & Minikel, E. V. Disease stages and therapeutic hypotheses in two decades of neurodegenerative disease clinical trials. Scientific Reports 12, 17708, doi:10.1038/s41598-022-21820-1 (2022).

52 Leonenko, G. et al. Genetic risk for alzheimer disease is distinct from genetic risk for amyloid deposition. Annals of neurology 86, 427–435, doi:10.1002/ana.25530 (2019).

53 Liu, G. et al. Genome-wide survival study identifies a novel synaptic locus and polygenic score for cognitive progression in Parkinson’s disease. Nat Genet 53, 787–793, doi:10.1038/s41588-021-00847-6 (2021).

54 Vorberg, I. M. All the Same? The Secret Life of Prion Strains within Their Target Cells. Viruses 11, doi:10.3390/v11040334 (2019).

55 Peak, T. C. et al. Syntaxin 6-mediated exosome secretion regulates enzalutamide resistance in prostate cancer. Molecular Carcinogenesis 59, 62–72, doi:10.1002/mc.23129 (2020).

56 Ito, S. I. & Tanaka, Y. Evaluation of LC3-II Release via Extracellular Vesicles in Relation to the Accumulation of Intracellular LC3-positive Vesicles. J Vis Exp, doi:10.3791/67385 (2024).

57 Fevrier, B. et al. Cells release prions in association with exosomes. Proceedings of the National Academy of Sciences 101, 9683–9688, doi:10.1073/pnas.0308413101 (2004).

58 Saman, S. et al. Exosome-associated Tau Is Secreted in Tauopathy Models and Is Selectively Phosphorylated in Cerebrospinal Fluid in Early Alzheimer Disease*. Journal of Biological Chemistry 287, 3842–3849, 10.1074/jbc.M111.277061 (2012).

59 Sangar, D. et al. Syntaxin-6 delays prion protein fibril formation and prolongs presence of toxic aggregation intermediates. eLife 13, e83320, doi:10.7554/eLife.83320 (2024).

60 Heisler, F. F. et al. Muskelin Coordinates PrP(C) Lysosome versus Exosome Targeting and Impacts Prion Disease Progression. Neuron 99, 1155–1169.e1159, doi:10.1016/j.neuron.2018.08.010 (2018).

61 Alves, R. N. et al. A New Take on Prion Protein Dynamics in Cellular Trafficking. Int J Mol Sci 21, doi:10.3390/ijms21207763 (2020).

62 Cherry, P. & Gilch, S. The Role of Vesicle Trafficking Defects in the Pathogenesis of Prion and Prion-Like Disorders. Int J Mol Sci 21, doi:10.3390/ijms21197016 (2020).

63 Shim, S. Y., Karri, S., Law, S., Schatzl, H. M. & Gilch, S. Prion infection impairs lysosomal degradation capacity by interfering with rab7 membrane attachment in neuronal cells. Scientific Reports 6, 21658, doi:10.1038/srep21658 (2016).

64 Nunziante, M. et al. Proteasomal dysfunction and endoplasmic reticulum stress enhance trafficking of prion protein aggregates through the secretory pathway and increase accumulation of pathologic prion protein. The Journal of biological chemistry 286, 33942–33953, doi:10.1074/jbc.M111.272617 (2011).

65 Lee, W. S. et al. Syntaxins 6 and 8 facilitate tau into secretory pathways. Biochemical Journal 478, 1471–1484, doi:10.1042/bcj20200664 (2021).

66 Bellenguez, C. et al. New insights into the genetic etiology of Alzheimer’s disease and related dementias. Nature Genetics 54, 412–436, doi:10.1038/s41588-022-01024-z (2022).

67 Nalls, M. A. et al. Evidence for GRN connecting multiple neurodegenerative diseases. Brain Commun 3, fcab095, doi:10.1093/braincomms/fcab095 (2021).

68 Ochoa, D. et al. Human genetics evidence supports two-thirds of the 2021 FDA-approved drugs. Nat Rev Drug Discov, doi:10.1038/d41573-022-00120-3 (2022).

69 Nelson, M. R. et al. The support of human genetic evidence for approved drug indications. Nat Genet 47, 856–860, doi:10.1038/ng.3314 (2015).

70 Plenge, R. M., Scolnick, E. M. & Altshuler, D. Validating therapeutic targets through human genetics. Nat Rev Drug Discov 12, 581–594, doi:10.1038/nrd4051 (2013).

71 Minikel, E. V., Painter, J. L., Dong, C. C. & Nelson, M. R. Refining the impact of genetic evidence on clinical success. Nature 629, 624–629, doi:10.1038/s41586-024-07316-0 (2024).

72 Raymond, G. J. et al. Antisense oligonucleotides extend survival of prion-infected mice. JCI Insight 5, e131175, doi:10.1172/jci.insight.131175 (2019).

73 DeVos, S. L. et al. Tau reduction prevents neuronal loss and reverses pathological tau deposition and seeding in mice with tauopathy. Sci Transl Med 9, doi:10.1126/scitranslmed.aag0481 (2017).

74 Mummery, C. J. et al. Results of the first-in-human, randomized, double-blind, placebo-controlled phase 1b study of lumbar intrathecal bolus administrations of antisense oligonucleotide (ISIS 814907; BIIB080) targeting tau mRNA in patients with mild Alzheimer’s disease. Alzheimer’s & Dementia 17, e051871, 10.1002/alz.051871 (2021).

75 Mummery, C. J. et al. Tau-targeting antisense oligonucleotide MAPTRx in mild Alzheimer’s disease: a phase 1b, randomized, placebo-controlled trial. Nature Medicine 29, 1437–1447, doi:10.1038/s41591-023-02326-3 (2023).

76 Mead, S. et al. Prion protein monoclonal antibody (PRN100) therapy for Creutzfeldt-Jakob disease: evaluation of a first-in-human treatment programme. The Lancet. Neurology 21, 342–354, doi:10.1016/s1474-4422(22)00082-5 (2022).

77 Bugiani, O. et al. Frontotemporal dementia and corticobasal degeneration in a family with a P301S mutation in tau. Journal of neuropathology and experimental neurology 58, 667–677, doi:10.1097/00005072-199906000-00011 (1999).

78 Sperfeld, A. D. et al. FTDP-17: an early-onset phenotype with parkinsonism and epileptic seizures caused by a novel mutation. Annals of neurology 46, 708–715, doi:10.1002/1531-8249(199911)46:5<708::aid-ana5>3.0.co;2-k (1999).

79 Lossos, A. et al. Frontotemporal dementia and parkinsonism with the P301S tau gene mutation in a Jewish family. Journal of neurology 250, 733–740, doi:10.1007/s00415-003-1074-4 (2003).

80 Yasuda, M. et al. A Japanese patient with frontotemporal dementia and parkinsonism by a tau P301S mutation. Neurology 55, 1224–1227, doi:10.1212/wnl.55.8.1224 (2000).

81 Schweighauser, M. et al. Cryo-EM structures of tau filaments from the brains of mice transgenic for human mutant P301S Tau. Acta Neuropathologica Communications 11, 160, doi:10.1186/s40478-023-01658-y (2023).

82 Baptista, J. & Pike, M. C. Algorithm AS 115: Exact Two-Sided Confidence Limits for the Odds Ratio in a 2 × 2 Table. Journal of the Royal Statistical Society. Series C (Applied Statistics) 26, 214–220, doi:10.2307/2347041 (1977).

