## Supplementary material for "Intracellular Trafficking SNARE Protein, Syntaxin-6, is a Modifier of Prion and Tau Pathogenesis *in vivo* and in Cellular Models": Methods

### Online Methods

All reagents, resources and cellular and animal models used in this work are detailed in the key resources table (see **Section 1.6**). The broad statistical approach for each study is outlined but the specific ‘n’ number and any deviations in the statistical analyses are detailed for each specific experiment in the relevant figure legend. All graphing and statistical analysis were conducted in Graphpad Prism or InVivoStat.

#### 1.1 Research Governance

All experimental procedures employing mouse-adapted prions were conducted in microbiological containment level 2 (CL2) or level 3 (CL3) facilities with strict adherence to safety protocols and guidelines. Work with mice was performed under approval and license granted by the UK Home Office (Animals (Scientific Procedures) Act 1986), which conforms to UCL institutional guidelines and Animal Research: Reporting of In Vivo Experiments (ARRIVE) guidelines (www.nc3rs.org.uk/ARRIVE/).

#### 1.2 General Animal Maintenance and Husbandry

##### 1.2.1 Housing Conditions

Breeding colonies were kept in individually ventilated cages at a temperature of 20-24°C, humidity of 45-65%, an average of 75 air exchanges per hour in the cages and a 12/12-hour light/dark cycle with the lights on at 7AM. The maximum caging density was five mice starting from weaning. Lignocel or EcoPure wood fibres were provided as bedding along with ‘Bed r’ nest’ nesting materials, wood blocks and a shelter.

##### 1.2.2 Animal Care

Mice were fed a standardised irradiated mouse diet and provided with reverse osmosis drinking water *ad libitum.* All materials, including cages, lids, feeders and water bottles were washed in a cage washer and autoclaved before use. All mice were frequently checked by veterinary and animal care staff and had not undergone prior procedures. All animals otherwise received no procedures except those reported in this work. Work was conducted in a CL3 facility, with staff required to air shower upon entry and exit. External animals were only brought in from other clean-screened units and following in-house vet approval.

##### 1.2.3 Genotyping

*DNA Extraction*

DNA was extracted from ear biopsies of mice using MyTaq Extract-PCR Kit (2:1:17 ratio of buffer A: buffer B: $H_20$). This was subsequently incubated at 75°C for 30 min and then 95°C for 15 min.

*Standard PCR Genotyping*

PCR amplification was performed using 1x MyTaq HS Red Mix with a triple primer set (**Table 1**).

**Table 1. Primers Used for Standard PCR Genotyping of *Stx6*$^{+/+}$, *Stx6*$^{+/-}$ and *Stx6*$^{-/-}$ mice.**

| PRIMER | SEQUENCE (5' - 3') |
|---|---|
| ***Stx6*$^{+/+}$ and *Stx6*$^{-/-}$ Lines** | |
| Stx6_geno_F1 | CGATCTGTGAGACTCATCGGG |
| Stx6_geno_R1 | GGGAGTCCTAACACCACCTTC |
| Stx6_geno_R2 | GGACACCATGCTTTCAAGATTT |

This was amplified using a Biorad Tetrad 2 using the cycling parameters shown in **Table 2**. PCR reactions were analysed by agarose gel electrophoresis (0.5% agarose in 1x TAE with gelred run at 4.6 V/cm) with image capture performed on a Universal Hood II Gel Doc System.

**Table 2. PCR Cycling parameters for Genotyping and *Stx6*$^{+/+}$ and *Stx6*$^{-/-}$ Lines.**

| Step | Temperature (°C) | Time | Cycles |
|---|---|---|---|
| Initial Denaturation | 95 | 3 min | 1 |
| Denaturation | 95 | 15 sec | |
| Primer Annealing | 59 | 15 sec | 35 |
| Extension | 72 | 20 sec | |
| Final Extension | 72 | 5 min | 1 |

*Copy Count Genotyping*

Copy count assays were used for genotyping the humanised tauopathy mice, using the housekeeping gene *Dot1l* as an internal control (**Table 3**).

**Table 3. Copy Count Assay qPCR Set Sequences for Genotyping Humanised Tauopathy Mice**

| qPCR Set | SEQUENCE (5' - 3') |
|---|---|
| **Dot1l PrimeTime qPCR Assay** | |
| Forward Primer | GTT GGC ATC CTT ATG CTT CAT C |
| Reverse Primer | GTG TGC TAC GCC TGA AAT AAA G |
| Probe | /5SUN/TT GAG AGC T/ZEN/G GCC CTG AAT GGT C/3IABkFQ/ |
| **P301S PrimeTime qPCR Assay** | |

| | |
|---|---|
| Forward Primer | TCCTAGAATTGATCCTGGCGTAA |
| Reverse Primer | AACTGGGAGTCGCTACAGTGAGT |
| Probe | FAM-AGCGAAGAGGCCCGCACCTCA-BHQ |

In brief, 3 µL DNA was added to 1x TaqPath™ ProAmp™ Master Mix with 1x PrimeTime qPCR assays (1:2 probe:primer ratio) in duplex in a MicroAmp™ Optical 96-Well Reaction Plate (total volume: 20 µL). Reactions were run on the Quant Studio 12K FLEX machine using Quantstudio 12K Flex Software v1.3 (**Table 4**).

**Table 4. Cycling parameters for Copy Count PCR for Genotyping Humanised Tauopathy Mice**

| Step | Temperature (°C) | Time | Cycles |
|---|---|---|---|
| Initial | 95 | 10 min | 1 |
| 1 | 95 | 15 sec | 40 |
| 4 | 60 | 1 min | |

Relative expression of the mutant tau allele was calculated using double delta $C_t$ analysis with normalisation to the housekeeping gene (*Dot1l*). First, the $2^{-Ct}$ was calculated for each allele with each experimental allele subsequently being divided by the $2^{-Ct}$ value of *Dot1l* to provide a $2^{-\Delta Ct}$ value for each sample. These were then normalised to a positive control, calculated by multiplying the $2^{-\Delta Ct}$ value of each sample by a normalisation factor (calculated by divided the number of alleles by the $2^{-\Delta Ct}$ value of the positive control) to provide a $2^{-\Delta\Delta Ct}$ value used for genotype assignment.

#### 1.3 Prion Transmission Studies into Mice

##### 1.3.1 Breeding Strategy

The *Stx6*$^{+/+}$ and *Stx6*$^{-/-}$ C57BL/6N homozygous lines (derived from C57BL/6NTac-Stx6$^{em1(IMPC)H}$ mice)[1] were first intercrossed to mitigate the effects of genetic drift, with heterozygote mice subsequently being intercrossed to generate littermate controls to populate the clinical endpoint groups of the 1% (w/v) RML transmission study. Clinical endpoint animals were randomly allocated to groups based on Mendelian inheritance. For reasons of feasibility, re-established homozygous independent lines were used to populate the other RML timed cull groups as well as the RML titration study. Animals were randomised by cage when assigning to groups.

##### 1.3.2 Study Design, Power Calculations and Statistical Analyses of Prion Transmission Studies

*Serial Dilution of RML into Stx6*$^{+/+}$ *and Stx6*$^{-/-}$ *Mice*

Mice were infected with a dilution series of 10% (w/v) RML-infected brain homogenate with the attack rate being the primary outcome measure for concentrations with a <90% partial attack rate ($10^{-5}$, $10^{-6}$, $10^{-7}$, $10^{-8}$). Animals which were culled due to unrelated health concerns before the elective 400 days post-inoculation (dpi) time cull were excluded, as well as animals where there was a discrepancy between the clinical and neuropathological observations (**Table 5**). This was a female-only study to allow comparability with previous experiments. The main comparison in these studies were prion-infected *Stx6*$^{+/+}$ mice with *Stx6*$^{-/-}$ mice at each dose administered. A single animal was considered an experimental unit.

**Table 5. Study Design of RML Serial Dilution Titration Study in *Stx6*$^{+/+}$ and *Stx6*$^{-/-}$ Mice.** Dose refers to the concentration of 10% (w/v) RML-infected brain homogenate inoculated. The number of infected mice refers to the number inoculated with the number of exclusions detailed in brackets.

| Dose | # Infected *Stx6*$^{+/+}$ Mice | # Infected *Stx6*$^{-/-}$ Mice Animals |
|---|---|---|
| $10^{-5}$ | 15 | 15 (1) |
| $10^{-6}$ | 15 (2) | 15 (2) |
| $10^{-7}$ | 15 (3) | 15 (1) |
| $10^{-8}$ | 15 (4) | 15 |

As this was the first study of this kind in *Stx6*$^{-/-}$ mice, we proposed a shift in the attack rate by a 1 log dilution as a hypothesis on which to base power calculations. For example, wild type C57BL/6 mice infected with RML prions have an attack rate of 100% when inoculated with $10^{-6}$ RML concentration and a 20% attack rate with $10^{-7}$ concentration[2]. Based on our hypothesis, these numbers were used to determine the sample size using Fisher's exact test (two sided; 90% power). The suggested 8 animals/group, which was increased to 15 in line with precedent in similar types of studies and to allow for losses due to incurrent illness. This provided 90% power (2-sided) to detect a change in attack rate from 100% to 56%, or from 80% to 29% at a single dilution level. Statistical analyses for this experiment are described in **Section 1.3.5.**

*Two-Phase Kinetics RML Transmission Study in Stx6*$^{+/+}$ *and Stx6*$^{-/-}$ *Mice*

**Table 6. Timed Cull Schedule in the Two-Phase Kinetics Prion Transmission Study in *Stx6*$^{+/+}$ and *Stx6*$^{-/-}$ Mice.** Table detailing the predefined time points expressed as days post-inoculation (dpi) that animals were culled along with the numbers of animals in each arm of the study and any censored animals, which died due to unrelated health concerns.

| Time (dpi) | # RML-Infected | # Censored | # PBS-Inoculated |
|---|---|---|---|
| 30 | 5 $Stx6^{-/-}$ + 5 $Stx6^{+/+}$ | | - |
| 50 | 5 $Stx6^{-/-}$ + 5 $Stx6^{+/+}$ | 2* | 5 $Stx6^{-/-}$ + 5 $Stx6^{+/+}$ |
| 70 | 10 $Stx6^{-/-}$ + 10 $Stx6^{+/+}$ | | - |
| 90 | 10 $Stx6^{-/-}$ + 10 $Stx6^{+/+}$ | | - |
| 110 | 10 $Stx6^{-/-}$ + 10 $Stx6^{+/+}$ | | 5 $Stx6^{-/-}$ + 5 $Stx6^{+/+}$ |
| 130 | 10 $Stx6^{-/-}$ + 10 $Stx6^{+/+}$ | 1** | - |
| 140 | 10 $Stx6^{-/-}$ + 10 $Stx6^{+/+}$ | 2** | 5 $Stx6^{-/-}$ + 5 $Stx6^{+/+}$ |
| Endpoint | 30 $Stx6^{-/-}$ + 30 $Stx6^{+/+}$ | | - |

*These groups lost one animal/genotype during the procedure of inoculation so had to be removed from the study.

**Animals diagnosed with prion disease prior to their scheduled time cull so were excluded from follow-up analysis as they were not age-matched (1 $Stx6^{+/+}$ animal at 130 dpi and 1/genotype at 140 dpi).

Following inoculation with 1% (w/v) RML prions, female $Stx6^{+/+}$ and $Stx6^{-/-}$ mice were culled at multiple predefined time points or at onset of clinical disease**.** As controls, some mice were inoculated with Dulbecco's Phosphate Buffered Saline (DPBS, –$Mg^{2+}$ and $Ca^{2+}$). The number of animals in each arm and information about censored animals is detailed in **Table 6**. The disease outcome measures at each time point are summarised in **Table 7**. This was a female-only study to allow comparability with previous experiments. The main comparison in these studies were prion-infected $Stx6^{+/+}$ mice with $Stx6^{-/-}$ mice at each time point. A single animal was considered an experimental unit.

Group sizes for clinical endpoint were determined through power calculations using the effect size and variability from published work[1] (two-sided; 80% power). The suggested 22 animals/group was increased to 30 to allow for losses due to unrelated health concerns. Group sizes for the scheduled time culls were based on estimated effect sizes of the primary outcome measures, with 10 animals/arm giving 90% power to detect an 15% change in prion titre and with 5 animals/arm giving 90% power to detect a 21% change in prion titre.

**Table 7. Outcome Measures of the Two-Phase Kinetics RML Prion Transmission Study in $Stx6^{+/+}$ and $Stx6^{-/-}$ Mice.**

| Time (dpi) | Outcome Measure |
|---|---|
| 30 | SCA; IHC: Abnormal PrP |
| 50 | SCA; IHC: Abnormal PrP |
| 70 | SCA; IHC: H&E, Abnormal PrP |
| 90 | SCA; WB; IHC: H&E, Abnormal PrP, GFAP, Iba1 |
| 110 | SCA; Serum NfL |

| | |
|---|---|
| 130 | SCA; IHC: H&E, Abnormal PrP |
| 140 | SCA; WB; IHC: H&E, Abnormal PrP, GFAP, Iba1, synaptophysin Serum NfL, cell-based toxicity assay |
| Endpoint | Time to first symptom, incubation times |

SCA, scrapie cell assay; IHC, immunohistochemistry; H&E, hematoxylin and eosin; GFAP, glial fibrillary acidic protein; Iba1, Ionized calcium-binding adapter molecule 1; NfL, neurofilament-light chain.

Unless stated otherwise in the figure legend, statistical differences of the biochemical data were assessed by an unpaired t-test between infected *Stx6*$^{+/+}$ and *Stx6*$^{-/-}$ mice as the main experimental comparison. For neuropathological studies assessing multiple brain regions for each animal, a 2-way repeated measures mixed model approach was used for statistical analysis using the unstructured covariance structure to model the within-subject correlations, with genotype as the treatment factor and brain region as the repeated factor.

In some of the exploratory studies (neurotoxicity assay, biomarker assessment), one-way ANOVA followed by Fisher's LSD test of the following pre-planned comparisons:

1. Differences between infected *Stx6*$^{+/+}$ and *Stx6*$^{-/-}$ mice as the main experimental comparison.
2. Differences between RML-infected *Stx6*$^{+/+}$ mice relative to PBS-inoculated *Stx6*$^{+/+}$ controls to establish a disease-associated deficit.
3. Differences between PBS-inoculated *Stx6*$^{+/+}$ mice and PBS-inoculated *Stx6*$^{-/-}$ mice to assess for baseline differences.

To satisfy the parametric test assumptions, transformations were conducted in some cases, as detailed in the figure legends. Differences in the clinical endpoints were assessed as described in **Section 1.3.5.**

##### 1.3.3 Inoculation and Neurological Observation

10% (w/v) RML-infected C56BL/6 brain homogenate was diluted to 1% (w/v) in DPBS. Any subsequent dilutions for lower infecting doses were performed using 1% (w/v) normal C57BL/6 brain homogenate in DPBS as diluent. Brain homogenate dilutions were prepared on the same day and frozen at -80°C until infection.

6-9-week-old mice were anaesthetized with a mixture of isoflurane and $O_2$ and intracerebrally inoculated into the right parietal lobe with 30 µL of prion-infected brain homogenate or DPBS, as previously described[3,4]. Matched groups (set time cull or set prion dose) were infected in the same inoculation session to reduce confounding factors or split into two inoculation batches with a balanced genotype ratio if the former was unfeasible.

Following inoculation, all mice were examined daily for early indicators of clinical prion disease including piloerection, sustained erect ears, intermittent generalized tremor, unsustained hunched posture, rigid tail, mild loss of coordination, and clasping hind legs when lifted by the tail. Definite diagnosis of clinical prion disease was made when mice exhibited any 2 early indicator signs in addition to 1 confirmatory sign, or any 2 confirmatory signs, marking the experimental endpoint. The confirmatory signs included ataxia, impairment of righting reflex, dragging of hind limbs, sustained hunched posture, or significant abnormal breathing. Technical staff who made judgements on diagnosing prion disease were blinded to study design and endpoints were called by the same observing team.

##### 1.3.4 Humane and Elective Time Culls

All animals were culled upon definite prion disease diagnosis or at a defined, elective timed cull. Any animals that did not develop clinical prion disease were culled at 400 dpi with post-mortem assessment of prion-related neuropathological features. Animals were not followed up beyond this to avoid misdiagnosis due to the overlap of prion disease symptoms and general, non-specific characteristics of aged mice.

Animals were sacrificed by $CO_2$ asphyxiation. Brains were removed, dissected on the sagittal plane with the right hemisphere flash frozen and stored at -80°C, whilst the left hemisphere was fixed in 10% (v/v) formal buffered saline. Terminal blood was taken via post-mortem cardiac puncture and serum prepared by centrifuging for 10 min at 2600 rpm after allowing 10 min for clotting in Sarstedt Microvette Serum tubes.

##### 1.3.5 Endpoint Survival Statistical Analyses

*Attack Rate Analysis in the Serial Dilution of RML Transmission Study*

Attack rate was calculated as the cumulative number of incident cases of prion disease relative to the number of animals inoculated, with exclusions detailed in **Table 5**. Statistical differences between genotypes were determined by binary logistic regression analysis with genotype and dose as factors. This was complemented by calculating differences in the odds ratio of prion disease development across the different groups using the Baptista-Pike method to determine confidence intervals.

*Estimation of the "Effective Dose" Using the Spearman-Karber Method in the Serial Dilution of RML Transmission Study*

To estimate the "effective" dose infected into *Stx6*$^{+/+}$ and *Stx6*$^{-/-}$ mice in the RML serial dilution study, an adaption of the Spearman-Karber method[5] was used with definitions detailed in **Table 8** along with the following equation:

$\text{Log}_{10}$ 50% end point dilution = $- (x_o - d/2 + d \sum r_i/n_i)$

**Table 8. Adapted Definitions of Components of the Spearman-Karber Method.** Description of the metric components of the Spearman-Karber method that were adapted to be applicable to this study.

| Metric | Description |
|---|---|
| $X_0$ | $\text{Log}_{10}$ of the reciprocal of the highest dilution at there was an attack rate of >90%. |
| D | $\text{Log}_{10}$ of the dilution factor |

| | |
|---|---|
| $N_i$ | Average number of animals in each individual dilution after discounting unrelated deaths. |
| $\sum R_i$ | Sum of total number of positive animals (out of $n_i$) with summation started at $x_0$. |

*Comparison of Time to First Symptom and Time to Definite Clinical Prion Disease Diagnosis*

In the two-phase kinetics RML prion transmission study, disease-free incubation period was defined as the number of days from inoculation to the onset of the first prion disease-associated neurological symptom. Incubation period was defined as the number of days from inoculation until definite prion disease diagnosis[3]. Log-rank test survival analysis and the Kaplan-Meier method were employed to test for differences in the median time of these two parameters. Due to the anticonservative nature of the statistics, a 20% difference was defined as a biologically meaningful effect size. Animals culled due to unrelated health problems or that remained alive at the end of the study without a prion disease diagnosis were censored (detailed in **Table 6**).

Differences in disease duration were estimated as the difference between the time of first symptoms and definite prion disease diagnosis.

##### 1.3.6 Immunohistochemical Assessment

The fixed left hemisphere of the brain was embedded in paraffin wax followed by serial sectioning (4 μm nominal thickness) as previously described[6,7]. Sections were then deparaffinised prior to investigation of abnormal PrP (ICSM35 antibody), spongiosis (H&E), microgliosis (Iba1 antibody), astrocytosis (GFAP antibody) and synapse loss (anti-synaptophysin antibody) on the Ventana Discovery XT automated IHC staining machine with haematoxylin as the counterstain. Sections were treated using Ventana proprietary detection reagents before staining utilizing 3,3′-diaminobenzidine tetrahydrochloride as the chromogen (DAB Map Detection Kit) (**Table 9**). The Gemini AS Automated Slide Stainer was used for haematoxylin staining using a conventional approach.

**Table 9. Overview of Experimental Conditions for Prion-Related Neuropathological Assessment.**

| Primary Antibody | Dilution | Secondary Antibody | Dilution | Pretreatment |
|---|---|---|---|---|
| ICSM35 | 1:1000 | Biotinylated anti-mouse antibody | 1:200 | Treatment with discovery CC1 solution (95°C for 60 min) followed by Protease 3 (12 min) |
| Anti-GFAP | 1:1000 | Biotinylated anti-rabbit antibody | 1:200 | Protease 1 (4 min) |
| Anti-Iba1 | 1:1000 or 1:250 | Biotinylated anti-rabbit antibody | 1:200 | Treatment with discovery CC1 (95°C for 60 min) |

| | | | | |
|---|---|---|---|---|
| Anti-synaptophysin | 1:5000 | Biotinylated anti-rabbit antibody | 1:200 | Treatment with discovery CC1 solution (95°C for 60 min) |

Slides were digitally scanned on a LEICA SCN400F scanner and the NanoZoomer 360 at ×40 magnification, images captured from the NDP.serve3 or NZConnect software and composed with Adobe Photoshop.

*Immunohistochemistry Quantification and Statistical Analysis*

Animals were scored as positive or negative for abnormal PrP. For H&E, neuronal loss was scored in the hippocampus using 4-point score. Whole brain region immunostaining of GFAP and Iba1 was quantified using QuPath (v0.4.3) software with the experimenter blinded to the genotype of the sections. Colour deconvolution was performed followed by pixel classification to select tissue for analysis (whole brain). Artifacts were manually removed before pixel classification being applied to select regions positive for DAB staining allowing calculation of the % area stained relative to total tissue area analysed. A similar analysis was also done specifically in the thalamus, cortex and hippocampus, brain regions implicated in prion disease, with analysis being performed following manual annotation of the brain regions. Animals scored as non-infected by ICSM35 and H&E staining were excluded from the analysis.

##### 1.3.7 Biochemical Assessment

20% (w/v) homogenates were prepared in DPBS by ribolysing with 1.4 mm ceramic homogenisation beads at 6,500 rpm for 45 sec using the Ribolyser Precellys 24. Homogenates were stored at -80°C until use.

For immunoblotting, 1 µL benzonase was incubated with 20 µL aliquots of 10% (w/v) brain homogenates in DPBS. Following a 1 hr incubation with proteinase K (PK, 50 µg/mL) at 37°C and 800 rpm, samples were mixed with 2X sample buffer (125 mM Tris-HCl, 20% (v/v) glycerol, pH 6.8, containing 4% (w/v) SDS, 4% (v/v) 2-mercaptoethanol, 8 mM 4-(2-aminoethyl)benzenesulfonyl fluoride, and 0.02% (w/v) bromphenol blue). Samples were boiled at 100°C for 10 min, centrifuged at 16,100 *g* for 1 min, before being run on a 12% (w/v) NuPAGE gel at 200V for 55 min with MOPs running buffer and SeeBlue Prestained Molecular Weight Marker. They were then transferred to a PVDF membrane at 35V for 120 min.

Following blocking in PBS with 0.05% Tween-20 (PBST) with 5% (w/v) non-fat dried skimmed milk powder, membranes were probed with ICSM35 anti-PrP antibody (1:5000; RML) in PBST overnight. Following washing, the membranes were incubated with a 1:10,000 dilution of alkaline-phosphatase-conjugated goat anti-mouse IgG secondary antibody (A2179) in PBST. After washing (1 h with PBST followed by 2 × 5 min with 20 mM Tris pH 9.8 containing 1 mM $MgCl_2$), blots were incubated for 5 min in chemiluminescent substrate (CDP-Star) and visualized on Amersham Hyperfilm.

Alternatively, 20% (w/v) brain homogenates were digested with a working concentration of 100 µg/ml pronase for 40 min at 37°C at 1000 rpm. Digestion was terminated by the addition of EDTA at a final concentration of 100 mM with western blotting being performed as described above.

##### 1.3.8 Cell-Based Assessment of Infectivity of Infected Brain Homogenates

Infectivity in brain homogenates were assessed in either the automated scrapie cell assay (ASCA) or the SCA in endpoint format (SCEPA) with the exception of the excluded animals (**Table 6**) and the clinical endpoint group where animals were not age-matched. A more detailed account of the ELISpot revelation is described in **Section 1.4.6.**

*Automated SCEPA*

The infectivity of brain homogenates prepared from animals culled at 30 dpi, 50 dpi and 70 dpi were assessed by SCEPA as previously described[8], with the research team blinded to sample identity. In brief, an N2a-derived cell line (PK1/2 subclone) was plated in 96-well plates at 15,000 cells/well in OFBS medium (**Table 11**). The following day the cells were infected with $1 \times 10^{-4}$ or $1 \times 10^{-6}$ concentrations of 10% (w/v) brain homogenate (24 technical replicates/sample/dilution) diluted in OFBS using a Biomek FX liquid handling robot. Three days post infection, cells were split 1:3 and grown to confluence over a 2-day period. After two further 1:3 splits, cells underwent four 1:8 splits over 3-4-day periods with ELISpot plates being prepared at post-split 5 and 6. Here, ~25,000 cells/well were transferred to activated ELISpot plates, prior to PK treatment and development using the ICSM18 anti-PrP monoclonal antibody followed by an alkaline-phosphatase–conjugated anti-IgG1 polyclonal secondary antibody. The Bioreader 5000-Eβ system was used to quantify PK-resistant PrP-positive cells.

Infectivity of each sample was calculated from the ratio of positive to negative wells, aiming for a ratio between 30% and 70% for one of the two tested concentrations. Samples with a spot count above a defined cut-off, calculated from the average of the non-infected wells + (3 x standard deviation of non-infected wells) were scored as positive. TCIU/mL were calculated according to the Poisson equation, $P_{(0)} = e^{-m}$, where $P_{(0)}$ is the probability of a well remaining uninfected (i.e. non-infected wells/total number of wells). Therefore, TCIU/mL = m x dilution x 3.3, where m = ln [total wells/empty wells]. Samples with very low ratios at $1 \times 10^{-6}$ and with very high ratios at $1 \times 10^{-4}$ underwent a further round of SCEPA whereby cells were infected with $1 \times 10^{-5}$ concentration of the sample. Any samples with 100% infectivity at $1 \times 10^{-4}$ or $1 \times 10^{-6}$ were repeated in the SCA.

*Automated SCA*

The ASCA was used to assess infectivity in samples expected to be above $10^{4.5}$ TCIU $ml^{-1}$ in 10% (w/v) brain homogenate (90 dpi, 110 dpi, 130 dpi, 140 dpi and some 70 dpi repetitions). ASCA was performed similarly to SCEPA with the exception that $3 \times 10^{-5}$, $3 \times 10^{-6}$ and $3 \times 10^{-7}$ concentrations of brain homogenate were used for infection and four 1:8 splits were performed with intervals ranging from 3-4 days from the time of infection. Additionally, a reference serial dilution of 10% (w/v) RML brain homogenate of known infectivity titre (I8700, 8.64 logs TCIU $ml^{-1}$) was performed to generate a standard curve to deduce the prion titres of the experimental samples. Samples with no measurable activity were repeated.

*Data Analysis*

Samples with no measurable activity from 70 dpi onwards were excluded from analysis as this was not biologically feasible (3 *Stx6*$^{+/+}$ and 6 *Stx6*$^{-/-}$ mice). Zero values at 30 dpi and 50 dpi could not be excluded due to the very low titres detected in the positive cases. Therefore, they were assigned with a 3.1 log TCIU, the assay lower limit of detection based on historical data.

Curves were fitted using a logistic growth model, Y(1-Y/YM), where Y was the current titre and YM was the maximum titre.

##### 1.3.9 Cell-Based Assessment of Neurotoxicity of Infected Brain Homogenates

*Brain Homogenate Triple Extraction*

20% (w/v) brain homogenates were combined from 5 animals/genotype culled at 140 dpi to produce a 10% (w/v) brain homogenate pool in DPBS, which was supplemented with 1x Halt™ protease inhibitor cocktail (EDTA-free). Following a triple extraction method (unpublished), the brain homogenates were centrifuged for 10 min at 20°C at 1000 rpm. The supernatant was diluted in neurobasal media supplemented with 2% (vol/vol) B27, 0.25% (vol/vol) GlutaMAX, and 100 U/mL of penicillin–streptomycin to the final concentration of $5\times10^{-4}$ and $2.5\times10^{-4}$. These were applied to mouse primary cortico-hippocampal cultures prepared from embryonic day 17 FVB mouse brains plated at 8000 cells/well. Primary cells were used at the age of 8-10 days *in vitro*. There were 10 technical replicates per treatment with infected brain homogenates and 4 technical replicates per treatment with uninfected brain homogenates. These were tested in three independent experiments comprising independent cell cultures and independent brain homogenates extracts.

*IncuCyte Live Image Acquisition and NeuroTrack Analysis*

Cells were incubated at 37°C and 5% $CO_2$ within an IncuCyte S3 live cell imaging apparatus, with four images per well being acquired with a 20× objective lens in phase contrast from the same well coordinates. Images were subsequently analysed with the NeuroTrack module, IncuCyte S3 v.2018B with the following parameters typically being used: segmentation mode—brightness; segmentation adjustment—1.1; neurite sensitivity—0.35; filtering—best; neurite width—2 µm.

Changes in neurite length (average from 4 images in each well) upon treatment were normalised to the baseline neuronal length acquired for 18 hr before treatment, which was set at 100%. Statistical analysis was performed on normalised neurite length values 12 hr post treatment pooled from three independent experiments.

##### 1.3.10 Biomarker Assessment

Serum NfL levels were measured at 110 and 140 dpi in singulate using the single-molecule-array (Simoa) NF-light V2 Advantage Kit according to the manufacturer's instructions. This was run on the Simoa HD-X platform with the experimenter's blind to study design. Samples with no infectivity as determined by the ASCA/IHC were excluded from the final analysis.

##### 1.3.11 RNA-Sequencing

~50 mg cortical brain tissue from uninfected *Stx6*$^{+/+}$ and *Stx6*$^{-/-}$ mice (5 biological replicates/genotype) was homogenised in 900 µL DNA/RNA Shield in 2.0 mm Bashing Bead Lysis tubes using Precellys 24 (2 x 30 sec at 5500 rpm). Following a 5 min incubation on ice, samples were centrifuged at 16,000 *g* for 1 min with 300 µL of the cleared lysate subsequently being mixed with 300 µL TRI reagent in a RNAase-free tube before addition of 600 µL 98% ethanol. RNA was subsequently extracted using the Directzol RNA Miniprep kit according to the manufacturer's instructions including an on-column DNase step. An additional DNase digestion and RNA purification procedure was subsequently performed using the RNA Clean and Concentrator kit according to the manufacturer's instructions.

Quality control was performed on the Agilent Technologies 2200 Tape Station using the high sensitivity RNA reagents and on the NanoDrop ND-1000 Spectrometer as per the manufacturer's instructions. One *Stx6*$^{+/+}$ sample was discarded due to poor RNA integrity. 90 ng of total RNA (RIN$^{e}$: >7.9) was used with the Universal Plus Total RNA-Seq with Mouse AnyDeplete kit with library preparation being conducted according to the manufacturer’s instructions. Resulting libraries were visualized on the TapeStation to assess integrity using a D1000 ScreenTape and reagents, according to the manufacturer’s instructions.

Libraries were diluted to 10 nM and then pooled to 4 nM using Qiagen elution buffer. The pool was subsequently diluted to 800 pM with resuspension buffer Tween 20, which was sequenced using the Illumina NextSeq 2000 Kit P2 100 cycles (sequencing configuration: 61-8-8-61) at a loading concentration of 800pM after the addition of 2% PhiX sequencing control library. The data was demultiplexed and converted into a FASTQ file using BCLconvert v3.7.5. Data was analysed using the R software[9] and Bioconductor[10] packages including DESeq2[11,12] and the SARTools R package[13].

#### 1.4 Prion-Based Cellular Work

##### 1.4.1 Statistics

Unless stated otherwise in the figure legend, statistical differences were assessed by one-way ANOVA followed by Fisher’s LSD test of pre-planned comparisons. To satisfy the assumptions of ANOVA, transformations were conducted in some cases, as detailed in the figure legends.

##### 1.4.2 General Tissue Culture Practice and Cell Culturing

Cells were routinely split 1:6 every 3 days or 1:8 every 4 days and kept in an incubator at 37°C/5% $CO_2$ and cultured in the media detailed in **Table 11**.

**Table 10. Media Composition for Cell Lines Employed.**

| Cell Line | Culture Media |
|---|---|
| CAD5 | Opti-MEM + 10% Bovine Growth Serum (BGS) + 1% penicillin-streptomycin (PS) (OBGS) |
| PK1 | Opti-MEM + 10% Fetal Bovine Serum (FBS) + 1% PS (OFBS). |
| HEK | DMEM + GlutaMAX-I + 10% FBS + 1% PS (C-DMEM) |

##### 1.4.3 Generation of Cell Lines with Manipulated *Stx6* Expression

Cell lines were generated by different approaches, with specific experimental conditions detailed in **Table 12** and plasmids sequences specified in **Table 13**.

*PiggyBac Transgenesis*

500,000 cells/well were seeded into a 6-well plate in 2 mL media. The following day, a premix containing the PiggyBac transposon vector (mCherry-CAG-hyPBase) and PiggyBac transposase vector (**Table 13**), 175 µL Opti-MEM and 3.5 µL PLUS reagent was added to 150 µl Opti-MEM with 6 µl Lipofectamine LTX Reagent. After a 5-10 min incubation period at room temperature (RT), 250 µL of this transfection mix was added to the cells. The following day the cells were placed under puromycin selection (2 µg/mL for CAD5 cells; 3 µg/mL for PK1 cells; 4 µg/mL for iS7 cells) with media exchanges every 2-3 days.

**Table 11**. **Approaches of Cell Line Generation with Specific Experimental Conditions.** Cell lines were either generated by PiggyBac transgenesis or stable transfection as specified by this table. For PiggyBac transgenesis, cells were co-transfected with a PiggyBac transposon vector (pPB-transposon) and a vector encoding PiggyBac transposase (PBase). For overexpression cell lines generated by PiggyBac transgenesis, a titration of pPB-transposon was performed to limit the copy number integrations. For stable transfections, the pPB-transposon vector was either used in the absence of the PBase vector or alternatively pGFP-V-RS vectors were used.

| **Cell Line** | **Method of Generation** | **Specific Conditions** |
|---|---|---|
| PK1 *Stx6* Knockdown | PiggyBac transgenesis and pool analysis. | 1 µg pPB-transposon + 1 µg Pbase vector |
| PK1 *Stx6* Overexpression | Stable transfection and pool analysis. | OE1: 1250 ng pPB-transposon<br>OE2: 2500 ng pPB-transposon |
| CAD5 *Stx6* Knockdown | Stable transfection and single cell cloning. | ~2500 ng pGFP-V-RS vector (D clones) or 875 ng pGFP-V-RS vectors encoding shSTX6_A, shSTX6_B, shSTX6_C and shSTX6_D for ALL condition |
| CAD5 *Stx6* Overexpression | PiggyBac transgenesis and pool analysis. | 1 µg Pbase vector +<br>OE1: 1 µg pPB-transposon<br>OE2: 0.1 µg pPB-transposon<br>OE3: 0.05 µg pPB-transposon |
| IF11 *Stx6* Knockdown | PiggyBac transgenesis and pool analysis. | 1 µg pPB-transposon + 1 µg Pbase vector |

OE, overexpression.

*Stable Transfection*

Transfection was performed as described above with the exclusion of the PiggyBac transposase vector. One day post-transfection, the cells were expanded to a 10 cm dish before being placed under antibiotic selection the following day. Media exchanges were performed every 2-3 days for approximately 2-4 weeks until the appearance of distinct colonies of cells. For cell pools, these colonies were pooled into a single suspension and propagated as a cell population. For single cell clones, individual clones were picked and expanded independently to provide genetically homogenous and clonal expansion.

**Table 12**. **Plasmid Constructs Used for Stable Cell Line Generation.** All plasmids had a puromycin selection cassette.

| **Construct** | |
|---|---|
| **shRNA Plasmids** | **Targeted Sequence (5'-3')** |
| pPB-EGFP shSTX6_4 | TGGAATGCTGGAGTGGCAGATCGCTATGG |
| pPB-EGFP shNSC | GCACTACCAGAGCTAACTCAGATAGTACT |

| | |
|---|---|
| pPB-EGFP shPRNP | GAGACAATCTAAACATTCT |
| pGFP-V-RS shSTX6_A | CAGATGTCAGCTTCATCTGTGCAGGCACT |
| pGFP-V-RS shSTX6_B | GGAGAGGTACAGAAAGCAGTCAACACTGC |
| pGFP-V-RS shSTX6_C | CCAGTGGTGTGCCATAGCCATCCTCTTCG |
| pGFP-V-RS shSTX6_D | TGGAATGCTGGAGTGGCAGATCGCTATGG |
| pGFP-V-RS shNSC | GCACTACCAGAGCTAACTCAGATAGTACT |
| pRetroSuper PRNP (Si8) | GAGACAATCTAAACATTCT |
| **Overexpression Plasmids** | **Encoded Sequence** |
| pPB-EGFP mStx6 | Mouse *Stx6* transcript (NM_021433.3) |
| pPB-EGFP ORF-Stuffer | Amino acid 2-83 of E. coli beta-galactosidase |

##### 1.4.4 Western Blotting

*Cell Lysate Preparation*

For syntaxin-6 and $PrP^C$ western blotting, cell suspensions were harvested by centrifugation at 500 *g* for 4 min with the cell pellet being subsequently washed in ice-cold DPBS. Pellets were lysed in RIPA buffer supplemented with 1x HALT protease inhibitor. Following a 10 min incubation on ice with frequent vortexing, the cell lysates were centrifuged at 16 000 *g* for 10 min at 4˚C. The supernatant was collected and stored at -20˚C until use.

For abnormal PrP, cells were collected from each well of a 6-well plate in 1 mL ice-cold DPBS on ice and spun at 300 *g* for 4 min at 4°C. The pellet was lysed in RIPA buffer supplemented with 4 μL/ml benzonase on ice with gentle agitation employed every 5 min for 20 min.

*Protein Quantification by the Bicinchoninic Acid (BCA) Assay*

For total protein concentration estimation, Pierce™ BCA Protein Assay kit was used according to the manufacturer's instructions. Serial dilutions of the calibrator (BSA) were prepared in the relevant buffer (0, 0.125, 0.25, 0.5, 0.75, 1, 1.5, 2 mg/mL). Lysates were diluted in RIPA buffer and 20 μL of each sample loaded in duplicate in a clear 96-well microplate. The working solution was then added at a final volume of 200 μL per well. The plate was then sealed and incubated for 30 min at 37°C.

Absorbance was measured at 560 nm on the Tecan Infinite F200 Pro plate reader using iControl software. CV% values (STD/Average*100) were calculated to ensure low variability between duplicate values. To determine protein concentration, an average of the absorbance values of the duplicates was calculated. The calibrator was used to generate an 8-point standard curve to which a 4-parameter logistic curve was fitted. The unknown concentrations of total protein were extrapolated from the standard curve with the final concentration being corrected by multiplying by the dilution factor.

*Immunoblotting*

Lysates were diluted in DPBS and 4x Laemelli sample buffer to obtain a final 1x concentration with 355 mM final 2-mercaptoethanol. Following boiling at 95°C for 5 min, samples were loaded onto a 4-12% Bis-Tris polyacrylamide gel in addition to the SeeBlue Protein ladder. Following electrophoresis at 180V for 1 hr, protein was electroblotted onto a nitrocellulose membrane at 35 V for 2 hr. Following a 1 hr incubation at RT in Odyssey PBS Blocking Buffer with agitation, membranes were probed either with anti-syntaxin-6 (clone C34B2; 1:500) or with monoclonal anti-PrP antibody (clone ICSM18; 1:1,250) overnight at 4°C. 1:1 dilution of Odyssey Blocking Buffer and PBST was used as diluent for all antibody incubation steps.

For syntaxin-6 immunodetection, membranes were washed 3x with PBST for 5 min with agitation, followed by a 1 hr incubation with anti-β actin (ab6276; 1:5000) at RT with agitation. After 3x 5 min washes in PBST, membranes were probed with fluorophore-conjugated secondary antibodies IRDye 800CW Donkey anti-rabbit IgG (1:4000) and IRDye 680RD goat anti-mouse IgG (1:20,000) for 1 hr at RT with agitation and protected from light. Following 3x 5 min washes in PBST, membranes were imaged in PBS using the Odyssey® Imaging System (LI-COR; Model 9120).

Revert™ 700 Total Protein Stain was used for normalisation for some syntaxin-6 western blots, which was used according to the manufacturer's instructions. Membranes were imaged in the 700 nm channel before being blocked and treated as described above excluding the β-actin immunostaining.

For PrP immunodetection, membranes were washed with PBST for 5 min x3 with agitation, followed by a 1 hr incubation with IRDye 680RD goat anti-mouse IgG (1:10,000, LI-COR) at RT with agitation protected from light. After 3x 5 min washes in PBST, membranes were imaged in PBS using the Odyssey Reader System. Membranes were subsequently incubated with rabbit anti-β actin (A2066; 1:1000). Following 3x 5 min washes in PBST, membranes were probed with IRDye 800CW Donkey anti-rabbit IgG (1:10,000) for 1 hr at RT with agitation. Following 3x 5 min washes in PBST, membranes were imaged in PBS.

Abnormal PrP western blotting was performed as described in **Section 1.3.7** with gel electrophoresis being performed using 12% NuPAGE gels run at 200 V for 50 min followed by 2 hr transfer at 35 V with 6D11 being used as the primary antibody (1:5000, ON).

*Western Blot Analysis*

Image Studio™ Software (licor.com/islite) was used to quantify the fluorescent signals from the 700 nm and 800 nm channels. Rectangles were drawn around the target bands as well as the loading control bands. Median background subtraction method was used. Following quantification, the *Lane Normalisation Factor* (LNF) was calculated as follows:

$$LNF = \frac{\beta - Actin\ Signal\ for\ Each\ Lane}{\beta - Actin\ Signal\ from\ Lane\ with\ Highest\ \beta - Actin\ Signal}$$

The normalised signal was then calculated using the following formula:

$$Normalised\ Signal = \frac{Target\ Signal\ for\ Each\ Lane}{LNF\ for\ Each\ Lane}$$

##### 1.4.5 Growth Rate Analysis

3.5 x $10^5$ cells were seeded in a T25 flask with cell doubling time being calculated after 4 days using the following equation, where $N_4$= cell number on day 4 and $N_0$ = 3.5 x $10^5$.

$$Doubling\ time\ (days) = \frac{4}{\log_2 \frac{N_4}{N_0}}$$

##### 1.4.6 Scrapie Cell Assay (SCA)

*Automated Scrapie Cell Assay*

The ASCA was performed as previously described[8] with some modifications. Cells were counted using a C-CHIP disposable haemocytometer and subsequently seeded at 18,000 cells/well in a 96-well plate. The following day, the cells were infected with a dilution series of 10% (w/v) prion-infected brain homogenate. Uninfected cells were also included to provide an estimation of the background spot count. Cells were passaged 1:6 every 3 days or 1:8 every 4 days, using the automated Biomek FX liquid handling robot.

After a set number of passages, 85 µL of the 270 µL cell suspension was plated on ELISpot IP Filter Plates (PVDF membrane, 0.45 µm), which had been activated with 30% (v/v) ethanol. Following two PBS washes, the membranes were dried at 50°C. Plates were subsequently treated with proteinase K (1:10,000) in lysis buffer (50 mM Tris HCl pH 8, 150 mM NaCl, 0.5% (w/v) sodium deoxycholate, 0.5% (v/v) Triton X-100) at 37°C for 1 hr. Plates were washed twice in PBS before being treated with 3M guanidine thiocyanate in 10 mM Tris.HCl (pH 8) and incubated at RT for 20 min for decontamination and antigen retrieval. Following seven PBS washes, plates were incubated with 0.5x SuperBlock TBS blocking buffer for 30 min at RT.

For immunodetection, plates were incubated with an anti-PrP antibody (clone ICSM18; D-Gen Ltd; 0.4 µg/mL) in Tris Buffered Saline with Tween 20 (TBST)/1% milk power at RT for 1 hr. After 5 washes with TBST, plates were incubated with an alkaline phosphatase-linked anti-mouse secondary antibody (IgG1-AP; 1:9000) in TBST/1% milk power at RT for 1 hr. Following 5 further washes in TBST, the plates were dried and treated with the alkaline phosphatase conjugate substrate and incubated for 30-60 min at RT to allow visualisation of the spots. Following two washes with dd$H_2O$, plates were dried and the number of PK-resistant PrP infected cells were counted using the Bioreader 5000-Eβ.

For the secreted infectivity of iS7 cells with syntaxin-6 manipulation, 100 µL of 1:10 dilution of cells in DPBS was plated on ELISpot. For assessment of PK1 *Stx6* OE *de novo* infection secreted infectivity, 85 µL of the 270 µL cell suspension was plated on ELISpot.

*Manual SCA*

For infection, a cell suspension (50,000 cells/mL) was prepared with 3 µl exosome-enriched cell supernatants/mL (prepared from iS7 cells as described in[14]) with 300 µl plated/well in a 96-well plate for 3-4 days. Cells were subsequently split 1:5 or 1:6 every 3-4 days. For chronically infected cells, 300 µl cells (50,000 cells/mL) were plated in 96-well plates with ELISpot assessment of prion steady state levels being conducted across three passage numbers.

The SCA was performed as above with the following modifications: ELISpot plates were activated in 50% ethanol, differing volumes of cells were plated on ELISpot (**Table 14**), blocking was performed for 1 hr, the primary antibody employed was 6D11 (1:10,000) and 5x TBS-T washes were conducted following the secondary antibody incubation.

**Table 13. Volumes of Cells Plated on ELISpot Plates.**

| Cell Line | Volume Plated on ELISpot (µL) |
|---|---|
| PK1 *Stx6* Knockdown | 50 µL of neat cell suspension |
| PK1 *Stx6* Overexpression | 100 µL of 1:10 dilution of cells in DPBS |
| CAD5 *Stx6* Overexpression | 40 µL of neat cell suspension |
| iS7 *Stx6* Knockdown | 50 µL of 1:10 dilution of cells in DPBS |

*Haematoxylin Staining*

Haematoxylin was filtered with a 0.22 µm filter before being diluted 1:10 in $H_2O$. 100 µL/well was added for 1 min to the ELISpot plate (post-immunostaining) followed by two washes with $H_2O$. Plates were subsequently read using the Bioreader 5000-Eβ reader with high sensitivity parameters. The haematoxylin total cell count was used to normalise the raw spot count providing an estimation of the proportion of infected cells in assays where the cell count was not saturated.

##### 1.4.7 Confocal Microscopy

500 µL cell suspension (50,000 cells/mL) were plated in 8-well chambered glass coverslides and incubated at 37°C/5% $CO_2$ for 4 days before fixation for 12 min with 3.7% formaldehyde in PBS. As previously described for $PrP^d$[14,15], the cells were washed once in PBS before a 1 min incubation with chilled acetone. Following one PBS wash, cells were incubated for 10 min with 3.5 M guanidine thiocyanate before five further washes in PBS.

Cells were incubated at 4°C overnight with primary antibodies (**Table 15**) diluted into Superblock (1:4 dilution in PBS/10% PS). After one wash with PBS, cells were labelled with fluorescence-conjugated secondary antibodies (1:1000) and DAPI (1:10,000) and incubated at 4 °C overnight. Following a final wash with PBS, cells were stored in PBS/10% PS at 4°C until imaging.

**Table 14**. **Primary Antibody Dilutions.** Full antibody details can be found in the key resources table.

| Antibody | Dilution |
|---|---|
| Anti-PrP (6D11) | 1:10,000 |
| Anti-PrP (5B2) | 1:500 |
| Anti-Lamp1 (1D4B) | 1:1000 |
| Anti-Eea1 (C45B10) | 1:1000 |
| Anti-TGN46 (ab16059) | 1:1000 |

*Image Acquisition*

A Zeiss LSM710 laser-scanning microscope was used for image acquisition using a ×63 objective (1.4 oil) with Immersol immersion oil. The 460-540 nm and 565–640 nm bandpass filters were employed to measure AlexaFluor488 and Rhodamine Red-X fluorescence, subsequent to excitation with an argon laser at 488 nm and a diode-pumped solid state laser at 561 nm, respectively.

For PK1 *Stx6* knockdown cells infected with exosomes, an automated scan of 7 x 7 tiles, with approximately 25 cells per tile, was set up for each condition (images with <11 cells were excluded). Using Volocity, the signal of 5B2 was digitally increased by 3 iterations of dilation, before quantifying its intersection with 6D11, which was used as a proxy of plasma membrane staining based on[14]. To quantitatively and specifically assess perinuclear $PrP^d$, the 6D11 fluorescence intensity and area was determined excluding 6D11 signal touching the intersection with 5B2. For each image, the perinuclear $PrP^d$ signal was normalised to the number of cells in the field view as determined by Volocity quantification of the DAPI count. Tile scans were also conducted in chronically infected cells for quantification of the total signal intensity and area of 5B2 and 6D11 staining using Volocity.

To determine levels of colocalisation of 6D11-positive $PrP^d$ with intracellular markers in chronically infected cells, ~20 magnified images were acquired, which were subsequently used to determine Pearson's correlation with Costes threshold correction[16] using Velocity. Negative values were computed as zero. Additionally, the Volocity cropping function was used to manually isolate the perinuclear 6D11-positive $PrP^d$ signal which was subsequently quantified in terms of fluorescence intensity and area.

To quantitatively assess morphological differences in fibril-like $PrP^d$ aggregates at the plasma membrane, stacks of 10 images in the z-direction above the basement membrane were recorded with a step size of 0.1 µm with stacks subsequently being aggregated into a maximum intensity projection. Length of fibrils were measured using the line tool on Volocity in a blinded fashion.

##### 1.4.8 Cell Viability Assay

Cell viability was assessed using the CellTitre-Glo luminescent assay. In brief, cells were split into a CELLSTAR white 96-well plate with a micro-clear bottom. After 3-4 days of growth, media was removed from cells and the reconstituted CellTitre-Glo buffer/substrate mix was added (75 µL/well). On the TECAN Infinite F200 PRO machine, the plate underwent shaking for 2 min followed by a 10 min incubation before luminescence was recorded.

##### 1.4.9 Flow Cytometry Quantification of Total and Cell Surface PrP Levels

Cells were resuspended and counted on the Countess 3 FL Automated Cell Counter using the live cell count. 0.5 $\times 10^6$ cells were centrifuged at 1500 rpm for 5 min at 4°C, with these centrifugation settings being used for all further centrifugations. Cells were then resuspended in 250 µL of BD Cytofix/Cytoperm Fixation/Permeabilisation solution for 20 min on ice, followed by two washes 1× BD Perm/Wash™ buffer. For assessment of cell surface PrP levels, this permeabilisation step was excluded and instead the cell pellet was washed once in PBS/1%FBS.

Cells were subsequently incubated with an anti-PrP antibody (ICSM18 subclone, 10 µg/mL) diluted in PBS/1%FBS for 30 min on ice. Following another wash in PBS/1%FBS, the cells were incubated with an AlexaFluor 647-conjugated secondary antibody diluted in PBS/1%FBS for 30 min on ice (1:1000). Following another wash step, the cells were resuspended in 100 µL PBS/1%FBS, which was analysed using a Beckman Coulter CytoFLEX equipped with blue 488nm and red 633nm lasers, with AF647 being detected using 660/10 filter.

Cells were gated based on the forward and side scatter profiles using forward scatter width to identify single cells. A total of 50,000 events in the single-cell gate were recorded per sample. A secondary only control was included in all experiments to aid gating. Further analyses were carried out in FlowJo software with median fluorescence of anti-IgG-AF647 binding to ICSM18 being used to determine differences in total or cell surface PrP levels.

##### 1.4.10 RT-qPCR Assessment of *Prnp* RNA Levels

*Prnp* RNA levels were assessed by performing two-step RT-qPCR according to the manufacturer's instructions using the Quantitect Reverse Transcription Kit with 2 µl RNA input. cDNA was amplified using QuantiText Primer Assays against *Prnp* (Mm_Prnp_1_SG) and ActB (Mm_Actb_1_SG) as the loading control.

##### 1.4.11 Cycloheximide Assay Assessing PrP Degradation Kinetics

PK1 overexpression cell lines were seeded in 6-well format (0.3 x $10^6$ cells/well). After 48 hr, media was exchanged with fresh media containing 100 µg/mL cycloheximide or DMSO vehicle control. Cells were subsequently harvested at 0 hr, 3 hr and 6 hr (3 replicas/cell line/time point) and immediately fixed in Fixation/Permeabilization solution. The following day, total PrP levels were assessed by flow cytometry as described in **Section 1.4.9.** The initial fluorescence at 0 hr was used for normalisation of each cell line to monitor relative protein decay over time.

##### 1.4.12 Assessment of Secreted Infectivity

*iS7 Cells with Transient Stx6 Knockdown*

iS7 cells (1.33 x $10^5$ cells/mL) were reverse transfected with pools of 30 custom-designed siRNAs (siPools)[17] against *Stx6* or a non-silencing control. In brief the siPools were prepared at a stock concentration of 10 µM in nuclease-free water. 2 µL siPool and 6 µL RNAimax transfection reagent were incubated with 42 µL Opti-MEM for a 5 min incubation. 950 µL OFBS was subsequently added to the mix. 150 µL was then added to each well of a 96-well plate (12 wells/condition) followed by 150 µL cell suspension and left to grow for 3 days. On day 3, the media was replaced for a 7-hr incubation. Media from the 12 wells/condition was pooled, clarified at 10,000 *g* for 10 min and stored in the fridge until the time of infection. Cells were collected and washed in DPBS to confirm successful knockdown by immunoblotting as described in **Section 1.4.4**.

*PK1 Cell Lines with Stable Syntaxin-6 Manipulation*

For secreted infectivity assessment following acute infection of PK1 cell lines with stable *Stx6* manipulation, after 3-4 days of growth, the conditioned media from 3 wells of a 6 well plate/cell line was harvested. This was clarified at 4,700 *g* for 10 min and stored at -20°C until the time of infection.

*iS7 Cell Lines with Stable Syntaxin-6 Manipulation*

iS7 cells (180,000 cells/mL) were seeded in a 12-well plate (4 wells/cell line) and left to grow for 4 days. On day 4, the media was replaced with 1 mL fresh media/well. Following a 6 hr incubation, the conditioned media was collected and pooled per cell line and subsequently clarified at 10,000 *g* for 10 min and stored this at -20°C until the time of infection.

*Infection*

The clarified supernatant was used to infect reporter PK1 cells that had been seeded the day before with the SCA being conducted as described in **Section 1.4.6** with cells being diluted 1:10 prior to plating on ELISpot across four splits.

#### 1.5 Time Course Investigation of *Stx6*$^{+/+}$;h*Tau*$^{P301S/P301S}$ and *Stx6*$^{-/-}$;h*Tau*$^{P301S/P301S}$ Mice

##### 1.5.1 Breeding Strategy

*Stx6*$^{-/-}$ mice on the C57BL/6N background[1] were crossed with Thy1-hTau.P301S (CBA.C57BL/6J) mice[18] (referred to as h*Tau*$^{P301S/P301S}$ mice hereafter) to generate a fully heterozygote cohort, which were subsequently intercrossed. These offspring were used to establish littermate-derived *Stx6*$^{+/+}$;h*Tau*$^{P301S/P301S}$, *Stx6*$^{-/-}$;h*Tau*$^{P301S/P301S}$, *Stx6*$^{+/+}$;m*Tau*$^{+/+}$ *and Stx6*$^{-/-}$;*mTau*$^{+/+}$ animals to populate the experiment with lines subsequently being maintained as homozygous x homozygous breeding pairs. General animal husbandry and genotyping was conducted as detailed in **Section 1.2,** however due to the severity of the phenotype of the h*Tau*$^{P301S/P301S}$ mice, the breeding life span of females was restricted to 3 litters only or if either breeder was observed to have first signs of the disease phenotype.

Experimental animals were born within a 10-day time window to ensure behavioural analysis was performed on animals of a similar age. For all genotypes except wildtype animals (15 females and 5 males), there was an equal sex balance of experimental animals.

##### 1.5.2 Study Design, Power Calculations & Statistical Analysis

For animals undergoing long-term phenotypic assessment, power calculations were conducted for the primary clinical endpoint, which determined that an n=20 per arm would provide 90% power to detect a 7% change in time to cull, which would provide sufficient animals for robust behavioural characterisation whilst accounting for incurrent loss of animals. This was a mixed sex study in line with 3Rs increasing the generalisability of the findings. The number of animals in each arm of the study as well as censored animals are detailed in **Table 16**. A single animal was considered an experimental unit.

**Table 15. Study Design of Time Course Investigation of *Stx6*$^{+/+}$;h*Tau*$^{P301S/P301S}$ and *Stx6*$^{-/-}$;h*Tau*$^{P301S/P301S}$ Mice.** Total number in each arm of the clinical endpoint groups are detailed below. In brackets are the number of animals which underwent behavioural analysis (rotarod, gait assessment). Censored animals are documented in the additional notes.

| Mouse Line | # Female | # Male |
|---|---|---|
| *Stx6*$^{+/+}$;h*Tau*$^{P301S/P301S}$ | 10 (8) | 10 (7) |

| | | |
|---|---|---|
| *Stx6*$^{-/-}$;h*Tau*$^{P301S/P301S}$ | 10[a] (8) | 10 (7) |
| *Stx6*$^{+/+}$;m*Tau*$^{+/+}$ | 15 (10) | 5 (5)[b] |
| *Stx6*$^{-/-}$;m*Tau*$^{+/+}$ | 10 (8) | 10 (7) |

[a]One animal culled due to an eye defect at 6.8 months so was censored in survival analysis but data points were included for the behavioural analysis.

[b]One animal lost in month 5 and a further one lost in month 6, resulting in missing rotarod data.

Age-matched time culls were also performed at 1 month, 2 months, 3 months and 5 months (n=10/genotype).

*Statistical Analyses*

The main experimental comparison was pre-defined to be between *Stx6*$^{+/+}$;h*Tau*$^{P301S/P301S}$ and *Stx6*$^{-/-}$;h*Tau*$^{P301S/P301S}$ mice. As secondary comparisons, *Stx6*$^{+/+}$;m*Tau*$^{+/+}$ and *Stx6*$^{-/-}$;m*Tau*$^{+/+}$ mice were compared to inform on baseline deficits, as well as a comparison between *Stx6*$^{+/+}$;m*Tau*$^{+/+}$ and *Stx6*$^{+/+}$;h*Tau*$^{P301S/P301S}$ mice to inform on the disease-relevance of the readout.

Unless stated otherwise in the figure legend, statistical differences were assessed by one-way ANOVA followed by Fisher's LSD test of the pre-planned comparisons stated above. To satisfy the assumptions of ANOVA, transformations were conducted in some cases, as detailed in the figure legends. For neuropathological studies assessing multiple brain regions for each animal, a 2-way repeated measures mixed model approach was used for statistical analysis using the unstructured covariance structure to model the within-subject correlations, with genotype as the treatment factor and brain region as the repeated factor. Graphing and statistical analysis were conducted in Graphpad Prism and InVivoStat.

##### 1.5.3 Rotarod Analysis to Assess Motor Function

Rotarod performance was assessed monthly (mixed sex) as previously described[19] with the weight of each animal being recorded prior to each session. For the first month, training sessions were conducted over 3 consecutive days to allow mice to reach a steady baseline level of performance, which was subsequently reduced to 1 day of training prior to the test day in the subsequent months.

The rotarod apparatus was placed in a quiet space with mice being acclimatised to the new environment prior to testing. Each training session consisted of four trials at a constant speed (20 rpm) for a maximum of 60 sec, with the latency to fall being manually recorded within this time period. In the testing session, mice underwent two trials at 5 increasing speed levels (8, 16, 24, 32, 40 rpm) for a maximum of 60 sec. The 24 rpm speed was tested first. Providing all animals passed, they were tested at the higher speeds and assumed to pass the lower speeds. If there was a <100% pass rate, the lower speeds were subsequently assessed. The mean latency to fall (for the two trials at each speed level) was calculated and used in subsequent analysis.

At 6 months, animals with clasping hindlimbs were not assessed on the rotarod for welfare reasons and were imputed as zero for statistical analysis. Two *Stx6*$^{+/+}$;m*Tau*$^{+/+}$ mice were culled due to unrelated health problems so were not tested on months 5 and 6. The data points proceeding the deaths of these animals were still included. All genotypes and sexes were tested in the same session, at the same time of the day each month, with the experimenter blind.

##### 1.5.4 Ink Blot Analysis for Gait Assessment

Footprint assessment was performed in animals at 5.5 months of age (mixed sex) to detect gait abnormalities. Here, the paws of mice were inked with a nontoxic dye before they were released on one end of an absorbent paper track, which had a width of 10 cm. The base of each pawprint was marked to provide a point for the subsequent measurements taken.

Each gait parameter (n=12) was analysed using one-way ANOVA, with genotype as the treatment factor and sex as a blocking factor. This was followed by the planned comparison between *Stx6*$^{+/+}$;h*Tau*$^{P301S/P301S}$ and *Stx6*$^{+/+}$;m*Tau*$^{+/+}$ mice to prioritise the most robust disease-relevant gait parameters with a deficit (set P-value threshold = 0.0042; unidirectional selection of measurements with a deficit **Table 17**). These were used to assess for phenotypic rescue in *Stx6*$^{-/-}$;h*Tau*$^{P301S/P301S}$ mice.

**Table 16. Gait Measurements and Basis for Selection of Disease-Relevant Gait Parameters.** The 12 measurements taken are detailed with the preceding “L” or “R” referring to left and right side of the body respectively; and the following “F” or “H” referring to the fore and hind limbs respectively. Disease-relevant parameters with a deficit were identified by comparing *Stx6*$^{+/+}$;h*Tau*$^{P301S/P301S}$ and *Stx6*$^{+/+}$;m*Tau*$^{+/+}$ mice (P-value threshold = 0.0042) and are highlighted in grey.

| | ***Stx6*$^{+/+}$;h*Tau*$^{P301S/P301S}$ *vs.* *Stx6*$^{+/+}$;m*Tau*$^{+/+}$ Mice** | |
|---|---|---|
| **Measurement** | **Difference (Mean [SEM])** | **P-Value** |
| Stride Length (RH-RH) | -2.12 [0.35] | < 0.0001 |
| Stride Length (LH-LH) | -2.01 [0.39] | < 0.0001 |
| Ipsilateral Together (RH_RF) | 0.60 [0.18] | 0.0021 |
| Ipsilateral Together (LH_LF) | 0.79 [0.20] | 0.0003 |
| Ipsilateral Apart (RF_RH) | -1.45 [0.33] | 0.0001 |
| Ipsilateral Apart (LF_LH) | -0.93 [0.43] | 0.037 |
| Contralateral Difference (Fore) | -1.02 [0.32] | 0.0021 |
| Contralateral Difference (Hind) | -0.84 [0.33] | 0.014 |
| Stride Width (LF_RF) | 0.09 [0.11] | 0.40 |
| Stride Width (LH_RH) | -0.25 [0.10] | 0.016 |
| Ipsilateral Width (RF_RH) | -0.19 [0.09] | 0.035 |
| Ipsilateral Width (LF_LH) | -0.08 [0.14] | 0.56 |

To calculate the composite score, the population mean ($\bar{x}$) and standard deviation (SD) were calculated for each prioritised disease-relevant measurement across all groups. Subsequently, a Z score for each parameter was calculated for each animal, where $z = (x - \bar{x})/SD$, where x was the mean of each individual animal. An average Z score for each mouse was then calculated, with data finally being analysed using one-way ANOVA with genotype as the treatment factor and sex as a blocking factor followed by planned comparisons.

##### 1.5.5 Frailty Assessment

In a separate cohort of mice, *Stx6*$^{+/+}$;h*Tau*$^{P301S/P301S}$ (n=26), *Stx6*$^{-/-}$;h*Tau*$^{P301S/P301S}$ (n=18), *Stx6*$^{+/+}$;m*Tau*$^{+/+}$ (n=5) and *Stx6*$^{-/-}$;m*Tau*$^{+/+}$ (n=5) underwent frailty assessment in three batches (mixed sex)**.** Specifically, mice were assessed for the presence, absence and severity of the following 18 frailty characteristics: alopecia, loss of fur colour, dermatitis, righting reflex, coat condition, piloerection, eye discharge/swelling, microphthalmia, cataracts, nasal discharge, body condition, kyphosis, impaired gait during free walking, tremor, breathing rate/depth, tail stiffening, vestibular disturbance and distended abdomen. This represented a condensed set of parameters derived from a validated, established scoring system described in Rizzo *et al.* (2018)[20]. A 3-point score was assigned to each parameter: no deficit (0), mild deficit (0.5) or severe deficit (1). The assessment was conducted by the same experienced technician who was blind to genotype. Mice were evaluated for each measure independently followed by calculation of a cumulative score of all measures for each animal (max score = 18).

##### 1.5.6 Clinical Outcome Measures

Due to the severe phenotype of the h*Tau*$^{P301S/P301S}$ mice, from 5 months of age they underwent physical health checks twice a week. Upon noticing visible weight loss or abnormal gait (recorded as the time of first symptom), weighing commenced to establish a baseline followed by biweekly weighing. Once an animal lost 15% body weight, weighing was performed daily and diet was converted to a wet mash diet. Animals were culled and tissue collected as described in **Section 1.3.4** when they reached the 20% weight loss endpoint or if they developed full hind limb paralysis prior to the weight loss endpoint. Animal technicians observing symptom development were blinded to *Stx6* genotype.

Kaplan-Meier survival analysis followed by the Log-Rank test was performed to assess differences in time to first symptom and time to cull between *Stx6*$^{+/+}$;h*Tau*$^{P301S/P301S}$ and *Stx6*$^{-/-}$;h*Tau*$^{P301S/P301S}$ mice. These analyses were conducted with sexes combined given Mann-Whitney testing of sex differences revealed no differences.

##### 1.5.7 Tissue Collection

Tissues were collected as described in **Section 1.3.4** with spinal columns additionally being collected at 3 and 5 months. These were fixed in 10% (v/v) formal buffered saline and decalcified by rolling in 0.5M EDTA solution for 7 days prior to dissection into segments and processing into 3 paraffin blocks representing the cervical, thoracic and the lumbar region.

##### 1.5.8 Neuropathological Assessment

Brains were processed as described in **Section 1.3.6** with GFAP, Iba1 and synaptophysin staining being performed as previously described with the specific experimental conditions of novel stains detailed in **Table 18** with both sexes being assessed.

**Table 17. Overview of Experimental Conditions for Tau-Related Neuropathological Assessment.**

| Primary Antibody | Dilution | Secondary Antibody | Dilution | Pretreatment |
|---|---|---|---|---|
| NeuN | 1:3000 | Ab6720 | 1:200 | 8 min in superblock; ECC1 – CC1 at 95°C for 90 min |
| AT8 Biotin | 1:50 | N/A | N/A | 8 min in superblock; ECC1 – CC1 at 95°C for 90 min |
| MC1 | 1:100 | 1. Linker Rb vs Ms antibody<br>2. ab6720 | 1. 1:500<br>2. 1:200 | 8 min in superblock; CC1 at 95°C, 3 cycles at 32 min each i.e. 96 min |

NeuN-Positive Cell Density Image Analysis at 3 and 5 Months

NeuN-positive cell density was quantified in the superficial cortex as a previous study demonstrated neuronal loss in this region[21]. The cortex was manually annotated using NZConnect, with colour deconvolution subsequently being performed using QuPath. Then a machine learning-based pixel classifier was trained to distinguish tissue sections from glass and create whole tissue annotations. The region of interest (ROI) that was 500 µm deep from the cortical surface was obtained by first getting the ROI difference between the whole tissue annotation and the tissue annotation eroded by 500 µm. This band around the edge of the tissue section was then intersected with the cortex annotations to limit the ROI to within the cortex. Within the resulting annotation, Cellpose[22] was used to detect DAB-positive cells from the DAB deconvolved channel. Finally, an object classifier was applied to the resulting detections from Cellpose to filter out non-neuronal staining.

NeuN-positive cell density was calculated by:

$$\mathrm{NeuN}^{+}Cells/\mathrm{mm}^{2} = \frac{NeuN\ Cell\ Counts}{Area\ of\ Superficial\ Cortex} x\ 10^{6}$$

*Image Data Analysis*

Image data analysis was performed as described in **Section 1.3.6** with annotations being performed as follows:

Synaptophysin (5 Months): Images were annotated on QuPath focusing on regions that had been identified in the literature to exhibit robust synapse loss[23]. Specifically, a strip of the hippocampus, a strip of the cortex above the former, the dentate nucleus of the cerebellum and the brain stem were isolated. The thalamus was also annotated as an exploratory brain region. Stringent annotations were done avoiding white matter.

AT8 (3 & 5 Months): the cortex, hippocampus, thalamus, cerebellum, brain stem and superficial cortex (as described in NeuN image analysis) were annotated. As all brain regions at both 3 months and 5 months had a disease-associated increase, all regions were assessed for a modifying effect in *Stx6*$^{-/-}$;h*Tau*$^{P301S/P301S}$ mice. Statistics were done on $\log_{10}$ transformed data.

MC1 (3 & 5 Months): the cortex, hippocampus, thalamus, dentate nucleus of the cerebellum and brain stem were annotated. Only the thalamus, brain stem and dentate nucleus were assessed for a modifying effect in *Stx6*$^{-/-}$;h*Tau*$^{P301S/P301S}$ mice at 3 months as brain regions with a robust disease effect. All brain regions had a disease-associated increase in staining at 5 months so were assessed for rescue. Statistics were done on rank transformed data.

GFAP (5 Months): the cortex, hippocampus, thalamus, the dentate nucleus of the cerebellum and brain stem were annotated. The thalamus, brain stem and cortex showed a disease-related increase in GFAP staining so were prioritised to assess a modifying effect with syntaxin-6 knockout.

Iba1 (5 months): the cortex, hippocampus, thalamus, the dentate nucleus of the cerebellum and brain stem were annotated. Statistics were done on $Log_{10}$ transformed data. All brain regions examined showed a disease-associated increase in Iba1 staining so were assessed for a modifying effect with syntaxin-6 knockout.

AT8 Spines (3 and 5 months): Of the six AT8-stained thoracic sections of the spinal cord, the highest quality thoracic section was selected for quantification. One 5 month male *Stx6*$^{-/-}$;h*Tau*$^{P301S/P301S}$ mouse was excluded as the sections were at lumbar level. On Qupath, the whole area of the spinal cord as well as the grey matter were annotated. To extract the white matter area, the grey area was subtracted from the whole area. There was a disease-associated increase in staining in both the grey and white matter so both regions were assessed for a modifying effect with syntaxin-6 knockout.

##### 1.5.9 Biochemical Analysis

*Total Brain Homogenate Preparation*

20% (w/v) homogenates were prepared as described in **Section 1.3.7**. Following a 30 min incubation on ice, homogenates were distributed between a 20% (w/v) stock and a 10% (w/v) stock with 1x final Halt™ Protease and Phosphatase Inhibitor Cocktail. Aliquots of brain homogenates were prepared in prelubricated tubes to avoid multiple freeze-thaw cycles. Brain homogenate total protein was quantified using BCA assay as described in **Section 1.4.4**.

*Immunoblotting*

10% (w/v) homogenates with protease/phosphatase inhibitors (mixed sex) were clarified by centrifuging at 1,000 *g* for 5 min at 4°C with the supernatant being used for sample preparation. Immunoblotting was performed as described in **Section 1.4.4** with the exception that 30 µg total protein was loaded, electrophoresis was conducted at 150 V for 3 hr, Odyssey Blocking Buffer (TBS) was used for blocking and antibody dilution steps and TBS with 0.1% tween (TBST) was used for washing steps. IRDye 800CW Donkey anti-rabbit IgG was used at 1:5000 dilution and the IRDye 680RD goat anti-mouse IgG antibody at a 1:20,000 dilution. Phospho-tau species were detected first (AT8 or PHF1 at 1:500) following further immunodetection of total tau (K9JA, 1:5000) and β-actin (A5441, 1:5000).

Quantification was performed on Image Studio™ Software as described in **Section 1.4.4** with the exception that the total tau signal for each lane was first normalised to the LNF, which was subsequently used to correct the signal for the AT8/PHF1 tau signal. Finally, all values were normalised to an average of the *Stx6*$^{+/+}$;h*Tau*$^{P301S/P301S}$ animals.

##### 1.5.10 Biomarker Analysis

NfL was assessed in 3-, 4- and 5-month-old mice (mixed sex) as described in **Section 1.3.10.**

##### *1.5.11* Assessment of Seeding Differences Using Tau RD P301S FRET Biosensor Cells

Treatment of Tau RD P301S FRET Biosensor Cells with Mouse Brain Homogenates

Tau RD P301S FRET Biosensor cells were cultured in accordance with the general tissue culture practices described in **Section 1.4.2** using the media detailed in **Table 11.** Brain homogenates were prepared as described in **Section 1.3.7** as 10% (w/v) brain homogenates with protease and phosphatase inhibitors being subsequently clarified at 20,000 g for 15 min with the supernatant being transferred to single-use aliquots and total protein being quantified by BCA (see **Section 1.4.4**).

One day prior to treatment, Tau RD P301S FRET biosensor cells were seeded at 25,000 cells/well in 130 µL media into TPP 96-well plates. The following day, a dilution series of clarified 10% (w/v) brain homogenates were diluted in reduced-serum Opti-MEM. Diluted samples were subsequently mixed with an equal volume Opti-MEM supplemented with lipofectamine 2000 (1:8 dilution). Following a 20-30 min incubation at RT, 20 µL of the transfection mix was added to the cells *at* ~70% confluency, which were subsequently incubated at 37°C and 5% $CO_2$ within an IncuCyte S3 live cell imaging apparatus.

IncuCyte Imaging and Analysis of Aggregates formed in Tau Biosensor Cells

IncuCyte S3 Software (2022B/Rev2) acquired images using the 20× objective lens every 3 hr (16 images/well, same well coordinates) in phase contrast and green channels. Image analysis was performed using the Basic Analyzer module, employing a phase mask to quantify cell confluency and the green mask to quantify bright, punctate aggregates (FRET detected in GFP channel due to spectral overlap) as opposed to the baseline diffuse green background. Typical parameters for the green channel were as follows: segmentation - top-hat; radius- 10 µm; threshold- 4.75 GCU; edge split off.

#### 1.6 Key Resources Table

| Reagent or Resource | Source (Identifier) |
|---|---|
| **Antibodies & Conjugates [Clone] (Application)**<br>CL-WB: chemiluminescence western blot; F-WB: fluorescence western blot; FC: flow cytometry; SCA: scrapie cell assay; IHC: immunohistochemistry; IF: immunofluorescence. | |
| Anti-Syntaxin 6 Antibody [C34B2] (F-WB) | Cell Signalling Technologies (#2869) |
| Anti-PrP Antibody [ICSM18] (F-WB, FC, SCA) | D-Gen Ltd (PABC-107) |
| Anti-PrP Antibody [ICSM35] (IHC) | D-Gen Ltd |

| | |
|---|---|
| Anti-PrP Antibody [6D11] (IF, SCA, CL-WB) | Biolegend (808001) |
| Anti- PrP Antibody [5B2] (IF) | Santa Cruz (sc-47730) |
| Anti-β-Actin Antibody [AC-15] (F-WB) | Abcam (Ab6276) |
| Anti-β-Actin Antibody [AC-15] (F-WB) | Sigma (A5441) |
| Anti-β-Actin Antibody (F-WB) | Sigma (A2066) |
| Anti-Lamp1 Antibody [1D4B] (IF) | Santa Cruz Biotechnology (sc-19992) |
| Anti-Eea1 Antibody [C45B10] (IF) | Cell Signaling Technology (3288) |
| Anti-TGN46 Antibody (IF) | Abcam (ab16059) |
| Anti-Iba1 Antibody (IHC) | AlphaLaboratories (019-19741) |
| Anti-GFAP Antibody (IHC) | Dako (Agilent, Z0334) |
| Anti-Synaptophysin Antibody (IHC) | Invitrogen (PA5-27286) |
| AT8 Biotin Antibody (IHC) | Invitrogen (MN1020B) |
| Anti-NeuN Antibody (IHC) | Abcam (Ab177487) |
| Anti-Tau Antibody [K9JA] (F-WB) | DAKO (A0024) |
| Anti-Tau Antibody [MC1] (IHC) | Peter Davies (Described in Jicha *et al.* (1997)[24]) |
| Anti-pTau Antibody [PHF1] (F-WB) | Peter Davis (Described in Greenberg *et al.* 1992)[25]) |
| Anti-pTau Antibody [AT8] (F-WB, IHC) | Invitrogen (MN1020) |
| Biotinylated Polyclonal Rabbit Anti-Mouse Immunoglobulin Secondary Antibody (IHC) | Dako; Agilent (315-065-045-JIR) |
| Goat Anti-rabbit secondary antibody (IHC) | Abcam (Ab6720) |
| Alkaline Phosphatase conjugated anti-mouse IgG (CL-WB) | Sigma (A2179) |
| Alkaline Phosphatase anti-IgG (SCA) | Southern Biotechnology (1010-04) |

| | |
|---|---|
| IRDye 800CW donkey anti-rabbit (F-WB) | LI-COR (926-32213) |
| IRDye 680RD goat anti-mouse IgG (F-WB) | LI-COR (926-68070) |
| DAPI | Sigma (D8417-1MG) |
| Cross-adsorbed anti-mouse H+L-AF488 (IF) | Jackson (Stratech, 115-545-166-JIR) |
| Cross-adsorbed anti-mouse H+L-Rhodamine X (IF) | Jackson (Stratech, 115-295-166-JIR) |
| Cross-adsorbed anti-mouse IgG1-AF488. (IF) | Jackson (Stratech, 115-545-205-JIR) |
| Cross-adsorbed anti-mouse IgG2a-Rhodamine X (IF) | Jackson (Stratech, 115-295-206-JIR) |
| Cross-adsorbed anti-mouse IgG2b Rhodamine X (IF) | Jackson (Stratech, 115-295-207) |
| Cross-adsorbed anti-mouse IgG1 AlexaFluor647 (FC) | Jackson (Stratech, 115-505-205-JIR) |
| Cross-adsorbed anti-rabbit H+L-AF488 (IF) | Jackson (Stratech, 111-545-144-JIR) |
| Cross-adsorbed anti-rabbit IgG TRITC (IF) | Jackson (Stratech, 711-025-152-JIR) |
| Cross-adsorbed anti-rat IgG Rhodamine (IF) | Jackson (Stratech, 712-295-150-JIR) |
| **Experimental Models: Cell Lines** | |
| PK1/2 Cell Line (Derivative of Mouse Neuro2a Cells, ATCC, CCL-131) | Laboratory of Peter Klöhn (Described in Klöhn *et al.* (2003)[8]) |
| S7 (Derivative of N2aPK1 Cells) | Laboratory of Peter Klöhn (Described in Marbiah *et al.* (2014)[15]) |
| iS7 Cells (Persistently Infected S7 Cells) | Laboratory of Peter Klöhn (Described in Ribes *et al.* (2023)[14]) |
| CAD-2A2D5 (CAD5) Cells (Derivative from Cath.a-differentiated (CAD) Cells Described in Qi *et al.* (1997)[26] | Laboratory of Charles Weissman (Described in Mahal *et al.* (2014)[27]) |
| Tau RD P301S FRET Biosensor Cell Line | ATCC (CRL-3275) |
| **Experimental Models: Mouse Lines** | |

| | |
|---|---|
| C57BL/6NTac-Stx6$^{em1(IMPC)H}$ | UKRI-MRC Harwell as part of the IMPC (Allele MGI ID: 6266827; EM:12503) |
| Thy1-hTau.P301S (CBA.C57BL/6) | Laboratory of Michel Goedert (Described in Allen *et al.* (2002)[18] |
| **Biological Samples** | |
| RML-Infected CD1 Brain Homogenate (Cell Infection) | Laboratory of Jonathan Wadsworth (I8700) |
| 22L-Infected C57BL/6 Brain Homogenate (Cell Infection) | Laboratory of Jonathan Wadsworth (I23185) |
| MRC2-Infected SJL Brain Homogenate (Cell Infection) | Laboratory of Jonathan Wadsworth (I17800) |
| ME7-Infected C57BL/6 Brain Homogenate (Cell Infection) | Laboratory of Jonathan Wadsworth (I21487) |
| Exosome Fraction from Infected Cells (Cell Infection) | Laboratory of Peter Klöhn (Described in Ribes *et al.* (2023)[14]) |
| RML-Infected C57BL/6 Brain Homogenate (Mouse Infection) | Laboratory of Jonathan Wadsworth (I21488) |
| Normal C57BL/6 Brain Homogenate (Diluent for Mouse Infection) | Laboratory of Jonathan Wadsworth (14052) |
| **Recombinant DNA** | |
| pGFP-V-RS Vectors containing Stx6-Targeting 29-mer shRNA Sequence or Non-Silencing Control | Insight Biotechnology (TG517176A, TG517176B, TG517176C, TG517176D, TR30013) |
| pRetroSuper vector containing Prnp-targeting shRNA (Si8) | Laboratory of Parmjit Jat (Originally Described in Goold et al. (2011)[29]) |
| pRP[Exp]-mCherry-CAG>hyPBase | VectorBuilder (VB010000-9365tax) |
| pPB[Exp]-EGFP/Puro-mPGK>mStx6[NM_021433.3] | VectorBuilder (VB220821-1010eff) |
| pPB[Exp]-EGFP/Puro-mPGK>ORF_Stuffer | VectorBuilder (VB220828-1137yrp) |
| pPB[shRNA]-EGFP:T2A:Puro-U6>(mStx6[shRNA#4]) | VectorBuilder (VB221229-1281dfp) |

| | |
|---|---|
| pPB[shRNA]-EGFP:T2A:Puro-U6>(NSC[shRNA]) | VectorBuilder (VB230103-1034jjw) |
| pPB[shRNA]-EGFP:T2A:Puro-U6>(Prnp[shRNA]) | VectorBuilder (VB230103-1039ctx) |
| **Cell Culture Reagents and Consumables (Identity)** | |
| Opti-MEM (1X) Reduced Serum Medium | Gibco (31985-047) |
| DMEM (1X) + GlutaMAX-I | Gibco (31966-0211) |
| Fetal Bovine Serum, Heat Inactivated, Qualified, HI | Gibco (10500064) |
| Bovine Growth Serum Supplemented Calf | HyClone (SH30541.03) |
| Penicillin Streptomycin (Pen Strep) | Life Technologies (15140-122) |
| 0.05% Trypsin-EDTA 91X) | Gibco (25300-054) |
| Puromycin dihydrochloride | Sigma (P8833) |
| Dimethyl sulfoxide | Sigma (D8418-100ML) |
| Cryovial, 2.0ml, Sterile | GREINER BIO-ONE LTD (122278) |
| 25 mL Disposable Reagent Reservoir | Thermoscientific (8094) |
| Cytofix/Cytoperm™ Fixation/Permeabilization Kit | BD Biosciences (554714) |
| CellTiter-Glo® Reagent | Promega (G7570) |
| Greiner CELLSTAR® 96 well plates (for CellTiter-Glo ) | Greiner (655088) |
| TPP 96-well Plates | TPP (Z707910) |
| Poly-L-lysine | Sigma (P1274) |
| Distilled Water | Gibco (1523-147) |
| Nunc™ Lab-Tek™ Chambered Coverglass | Thermoscientific (155411) |
| Formaldehyde | Sigma (T9284) |
| Acetone | ThermoScientific (176800025) |

| | |
|---|---|
| Superblock Protein in 2.5 mM Tris, 0.15 M NaCl, pH 7.4 | Thermo Scientific (0037545) |
| Immersol immersion oil 518 F | Carl Zeiss (10539438) |
| Lipofectamine™ LTX Reagent with PLUS™ Reagent | Thermo Fisher (A12621) |
| Lipofectamine™ 2000 Transfection Reagent | Thermo Fisher (11668030) |
| Countess™ Cell Counting Chamber Slides | Invitrogen (C10228) |
| C-CHIP Disposable Hemocytometer Neubauer Improved | NanoEntek (DHC-N01-50) |
| Trypan Blue Stain 0.4% | Invitrogen (T10282) |
| Cycloheximide solution | Sigma (C4859) |
| 96 well plate | Greiner (650101) |
| 0.2 µm PTFE Membrane Filter | Fisher Scientific (156608) |
| Biosphere SafeSeal Tube 1.5 mL | Sarstedt (72.706.200) |
| 24 Well Cell Culture Cluster Flat Bottom with Lid | Costar (3524) |
| Nunclon Delta Surface 6-well dish | Fisher Scientific (140675) |
| Easy flask 25 filt nunclon d silicon | Thermoscientific (156367) |
| Easyflask 75 filt nunclon d silicon | Thermoscientific (156499) |
| Nunclon Delta Surface 10 cm dish | Thermoscientific (172931) |
| Mega-8 | AdipoGen Life Sciences (AG-CC1-0007-G005) |
| Triton X-114 | Sigma (9036-19-5) |
| Cyclodextrin | Cayman Chemical Company (21633) |
| **Biochemistry Reagents (Protein)** | |
| RIPA Buffer | Sigma (R0278) |

| | |
|---|---|
| Halt™ Protease Inhibitor Cocktail, EDTA-free (100X) | ThermoFisher Scientific (87785) |
| Halt Protease & Phosphatase Inhibitor Cocktail (100X) | ThermoScientific (78444) |
| Benzonase | EMD Millipore (70664) |
| Zirconium Ceramic Oxide Bulk Beads (1.4mm) | Fisherbrand (15515809) |
| Albumin Standard | Thermo Scientific (23209) |
| Pierce BCA Protein Assay Kit | Thermo Scientific (23227) |
| Microplate, 96-well, PS, F bottom, clear | Greiner Bio-One (655101) |
| 4x Laemelli Sample Buffer | Biorad (1610747) |
| NuPAGE™ LDS Sample Buffer 4X | Thermo Scientific (NP0007) |
| NuPAGE™ Sample Reducing Agent 10X | Thermo Scientific (NP0004) |
| NuPage™ MOPS Running Buffer 20X | Thermo Scientific (NP0001) |
| 2-Mercaptoethanol | Sigma (M3148) |
| 20x NuPAGE MES SDS running buffer | Novex (NP002) |
| SeeBlue Pre-stained Protein Standard | Thermo Fisher Scientific (LC5625) |
| NuPAGE™ 4 to 12%, Bis-Tris, 1.5 mm Gel, 10-well | Invitrogen (NP0335BOX) |
| NuPAGE™ 4 to 12%, Bis-Tris, 1.5 mm Gel, 15-well | Invitrogen (NP0336BOX) |
| NuPage™ 12% Bis-Tris Protein Gels (12% bis-tris gel) | Thermo Scientific (NP0341BOX) |
| Low Binding Lubricated Microcentrifuge Tubes 0.5 mL | Costar (3206) |
| Low Binding Lubricated Microcentrifuge Tubes 1.5 mL | Costar (3207) |
| Methanol | Fisher UK (M/4000/17) |
| 10X tris-glycine buffer | National Diagnostics (EC-880) |

| | |
|---|---|
| Nitrocellulose Membrane Filter Paper Sandwich | Themo Scientific (LC2001) |
| Immobilon-P PVDF | Millipore (IPVH00010) |
| Whatman Electroblotting Filter Paper | Whatman (3030 917) |
| Intercept Blocking Buffer PBS | LI-COR (927-70001) |
| Intercept Blocking Buffer TBS | LI-COR (927-60001) |
| Skimmed Milk Power | Sigma (70166) |
| Revert 700 Total Protein Stain Kit | LI-COR (926-11010) |
| Tween-20 | Sigma (P9416) |
| Phosphate Buffered Saline 10X | VWR (437117K) |
| TRIS-buffered saline (TBS, 10X) pH 7.4, for Western blot | Thermofisher (J62938.K2) |
| Amersham Hyperfilm MP | GE Healthcare (28906842) |
| Trizma Base | Sigma (T6066) |
| $MgCl_2$ | Sigma (M8266) |
| CDP-Star™ Chemiluminescent Substrate (CDP-Star) | Thermo Scientific (T2147) |
| D-PBS (1X) | Gibco (14190-094) |
| ELISpot IP Filter Plates (PVDF membrane, 0.45 µm) | Merck (MSIPN4550) |
| Proteinase K | Roche (3115828001) |
| Pronase (Protease from *Streptomyces griseus*) | Sigma (P5147) |
| Guanidinium Thiocyanate | Generon (GDR0244) |
| Superblock Dry Blend (TBS) Blocking Buffer | Pierce/Fisher (10426144) |
| TritonX-100 | SLS (T9284) |

| | |
|---|---|
| Trizma(R) Hydrochloride | SLS (T6666) |
| Deoxycholic Acid, Na Salt | Sigma (D6750) |
| AP Conjugate Substrate Kit | BIO-RAD (170-6432) |
| Haematoxylin | Sigma (05174.0500) |
| NF-light V2 Advantage Kit | Quanterix (104073) |
| **Biochemistry Work (RNA)** | |
| RNase Zap Wipes | Invitrogen (AM9788) |
| Qubit Assay Tubes | Invitrogen (Q32856) |
| Qubit RNA BR Reagent | Invitrogen (Q10211A) |
| Qubit RNA BR Buffer | Invitrogen (Q10211) |
| QuantiTect Primer Assay Mm_Actb_1_SG | Qiagen (QT00095242) |
| QuantiTect Primer Assay Mm_Prnp_1_SG | Qiagen (QT00101080) |
| Fast SYBR™ Green Master Mix | Thermofisher (4385612) |
| Quantitect Reverse Transcription Kit | Qiagen (205311) |
| MicroAmp Fast Optical 96-Well Reaction Plate with Barcode (0.1 mL) | Applied Biosystems (4346906) |
| RNase-Free 1.5 mL Microfuge Tubes | Ambion (AM12400) |
| TRI Reagent | Zymo Research (R2050-1-200) |
| Direct-zol RNA Miniprep | Zymo Research (R2052) |
| RNA Clean & Concentrator 6 kit | Zymogen (R1014) |
| 2.0 mm Bashing Bead Lysis tubes | Zymogen (S6003-50) |
| Universal Plus Total RNA-Seq with Mouse AnyDeplete kit | Tecan (9157-A01) |
| DNA/RNA Shield | Zymogen (R1100-50) |

| | |
|---|---|
| High Sensitivity RNA ScreenTape | Agilent (5067-5579) |
| High Sensitivity RNA ScreenTape Sample Buffer | Agilent (5067-5580) |
| D1000 ScreenTape | Agilent (5067-5582) |
| D1000 reagents | Agilent (5067-5583) |
| NextSeq 2000 Kit P2 100 cycles | Illumina |
| Elution buffer | Qiagen (19086) |
| PhiX sequencing control library | Illumina (FC-110-3002) |
| Optical Cap, 8x Strip | Agilent Technologies (401425) |
| Tubes, 0.2 mL, Thin Wall 8 Strips | Agilent Technologies (410092) |
| Loading Tips | Agilent Technologies (5067-5153) |
| **Biochemistry Work (DNA)** | |
| 0.2ml Skirted 96-well PCR Plate | Thermo Scientific (AB-0800) |
| Adhesive Sealing Sheets | Thermo Scientific (AB-0558) |
| MyTaq Extract-PCR Kit | Meridian Bioscience (BIO-21126) |
| MyTaq HS Red Mix | Meridian Bioscience (BIO25047) |
| UltraPure™ Agarose | Invitrogen (16500500) |
| UltraPure™ TBE Buffer, 10X | Invitrogen (15581044) |
| TAE Buffer | National Diagnostics (EC-872) |
| PAGE GelRed® Nucleic Acid Gel Stain | Biotium (41008) |
| Ambion Nuclease-Free Water | Invitrogen (AM9906) |
| TaqPath™ ProAmp™ Master Mix | ThermoFisher Scientific (A30865) |
| **Immunohistochemical Reagents** | |

| | |
|---|---|
| LEICA SCN400F scanner | LEICA Milton Keynes, UK |
| NanoZoomer 360 | C13220-01, Hamamatsu |
| DAB Map Detection Kit | Roche Tissue Diagnostics (05266360001, 760-124) |
| **Resources Related to the Animal Facility** | |
| Tecniplast Blueline IVC (for stock) | Tecniplast (1145) |
| Tecniplast Blueline IVS (for breeders) | Tecniplast (1284) |
| SARSTEDT Microvette® Serum tubes | Sarstedt (20.1308) |
| “Nestlet” | Dates and Ltd., Manchester, UK (CS3A01) |
| Rotarod for Mice | Ugo Basile Biological Research (Model 47600, V04) |
| **Equipment (Identity)** | |
| Countess III FL | Invitrogen (16832556) |
| Infinite F200 PRO | Tecan |
| Odyssey® Imaging System | LI-COR (Model 9120). |
| Hybaid PS 250 Electrophoresis Power Supply | Hybaid (176435049995) |
| Beckman Coulter CytoFLEX | Beckman Coulter |
| Incucyte S3 Live Cell Imaging Apparatus | Sartorius (4647) |
| Qubit 2.0 Fluorometer 2.0 | Invitrogen (Q32866) |
| Agilent Technologies 2200 TapeStation | Agilent (G2964AA) |
| NanoDrop ND-1000 Spectrometer | Thermo Scientific |
| Universal Hood II Gel Doc System | Bio-rad (721BR1519) |
| Ribolyser Precellys 24 | Berin Technologies |
| Biomek FX liquid handling robot | Beckman |

| | |
|---|---|
| Bioreader 5000-Eβ | BioSys Karben, Germany |
| XCell SureLock | Novex (EI0001) |
| Ventana Discovery XT Autostainer | Roche Tissue Diagnostics (315724) |
| Gemini AS Automated Slide Stainer | Epredia (15243973) |
| Simoa HD-X platform | Quanterix (QTX-103385) |
| **Software** | |
| ImageLab 6.0 | Biorad (https://www.bio-rad.com/en-uk/product/image-lab-software) |
| NDP.serve3 | NanoZoomer Digital Pathology (https://nanozoomer.unisa.edu.au/ndp/serve/home) |
| NZConnect 1.0.36 (IVD) | Hamamatsu (https://nzconnect.ion.ucl.ac.uk/nz/connect/home) |
| Qupath (Version 0.4.3) | Bankhead et al.[30] (https://qupath.github.io/) |
| i-control (Version 3.4.2.0) | Tecan |
| Odyssey Acquisition Software (Version 2.1) | LI-COR (https://www.licor.com/bio/las/) |
| Image Studio Lite Ver 5.2 | LI-COR (discontinued) |
| ZEN 2.3 SP1 (Black Edition) | Zeiss (https://www.micro-shop.zeiss.com/en/us/softwarefinder/software-categories/zen-black/zen-black-system/) |
| ZEN 2.5 (Blue Edition) | Zeiss (https://www.micro-shop.zeiss.com/en/de/softwarefinder/software-categories/zen-blue/) |
| ImageJ 1.53n | National Institutes of Health, USA |
| InVivoStat v4.7.0 | Clark et al. (2012)[31] |
| GraphPad Prism 9 | GraphPad (https://www.graphpad.com/features) |

| | |
|---|---|
| Incucyte 2022B Rev2 | Satorius (https://downloads.essenbioscience.com/get/incucyte-2022b-rev2-gui) |
| Tapestation Analysis Software A.02.02 (SP1) | Agilent |
| Adobe Illustrator 2024 | Adobe (https://www.adobe.com/uk/products/illustrator/free-trial-download.html) |

#### References


1 Jones, E. *et al.* Characterisation and prion transmission study in mice with genetic reduction of sporadic Creutzfeldt-Jakob disease risk gene Stx6. *Neurobiology of Disease* **190**, 106363 (2024). https://doi.org:https://doi.org/10.1016/j.nbd.2023.106363

2 Rangel, A. *et al.* Distinct patterns of spread of prion infection in brains of mice expressing anchorless or anchored forms of prion protein. *Acta Neuropathol Commun* **2**, 8 (2014). https://doi.org:10.1186/2051-5960-2-8

3 O'Shea, M. *et al.* Investigation of mcp1 as a quantitative trait gene for prion disease incubation time in mouse. *Genetics* **180**, 559-566 (2008). https://doi.org:10.1534/genetics.108.090894

4 Wadsworth, J. D. *et al.* Human prion protein with valine 129 prevents expression of variant CJD phenotype. *Science (New York, N.Y.)* **306**, 1793-1796 (2004). https://doi.org:10.1126/science.1103932

5 Hamilton, M. A., Russo, R. C. & Thurston, R. V. Trimmed Spearman-Karber method for estimating median lethal concentrations in toxicity bioassays. *Environmental Science & Technology* **11**, 714-719 (1977). https://doi.org:10.1021/es60130a004

6 Jones, E. *et al.* Knockout of Sporadic Creutzfeldt-Jakob Disease Risk Gene in Mice Extends Prion Disease Incubation Time. *bioRxiv*, 2023.2001.2010.523281 (2023). https://doi.org:10.1101/2023.01.10.523281

7 Sandberg, M. K. *et al.* Prion neuropathology follows the accumulation of alternate prion protein isoforms after infective titre has peaked. *Nature Communications* **5**, 4347 (2014). https://doi.org:10.1038/ncomms5347

8 Klöhn, P. C., Stoltze, L., Flechsig, E., Enari, M. & Weissmann, C. A quantitative, highly sensitive cell-based infectivity assay for mouse scrapie prions. *Proceedings of the National Academy of Sciences of the United States of America* **100**, 11666-11671 (2003). https://doi.org:10.1073/pnas.1834432100

9 R: A Language and Environment for Statistical Computing (2022).

10 Gentleman, R. C. *et al.* Bioconductor: open software development for computational biology and bioinformatics. *Genome Biology* **5**, R80 (2004). https://doi.org:10.1186/gb-2004-5-10-r80

11 Anders, S. & Huber, W. Differential expression analysis for sequence count data. *Genome Biology* **11**, R106 (2010). https://doi.org:10.1186/gb-2010-11-10-r106

12 Love, M. I., Huber, W. & Anders, S. Moderated estimation of fold change and dispersion for RNA-seq data with DESeq2. *Genome Biology* **15**, 550 (2014). https://doi.org:10.1186/s13059-014-0550-8
13 Varet, H., Brillet-Guéguen, L., Coppée, J.-Y. & Dillies, M.-A. SARTools: A DESeq2- and EdgeR-Based R Pipeline for Comprehensive Differential Analysis of RNA-Seq Data. *PloS one* **11**, e0157022 (2016). https://doi.org:10.1371/journal.pone.0157022
14 Ribes, J. M. *et al.* Prion protein conversion at two distinct cellular sites precedes fibrillisation. *Nature Communications* **14**, 8354 (2023). https://doi.org:10.1038/s41467-023-43961-1
15 Marbiah, M. M. *et al.* Identification of a gene regulatory network associated with prion replication. *The EMBO journal* **33**, 1527-1547 (2014). https://doi.org:10.15252/embj.201387150
16 Costes, S. V. *et al.* Automatic and Quantitative Measurement of Protein-Protein Colocalization in Live Cells. *Biophys J* **86**, 3993-4003 (2004). https://doi.org:https://doi.org/10.1529/biophysj.103.038422
17 Hannus, M. *et al.* siPools: highly complex but accurately defined siRNA pools eliminate off-target effects. *Nucleic Acids Res* **42**, 8049-8061 (2014). https://doi.org:10.1093/nar/gku480
18 Allen, B. *et al.* Abundant tau filaments and nonapoptotic neurodegeneration in transgenic mice expressing human P301S tau protein. *The Journal of neuroscience : the official journal of the Society for Neuroscience* **22**, 9340-9351 (2002). https://doi.org:10.1523/JNEUROSCI.22-21-09340.2002
19 Scattoni, M. L. *et al.* Early behavioural markers of disease in P301S tau transgenic mice. *Behav Brain Res* **208**, 250-257 (2010). https://doi.org:10.1016/j.bbr.2009.12.002
20 Sukoff Rizzo, S. J. *et al.* Assessing Healthspan and Lifespan Measures in Aging Mice: Optimization of Testing Protocols, Replicability, and Rater Reliability. *Current Protocols in Mouse Biology* **8**, e45 (2018). https://doi.org:https://doi.org/10.1002/cpmo.45
21 Hampton, D. W. *et al.* Cell-mediated neuroprotection in a mouse model of human tauopathy. *The Journal of neuroscience : the official journal of the Society for Neuroscience* **30**, 9973-9983 (2010). https://doi.org:10.1523/jneurosci.0834-10.2010
22 Stringer, C., Wang, T., Michaelos, M. & Pachitariu, M. Cellpose: a generalist algorithm for cellular segmentation. *Nature methods* **18**, 100-106 (2021). https://doi.org:10.1038/s41592-020-01018-x
23 Vogler, L. *et al.* Assessment of synaptic loss in mouse models of β-amyloid and tau pathology using [(18)F]UCB-H PET imaging. *Neuroimage Clin* **39**, 103484 (2023). https://doi.org:10.1016/j.nicl.2023.103484
24 Jicha, G. A., Bowser, R., Kazam, I. G. & Davies, P. Alz-50 and MC-1, a new monoclonal antibody raised to paired helical filaments, recognize conformational epitopes on recombinant tau. *Journal of neuroscience research* **48**, 128-132 (1997). https://doi.org:10.1002/(sici)1097-4547(19970415)48:2<128::aid-jnr5>3.0.co;2-e
25 Greenberg, S. G., Davies, P., Schein, J. D. & Binder, L. I. Hydrofluoric acid-treated tau PHF proteins display the same biochemical properties as normal tau. *The Journal of biological chemistry* **267**, 564-569 (1992).
26 Qi, Y., Wang, J. K., McMillian, M. & Chikaraishi, D. M. Characterization of a CNS cell line, CAD, in which morphological differentiation is initiated by serum

deprivation. *The Journal of neuroscience : the official journal of the Society for Neuroscience* **17**, 1217-1225 (1997). https://doi.org:10.1523/jneurosci.17-04-01217.1997
27 Mahal, S. P. *et al.* Prion strain discrimination in cell culture: the cell panel assay. *Proceedings of the National Academy of Sciences of the United States of America* **104**, 20908-20913 (2007). https://doi.org:10.1073/pnas.0710054104
28 McEwan, W. A. *et al.* Cytosolic Fc receptor TRIM21 inhibits seeded tau aggregation. *Proceedings of the National Academy of Sciences of the United States of America* **114**, 574-579 (2017). https://doi.org:10.1073/pnas.1607215114
29 Goold, R. *et al.* Rapid cell-surface prion protein conversion revealed using a novel cell system. *Nature Communications* **2**, 281 (2011). https://doi.org:10.1038/ncomms1282
30 Bankhead, P. *et al.* QuPath: Open source software for digital pathology image analysis. *Scientific Reports* **7**, 16878 (2017). https://doi.org:10.1038/s41598-017-17204-5
31 Clark, R. A., Shoaib, M., Hewitt, K. N., Stanford, S. C. & Bate, S. T. A comparison of InVivoStat with other statistical software packages for analysis of data generated from animal experiments. *J Psychopharmacol* **26**, 1136-1142 (2012). https://doi.org:10.1177/0269881111420313
